# New avenues for human blood plasma biomarker discovery via improved in-depth analysis of the low-abundant *N-*glycoproteome

**DOI:** 10.1101/2023.10.22.562384

**Authors:** Frania J. Zuniga-Banuelos, Marcus Hoffmann, Udo Reichl, Erdmann Rapp

## Abstract

To understand implications of protein glycosylation for clinical diagnostic and biopharmaceuticals, innovative glycoproteomic technologies are required. Recently, significant advances were made, particularly toward structure-focused *N-*glyco-proteomic analyses. The mass spectrometric analysis of intact *N-*glycopeptides using stepped collision fragmentation along with glycan oxonium ion profiling now enables to reliably discriminate between different *N-*glycan structures. Still, there are weaknesses that current *N-*glycoproteomic approaches must overcome: 1) handling of incorrect identifications, 2) identification of rare and modified *N-*glycans, and 3) insufficient glycoproteomic coverage, especially in complex samples. To address these shortcomings, we have developed an innovative *N-*glycoproteomic workflow that aims at providing comprehensive site-specific and structural *N-*glycoproteomic data on human blood plasma glycoproteins. The workflow features protein depletion plus various fractionation strategies and the use of high-resolution mass spectrometry with stepped collision fragmentation. Furthermore, by including a decision tree procedure established for data validation, we could significantly improve the description of the *N-*glycan micro-heterogeneity. Our data analysis workflow allows the reliable differentiation of ambiguous *N-*glycan structures like antenna-versus core-fucosylation plus the modified and rare *N-*glycans such as sulfated and glucuronidated ones. With this workflow, we were able to advance in the analysis of human blood plasma glycoproteins to concentrations as low as 10 pg/mL. A total of 1,929 *N-*glycopeptides and 942 *N-*glycosites derived from 805 human middle-to low-abundant glycoproteins were identified. Overall, the presented workflow holds great potential to improve our understanding of protein glycosylation and to foster the discovery of blood plasma biomarkers.

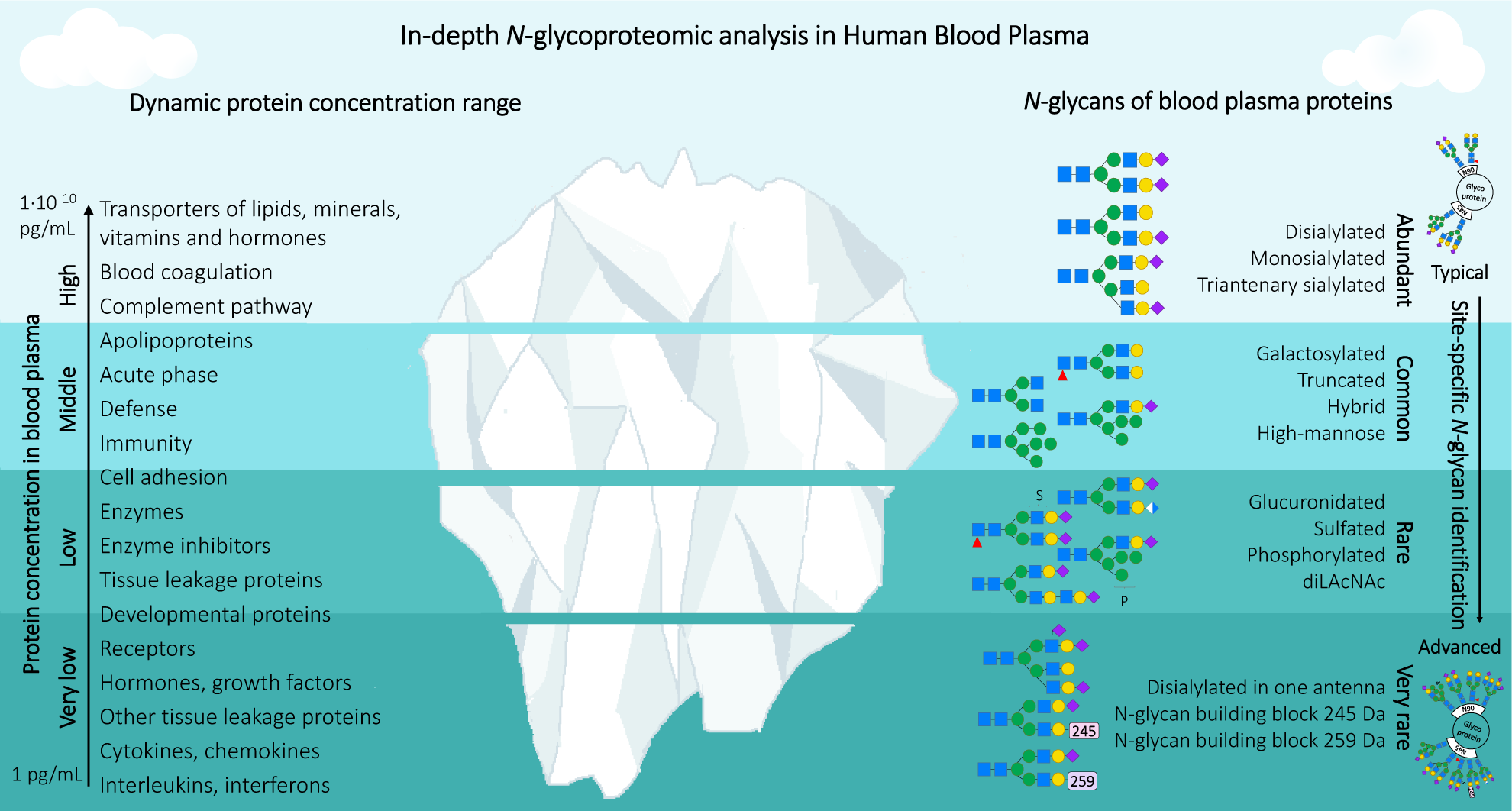

## 1. Introduction

Approximately 60% of the proteins reported as secreted to blood by Protein Atlas [1,2], are described as potentially glycosylated in UniProtKB database [3,4]. A number of clinical studies have revealed alterations in the *N-*glycosylation of human blood plasma proteins due to physiological and pathophysiological changes [5–7]. Glycoproteomic analysis via liquid chromatography coupled to tandem mass spectrometry (LC-MS/MS) is the core technology for detecting such alterations of *N-*glycosylation in blood plasma proteins in a site-specific manner [8]. In recent years, explorative large-scale glycopeptide-centered *N-*glycoproteomic studies on human plasma proteins, like IgG, enabled the identification of glycoforms that could serve as potential biomarkers or therapeutic targets [5,9–11]. Promising blood plasma biomarker candidates are tissue-leakage proteins or signaling molecules due to their organ specificity [9]. Nevertheless, limitations related to the analysis of blood plasma and MS-based *N-*glycopeptide identification itself still impede the full diagnostic potential of *N-*glycoproteomic analyses. Firstly, the average concentration of tissue-leakage proteins in blood plasma is 1·105–1·10^7^ times lower than the albumin concentration (35–50 mg/mL), the major blood plasma protein [12]. Due to the challenging concentration range, established workflows for blood plasma (glyco)proteomics frequently include both, a high-abundant protein (HAP) depletion step and a fractionation step either on the peptide or protein level [13,14]. A second constrain is that current MS-based workflows for intact *N-*glycopeptide identification give only a limited description of *N-*glycan structural information and an incomplete deduction of the input spectra. An example of the effect of these limitations is glucuronidated *N-*glycans, which were detected in blood plasma through glycomic experiments, but not reported yet through *N-*glycoproteomic analysis [15,16]. This results in differences between glycoforms identified by glycomics and glycoproteomics analysis, and points out the necessity to improve MS-based workflows for accurately describing the *N-*glycoproteome [17].

To conduct an optimal *N-*glycoproteomic search on a given LC-MS/MS dataset, the input search parameters (*N-*glycan list, protein list, modifications, etc.) must describe the characteristics of the different *N-*glycopeptides present in the sample. Furthermore, factors that neglect the existence of rare *N-*glycans (such as terminal glucuronic acid) and ignore post-glycosylational *N-*glycan modifications such as sulfation, phosphorylation, and acetylation can prevent the precise description of the *N-*glycoproteome. To date, *N-*glycoproteomic data analyses tend to yield variable *N-*glycan and glycoprotein identifications when different *N-*glycopeptide search strategies and bioinformatics tools are applied and compared [18,19]. The study from Kawahara *et al.* proposes that the main bottlenecks for obtaining precise results are the scoring algorithms and validation filters for glycopeptide spectra match (gPSM) [18]. Even though advanced false discovery rate (FDR) validation methods might improve the quality of gPSM, only a few software programs have demonstrated more efficient FDR validation methods, after a controlled search [20,21]. Thus, during an in-depth *N-*glycoproteomic data analysis, manual validation is still necessary for finding and reprocessing missed matches, as was also concluded by the study from Lee *et al*., 2016 [22].

Defining the *N-*glycan structure is another desired but underdeveloped aspect from the *N-*glycoproteomic analysis. The fragmentation of *N-*glycopeptides is favored by applying higher-energy collisional dissociation (HCD) based methods, while for *O-*glycopeptides, electron-transfer/higher-energy collision dissociation (EThcD) based methods are more useful [23]. In addition, diverse combinations of collisional energies applied during fragmentation spectra acquisitions reflect specific advantages towards the description of a glycopeptide [24,25]. A prominent set up is the HCD spectra acquired with a set of increasing normalized collisional energies (NCE), e.g. 20, 35, and 50 [26]. This HCD fragmentation with stepped energy (HCD.step) is more efficient since it simultaneously generates information corresponding to both the peptide and *N-*glycan composition [26,27]. In comparison, HCD spectra acquired at low NCE (HCD.low, e.g. fixed NCE 20) are rich in *N-*glycan structural information [26,28]. The structural B and Y ions support the differentiation of isomeric *N-*glycan structures at high confidence, as recently demonstrated by Shen *et al*. [29]. Therefore, via analyzing both spectra, the differentiation of isomeric *N-*glycan structures containing antenna or core fucosylation, bisecting HexNAc and diLAcNAc becomes possible. Furthermore, it is possible to collect more evidence regarding phosphorylation and sulfation modifications, critical for understanding glycoprotein-receptor interactions [30–36]. Nonetheless, the detection of new *N-*glycan compositions and structures by discovering unexpected ions corresponding to rare *N-*glycan building blocks (e.g. glucuronic acid) is an explorative task, which can only be accomplished by manual annotation.

Here we developed both, a sample preparation and a data analysis workflow that allows in*-*depth *N-*glycoproteomic analysis of human blood plasma. The preparative workflow not only enables the detection of glycoproteins within a concentration range of 10 orders of magnitude, but also better characterizes the micro-heterogeneity of their *N-*glycans. The intact *N-*glycopeptide LC-MS/MS-based analysis, established here, includes the acquisition of spectra using HCD.step and HCD.low fragmentation energies, which supports the differentiation of isomeric *N-*glycan structures. In order to conduct an in-depth exploration, no FDR cuts nor decoys were applied during the glycoproteomic search, and manual validation was performed instead. Manual validation of the *N-*glycopeptides was based on a decision tree established as part of the data analysis workflow. After the validation, we achieved the detection of 1929 *N-*glycopeptides, 942 *N-*glycosites, and 805 glycoproteins. Correction of the invalid or partially valid *N-*glycopeptide identifications by *de novo N-*glycan sequencing, disclosed rare *N-*glycan building blocks (like glucuronic acid). Explorative data analysis also allowed the reliable identification of *N-*glycan modifications such as sulfation, phosphorylation, and atypical branching structures with the linkage HexNAc-NeuAc. Overall, the application of the developed workflows enabled a high level of structural *N-*glycan information and the site-specific *N-*glycoproteomic analysis of even very low-abundant human blood plasma glycoproteins with high confidence (e.g., cysteine-rich secretory protein-3 6.31 pg/mL). The gained insights and improvements will support further glycoproteomic software development and are crucial for future *N-*glycoproteomic studies aiming at the identification of therapeutic targets or biomarkers in human blood plasma.

## 2. Materials and Methods

All chemicals used were LC-MS grade or of the highest purity available. Milli-Q water was used for preparing all aqueous solutions (18 MΩ×cm, <5 ppb, Millipore Milli-Q® Reference A+ system, Merck Millipore, Germany). LC-MS grade acetonitrile (ACN, #A955-212) was purchased from Fisher Scientific (Schwerte, Germany). 2,2,2-trifluoroethanol (TFE, #808259), potassium chloride (KCl, #104935), potassium phosphate dibasic (K_2_HPO_4_, #104873), and sodium phosphate dibasic (Na_2_HPO4, #106585) was purchased from Merck (Darmstadt, Germany). Sodium chloride (NaCl, #P029.3) was purchased from Carl Roth (Karlsruhe, Germany). Trifluoroacetic acid (TFA, #28904) was purchased from Fisher Scientific. Ammonium bicarbonate (ABC, #09830) was purchased from Merck. Formic acid (FA, #56302), DL-dithiothreitol (DTT, #D5545), iodoacetamide (IAA, #I1149), calcium chloride (CaCl_2_, #A4689), and sodium dodecyl sulfate (SDS, #75746) were purchased from Merck. Sequencing grade, modified trypsin was purchased from Promega (#V5111, Madison, WI, USA).

### 2.1. Sample preparation

The human blood plasma sample (HBP) was acquired from Affinity Biologicals (VisuCon^TM^-F Frozen normal control plasma, FRNCP0105, Ancaster, ON, Canada). The steps applied in the sample preparation workflow are depicted in **Figure 1**. The top 14 high-abundant blood plasma proteins (HAP) were depleted using single-use High Select™ Top 14 Abundant Protein Depletion Midi Spin Columns (#A36371, Thermo Fisher Scientific). These included albumin, IgG, IgA, IgM, IgD, IgE, kappa and lambda light chains, α-1-antitrypsin, α-1-acid glycoprotein, α-2-macroglobulin, apolipoprotein A1, fibrinogen, haptoglobin, and transferrin. Four immunoaffinity columns were equilibrated at room temperature by end-over-end agitation at 13 rpm (MultiBio RS-24, BIOSAN, Riga, Latvia), 1 h before sample application. Each column was loaded with 70 μL of untreated blood plasma. The columns were incubated for 15 min with end-over-end agitation at 13 rpm. The bottom of the column was opened to collect the flow-through by centrifugation at 1,000 x *g* for 1 min (Heraeus Multifuge X 1R centrifuge, Thermo Scientific, Osterode, Germany). This instrument was used for all centrifugation steps. The slurry was resuspended by adding 1 mL PBS (8 g/L NaCl_(aq)_, 0.2 g/L KCl_(aq)_, 0.2 g/L K_2_HPO_4(aq)_, 1.15 g/L Na_2_HPO_4(aq)_ pH adjusted to 7.4). The flow-through was recovered in the same tube by centrifuging at 1,000 x *g* for one min. The flow-through from four columns was pooled and referred to as “top 14-HAP depleted sample”. To avoid protein precipitation during the subsequent desalting process, SDS detergent to a final concentration 0.01% (v/v) was added to the top 14-HAP depleted sample. This protein sample was concentrated 10 times using a 3.5 kDa Amicon Ultra 15 mL device (#UFC500324, Merck) at 3,000 x g 80 minutes. The initial concentration of NaCl in PBS (130 mM) was reduced to a concentration below 0.7 mM NaCl by two centrifugation steps of 3,000 x g for 60 min, adding 10 mL Milli-Q water and one last centrifugation step of 3,000 x g for 30 min adding 3 mL Milli-Q water. The sample volume was reduced to approximately 400 μL using a rotational vacuum concentrator (RVC 2-33 CO plus, Christ, Osterode am Harz, Germany) at 1 °C, 0.1 mbar, approximately 2 h. This instrument was used for all drying steps. The sample was quantified by a Pierce™ BCA Protein Assay Kit (#23225, Thermo Fisher Scientific) and stored at −80°C.

**Figure 1.**
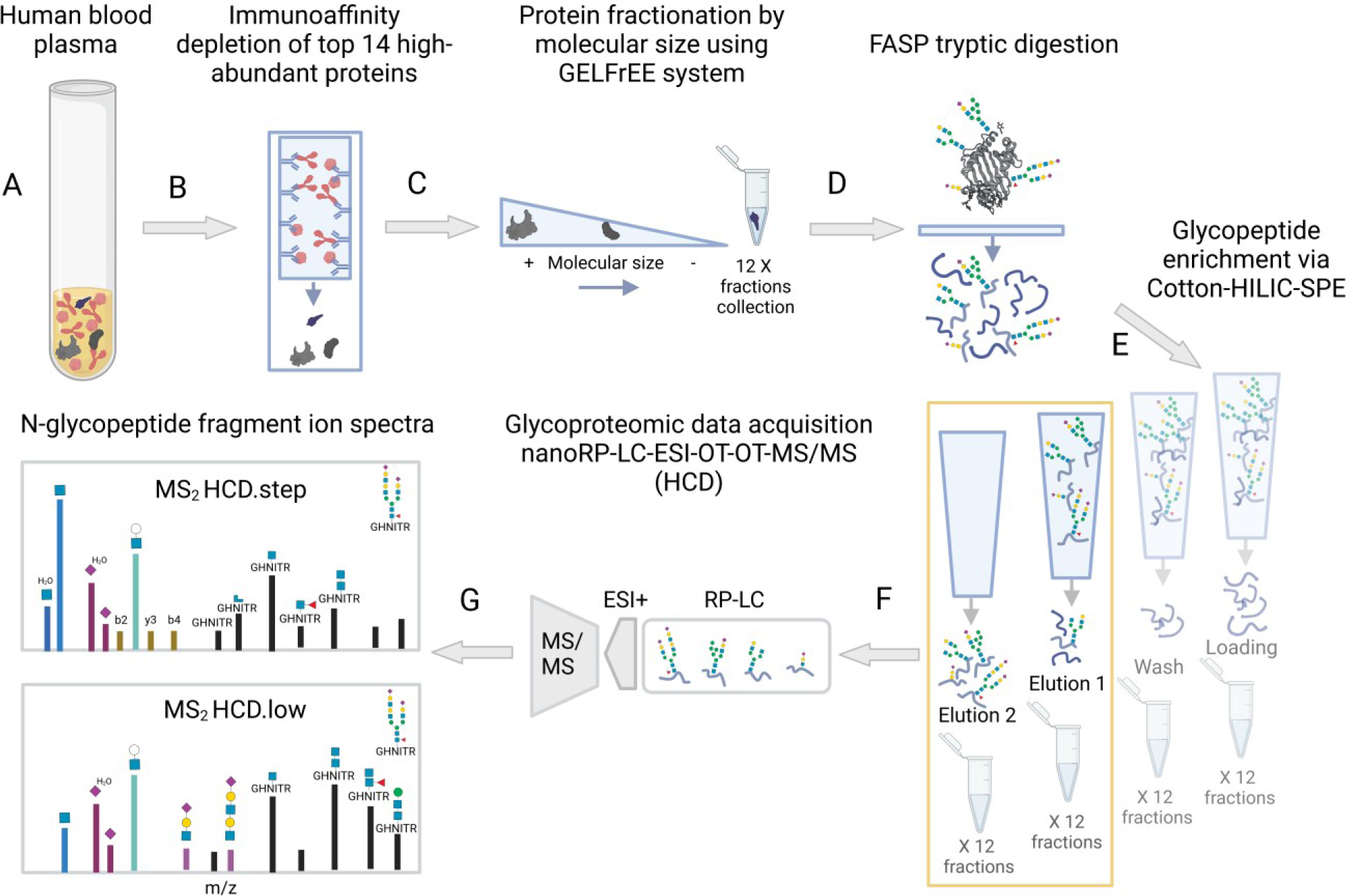
Sample preparation workflow and measurement. (**A**) Blood plasma/serum sample. (**B**) Immunoaffinity depletion of top 14 high-abundant blood plasma proteins. (**C**) Separation of top 14-HAP-depleted fraction by size through GELFrEE® system. Collection of 12 fractions. (**D**) Tryptic digestion via filter aided sample preparation (FASP) (**E**) Glycopeptide enrichment via cotton hydrophilic interaction liquid chromatography solid-phase extraction (Cotton-HILIC-SPE). Collection of four HILIC-fractions per protein-fraction. (**F**) Data acquisition through reverse-phase liquid chromatography electrospray ionization tandem mass spectrometry using orbitrap for both precursor and fragment ions. (**G**) Two MS/MS analyses were generated for each glycopeptide-enriched fraction: higher-energy collisional dissociation (HCD) step and low. Image created with BioRender.com

The sample fractionation by protein size was conducted through an eight-channel GELFrEE® 8100 Fractionation System (GELFrEE® 8100 Fractionation System, 8% Cartridge Kit, #42103, Abcam, Cambridge, UK). The cartridge was prepared according to the manufacturer instructions. For sample preparation, we used 800 μg of the desalted and concentrated sample, adjusting the volume to 448 μL with Milli-Q water, adding 120 μL of GELFrEE® acetate sample buffer and 32 μL 1 M DTT_(aq)_. The sample mixture was heated at 50 °C for 10 min. An amount of 200 µg protein sample was loaded in each cartridge channel. The electrophoresis method was programmed to generate 12 fractions. The electrophoresis loading-step started at 50 V and continued for 16 min. Then, 12 fractions were collected in the following 152.5 min. The amount of protein per fraction was determined by a Pierce™ BCA Protein Assay (Thermo Fisher Scientific).

Tryptic digestion was performed using the FASP (filter aided sample preparation) method, described in Hoffmann *et al*. [26,37]. This digestion protocol was also applied on untreated blood plasma and the top 14-HAP depleted sample using 60 μg of protein. For all filtration steps, 10 kDa Nanosep® Omega filters (OD010C35, Pall®) were used. The reagents, 0.4 M DTT and 0.55 M IAA, were dissolved first in 50 mM ABC_(aq)_ buffer pH 7.8 (ABC buffer) immediately before use and diluted 10 times in 8 M Urea_(aq)_ and 100 mM Tris-HCl_(aq)_ pH 8.5 (urea buffer). After each washing step, the filters were shaken and then centrifuged at 14,000 x g for 10 min. From each fraction derived from the GELFrEE®system, an amount of 60 μg of protein was loaded onto the filter and washed twice with 200 μL urea buffer. Then, a volume of 100 µL 40 mM DTT_(aq)_ was added and incubated for 20 min at 56 °C and 300 rpm on a ThermoMixer C (Eppendorf, Hamburg, Germany). After centrifugation at 14,000 x g for 10 min, 100 µL of 55 mM IAA_(aq)_ was added. The sample was incubated for 20 min in the dark, at room temperature and 300 rpm. Once reduced and carbamidomethylated, the protein sample was washed three times with 100 µL urea buffer, and three times with 100 μL ABC buffer. Trypsin was added at a ratio 1:60 (mg enzyme: mg protein sample) in a final volume of 100 μL 50 mM ABC_(aq)_ + 5% (v/v) ACN + 10 mM CaCl_(aq)_. The sample was incubated over night at 37 °C and 300 rpm (incubator Titramax 1000, Heidolph, Schwabach, Germany). The digest was recovered and collected by centrifugation. The membrane was washed once with 50 μL 50 mM ABC_(aq)_ +5% (v/v) ACN, and once with 50 μL water. The flow-through was kept and combined with the digest. The digest was divided into aliquots with 20 μg of peptides. The digests were dried using the rotational vacuum concentrator (1°C, 0.1 mbar, approximately 2 h).

Glycopeptide enrichment via cotton-hydrophilic interaction liquid chromatography-solid phase extraction (Cotton-HILIC-SPE) was based on a protocol from Selman *et al.* [38]. Cotton-tips were produced in-house by introducing a mercerized-cotton thread of 3 mm length into 250 μL RANNIN tips (Pipette Tips RT LTS 250µL SX 768A/8 Mettler Toledo). The washing solution (85% (v/v) ACN_(aq)_ 0.1% (v/v) TFA_(aq)_) and first-elution solution (78% (v/v) ACN_(aq)_ 0.1% (v/v) TFA_(aq)_) were prepared immediately before use. For each GELFrEE-fraction, 20 μg of lyophilized tryptic digest was solubilized in 50 μL 85% (v/v) ACN_(aq)_. The thread-filled tips were cleaned three times by pipetting 100 μL water and disposing the volume. The tips were equilibrated by pipetting five times 100 μL washing-solution. The peptide sample was loaded into the cotton-tips by slowly pipetting up and down the liquid 20 times. The liquid from the loading-step was discharged in a clean tube and kept (loading fraction). The tips were washed three times by pipetting 100 μL washing-solution into a clean tube (wash fraction). The elution 1 fraction was recovered by pipetting three times 100 μL first-elution solution into a clean tube. The elution 2 fraction was recovered by pipetting six times 100 μL H_2_O into a clean tube.

The four fractions were dried in vacuum and stored at −20°C. This Cotton-HILIC-SPE protocol was also applied on 20 μg of tryptic peptides from the untreated blood plasma and the top 14-HAP depleted sample. The HILIC-fractions were solubilized in 20 μL 2% (v/v) ACN_(aq)_ 0.1% (v/v) TFA_(aq)_ on the day of injection. Four microliter of sample were injected per LC-MS/MS run.

### 2.2. nanoRP-LC-ESI-OT-OT-MS/MS data dependent acquisition

Nano-reversed-phase liquid chromatography electrospray ionization orbitrap tandem mass spectrometry (nanoRP-LC-ESI-OT-OT-MS/MS) was performed using a Dionex UltiMate 3000 RSLCnano system (UHPLC, Thermo Scientific) coupled online to an Orbitrap Elite Hybrid Ion Trap-Orbitrap Mass Spectrometer. The UHPLC system was equipped with a C18 trap column (length 2 cm, particle size 5 µm, pore size 100 Å, inner diameter 100 µm, Acclaim PepMap™ 100, #164199 Thermo Scientific) and a C18 separation column (length 25 cm, pore size 100 Å, particle size 2 μm, inner diameter 75 μm, Acclaim PepMap™ RSLC nanoViper #164941, Thermo Scientific). For the separation gradient, the nano pump mobile phase A (2% (v/v) ACN_(aq)_, 0.1% (v/v) FA_(aq)_) and mobile phase B (80% (v/v) ACN_(aq)_, 10% (v/v) TFE_(aq)_, 0.1% (v/v) FA_(aq)_) were controlled at a flow rate of 300 nL/min at 40 °C. Then, 5 min after the sample injection, the loading pump mobile phase A (2% (v/v) ACN_(aq)_ 0.05% (v/v) TFA_(aq)_, flow rate 7 µL/min) switched the port valve to 1-2, connecting trap and separation column. The separation gradient was 4% B for 4 min, to 35% in 62 min, to 90% in 2 min, 90% for 6 min, to 4% in 2 min, and 4% for 24 min. Both precursor and fragment ion scans were acquired using orbitrap mass analyzer (OT-OT-MS/MS). Each sample was measured twice to acquire data of the glycopeptides fragmented at two higher-energy collisional dissociation (HCD) regimes: HCD.low with fixed (NCE) of 20 and HCD.step with stepped NCE of 35 (width 15%, 2 steps). Both data dependent-scan methods were acquired in positive mode, selecting the top 5 peaks with 10 sec dynamic exclusion, scan range 350–2000 m/z, and isolation width 4 m/z. orbitrap mass analyzer was used for both precursor ion scan (MS^1^) and fragment ion scan (MS^2^) selecting 30,000 and 15,000 resolution for each scan, respectively.

### 2.3. Data analysis

The HCD.step measurements of the HILIC-fractions (loading, wash, elution1, and elution 2) derived from the untreated blood plasma, the top 14-HAP depleted sample, and the top 14-HAP depleted and fractionated sample were searched for peptides through the human UniProtKB/SwissProt database (v2021-11-30, 20,306 canonical sequences) using Proteome Discoverer (version 2.5.0.400, Thermo Fisher Scientific). The parameters applied to the search engines Sequest HT and Mascot were set as follows: full specific tryptic digestion, two missed cleavages allowed, precursor and fragment mass tolerance: 10 ppm and 0.02 Da, respectively. Deamidation (N, Q), oxidation (M), acetyl / +42.011 Da (protein N*-*Terminus) were established as dynamic modifications, and carbamidomethyl / +57.021 Da (C) as static modification. Percolator was applied for peptide validation, 1% FDR on peptide level was applied.

The workflow of glycoproteomic data analysis is depicted in **Figure 2**. The Byonic™ search engine (v4.2.10 by Protein Metrics, Cupertino, CA, USA) was used to search for *N-*glycopeptides in the HCD.step and HCD.low measurements. HCD.step generates both, abundant peptide fragment ions (a, b, y) and some fragments ions from the peptide-linked glycan moiety (Y ions). This fragmentation energy generates the following four characteristic *N-*glycopeptide Y ions: (I) [M_peptide_+H-NH_3_]^+^, (II) [M_peptide_+H]^+^, (III) [M_peptide_+H +^0.2^X HexNAc]^+^ and (IV) [M_peptide_+H +HexNAc]^+^ (where NH_3_=17.0265 Da, ^0.2^X HexNAc=83.0371 Da; and HexNAc=203.0794 Da). The HCD.low MS^2^ spectra are rich in glycan fragment ions and Y ions with a longer portion of the *N-*glycan attached, which allows *N-*glycan structure interpretation. The following parameters were applied in Byonic: specific tryptic digestion, maximum two missed cleavages, cysteine carbamidomethylation as fixed peptide modification, methionine oxidation as common-1 modification, and asparagine deamidation and pyro Glu/Gln as rare-1 dynamic modifications. Precursor and fragment mass tolerance was set to 10 ppm and 20 ppm, respectively. Recalibration lock mass was 445.1201 m/z. Fragmentation-type was set to QTOF/HCD. The human canonical proteome UniProtKB/Swiss-Prot database (20,396 reviewed-sequences downloaded in May 2021) was applied for protein identification. For *N-*glycan identification, a customized *N-*glycan database of 288 compositions was applied. During the Byonic search, no decoy search was performed and there were no cuts on protein FDR. The *N*-glycoproteomic analysis of untreated blood plasma and top 14-HAP depleted sample (HCD.step acquisitions) was conducted using the same parameters.

**Figure 2.**
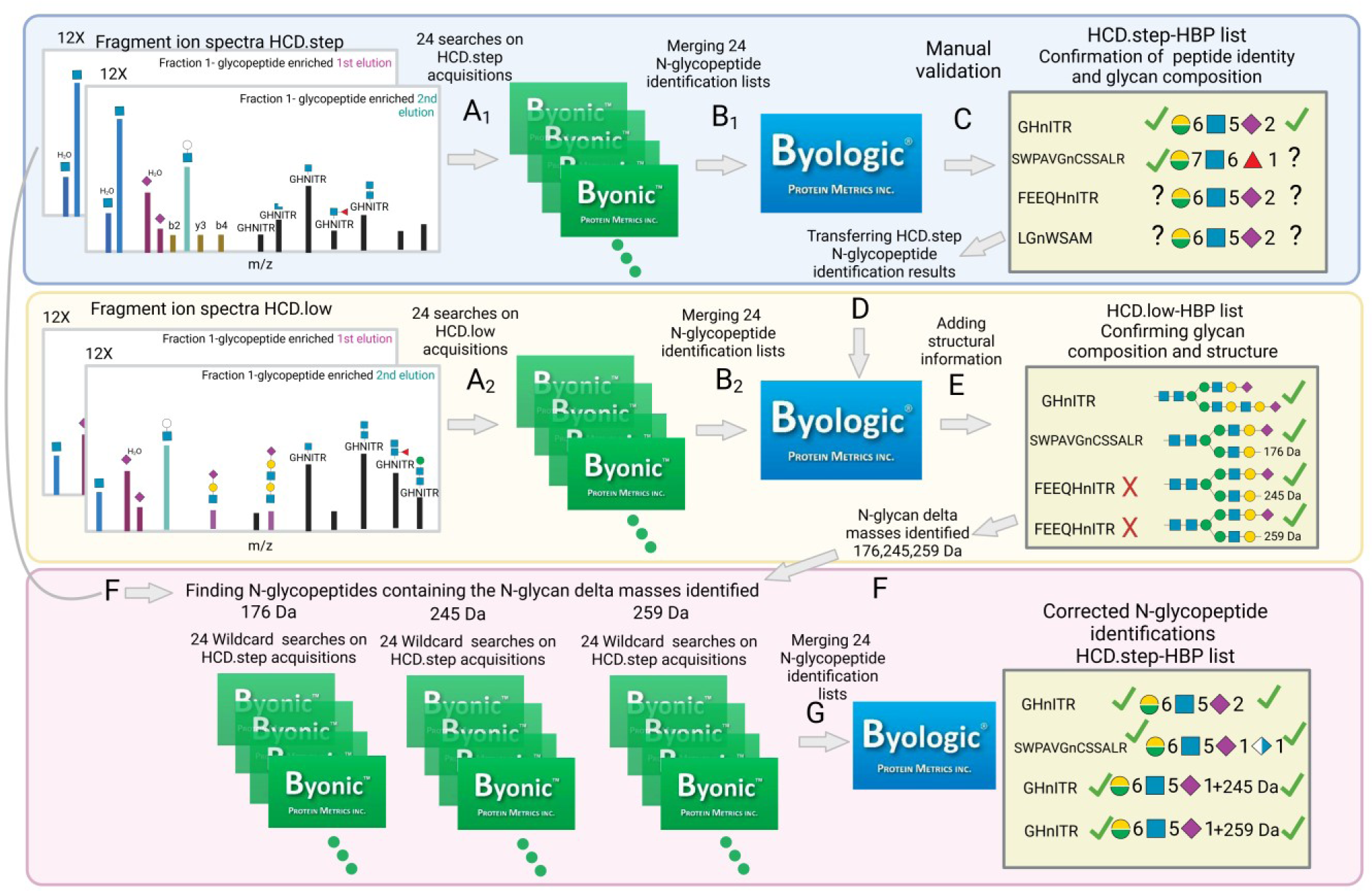
Human blood plasma (HBP) *N-*Glycoproteomic data analysis workflow. The intact *N-*glycopeptides, present in 24 glycopeptide enriched fractions (12 x elution 1 + 12 x elution 2) were analyzed by LC-MS/MS using both fragmentation energies, HCD.step and HCD.low. (**A**) The spectra file acquired at each fragmentation regime was searched for *N-*glycopeptides using Byonic™ software. (**B**) All individual searches corresponding to the same fragmentation energy were combined using Byologic™. After this step, two large *N-*glycopeptide identification lists were produced: HCD.step-HBP list and HCD.low-HBP list. (**C**) The *N-*glycopeptides contained in the HCD.step-HBP list were manually validated to confirm the peptide and *N-*glycan composition assigned by the software. (**D**) The *N-*glycopeptide-identifications confirmed were imported into the HCD.low-HBP list to substitute corresponding precursor ions potentially incorrectly identified, using their mass and retention time. (**E**) Features annotated in the HCD.step-HBP list were transferred to the HCD.low-HBP list and an additional revision for *N-*glycan structural evidence is conducted in the HCD.low-HPB list. (**F**) A second Byonic search focused on the new features identified was triggered. (**G**) Finally, a second HCD.step-HBP list containing corrected *N-*glycopeptide identifications was generated using Byologic™. Image created with BioRender.com.

The database with 288 entries of *N-*glycan compositions resulted from combining the “182 *N-*glycan Byonic database” with reported compositions, including multiple fucoses and compositions of phosphorylated or sulfated *N-*glycans deduced through a diagnostic search. The diagnostic search was performed on each HCD.step file applying two layout-filters created in Thermo Xcalibur Qual Browser (Version 2.2, Thermo Scientific) software (**Supplementary Figures S1** and **S2**). It allowed filtering MS^2^ spectra containing oxonium marker ions for sulfated *N-*glycans HexNAc_1_Sulfo_1_ [M+H]^+^ or HexNAc_1_Hex_1_Sulfo_1_ [M+H]^+^ and phosphorylated *N-*glycans Hex_1_Phospho_1_ [M+H]^+^ or Hex_2_Phospho_1_ [M+H]^+^. In the filtered MS^2^ spectra, the presumable peptide mass was determined by detecting the four characteristic Y ions ((I)-(IV)). Then, the corresponding *N-*glycan mass was calculated by subtracting the peptide mass from the precursor ion mass. The sulfated and phosphorylated *N-*glycan compositions were deduced using GlycoMod tool from expasy.org [39]. The compositions from sulfated and phosphorylated *N-*glycans were appended to the *N-*glycan database.

The search results derived from HCD.low and HCD.step acquisitions were imported into Byologic (v4.4-74-g75311a1df5 x64, Protein Metrics) in two sets. The searches corresponding to HCD.step acquisitions generated one Byologic™ file (HCD.step-HBP) and the ones corresponding to HCD.low acquisitions generated a second Byologic™ file (HCD.low-HBP). Manual glycopeptide validation was performed only on the HCD.step-HBP list using Byologic. The validated *N-*glycopeptide identifications in the HCD.step file were transferred to the HCD.low-HBP Byologic™ file in order to substitute the corresponding correct precursor ion identifications in the HCD.low-HBP list. This substitution was performed using Peptide Manager Byonic function. The validated *N-*glycopeptide identifications in HCD.step-HBP file were exported using “Export insilico to CSV” option. Then, the CSV file was imported into the HCD.low-HBP Byologic™ file through Peptide Manager “Intersect CSV” function. The parameters set for intersecting the corresponding *N-*glycopeptides were precursor mass error (10 ppm) and retention time error (5 minutes). Once transferred to the HCD.low-HBP Byologic™ file, the *N-*glycopeptide identifications were manually revised and additional structural information on the corresponding *N-*glycans was annotated.

Rare *N-*glycan compositions were identified during *N-*glycopeptide validation. This relates to three *N-*glycan building block masses that were not included in the first *N-*glycan database. In order to find the gPSMs for such identifications, the Byonic wildcard search function was applied. This function allows adding a delta mass within a user specified range. The mass ranges searched here were: 176, 245, and 259 [±1 Da]. These ranges were narrowed to the delta masses deduced during the validation (176.0314 Da, 245.0524 Da, and 259.0672 Da). One specific wildcard search was applied for each delta mass to all the HCD.step files. To reduce the search space, MS/MS filtering function was applied, allowing MS^2^ spectra containing at least two of the following masses: HexNAc_1_Hex_1_ [M+H]^+^/366.1395, HexNAc_1_ [M+H]^+^/204.0867, HexNAc_1_ [M-H_2_O+H]^+^/186.0761 or NeuAc_1_ [M-H_2_O+H]^+^/274.0921, mass tolerance 0.02 Da. In the wildcard search, the delta mass is assigned to the glycan modification and not the peptide sequence by setting the parameter restrict to residues to “g” (where “g” means glycan). For both wildcard searches, the same parameters as the first Byonic search were applied: specific tryptic digestion, maximum 2 missed cleavages, cysteine carbamidomethylation as fixed peptide modification, and methionine oxidation as common-1 modification, asparagine deamidation and pyro Glu/Gln as rare-1 dynamic modifications. Precursor and fragment mass tolerance was 10 ppm and 20 ppm, respectively. Recalibration lock mass was 445.1201 m/z. Fragmentation-type set to “QTOF/HCD”. To speed up the search, a shorter protein sequence list and *N-*glycan compositions list including only the elements present in the “True”-validated *N-*glycopeptide results were applied.

## 3. Results

During in-depth *N-*glycoproteomic analysis of human blood plasma proteins, one encounters two main causes of complexity. First, a few high-abundant proteins suppress the signal of hundreds of proteins present at lower abundance. Second, the heterogeneous and unpredictable nature of protein *N-*glycosylation requires sufficient evidence for describing both, the *N-*glycan structure and their position in a particular protein. In this study, we established a sample preparation and glycoproteomic data analysis workflow that enabled to reach the very low-abundant glycoproteome. Furthermore, it allowed the detection and site-specific description of rare *N-*glycan compositions and the identification of structural features linked to blood plasma glycoproteins at high confidence.

### 3.1. Exploring blood plasma (glyco)proteins at low concentration range and N-glycan micro-heterogeneity

To evaluate the performance of our workflow for identifying middle- and low-abundant human blood plasma proteins, a proteomic analysis was conducted. For each step of the sample preparation workflow, the number of proteins identified was compared (acceptance criterion: ≥ two unique peptides per protein). The protein identifications were linked to the blood plasma concentrations reported in Plasma Proteome Database (PPD) using visProteomics R package [40,41]. As presented in **Figure 3A**, the concentration of most of the proteins observed by analyzing the untreated blood plasma ranges from 1·10^9^ down to 1·10^6^ pg/mL. After depleting the top 14-HAP, the concentration range of the proteins detected was extended and covered now from 1·10^9^ down to 3·10^5^ to pg/mL. However, by integrating both, top 14-HAP depletion and protein size fractionation, the concentration range of the proteins identified could be further extended and the detection limit further lowered, reaching now from 1·10^9^ down to 6·10^3^ pg/mL. Moreover, the cysteine-rich secretory protein 3 could be identified at the very low concentration of 6.31 pg/mL. The list of proteins identified after each fractionation step is shown in the **Supplementary Tables S1-S3**.

**Figure 3.**
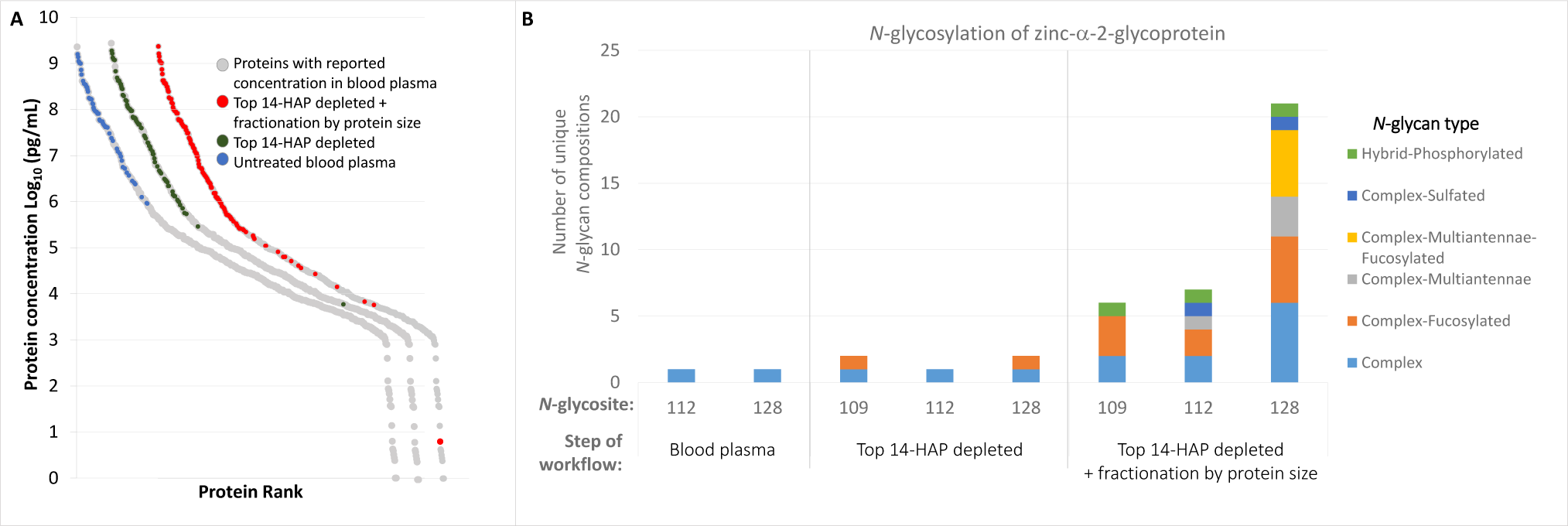
Gain obtained by applying the developed sample preparation workflow. (**A**) Concentration range of the proteins identified after a proteomic analysis of the sample at each step of the preparative workflow (**B**) Measurement of micro-heterogeneity of glycosylation for zinc-α-2-glycoprotein (P25311) after a glycoproteomic analysis at each step of the preparative workflow. Reported concentration of plasma proteins [40].

The developed sample preparation workflow also resulted in a significant improvement regarding the measurement of micro-heterogeneity of glycosylation. Exemplarily, in **Figure 3B**, *N-*glycan compositions detected on each specific-site of zinc-α-2 glycoprotein are compared at each step of the workflow. For site N_128_, for instance, 10 times more *N-*glycan compositions could be detected after the top 14-HAP depletion, compared to the direct analysis of blood plasma. Moreover, even 20 times more *N-*glycan compositions, when combining top 14-HAP depletion and protein size fractionation (**Supplementary Table S4**). Therefore, the new workflow effectively boosts the detection of a large variety of *N-*glycans in each site. A detailed comparison of the *N-*glycopeptides identified after each step of the workflow is shown in **Supplementary Tables S5-S7**. An overview of the *N-*glycopeptide identification results displayed in the **Supplementary Figure S3**, shows that the most common *N-*glycan attached to blood plasma glycoproteins is a diantennary complex-type *N-*glycan disialylated and non-fucosylated.

Overall, by comparing the (glyco)proteomic results after each step of the preparative workflow, we demonstrate that the developed workflow is successfully expanding the range for identification of blood plasma (glyco)proteins from middle to very low-concentration range and enabling a deeper description of the micro-heterogeneity of *N-*glycosylation.

### 3.2. Development of a data analysis workflow for the identification of intact N-glycopeptides

A dedicated data analysis workflow was developed and applied to validate the data generated by in-depth *N-*glycoproteomic LC-MS/MS analysis (**Figure 2**). The workflow relies on the revision of two *N-*glycopeptide lists grouped by the fragmentation energy applied: HCD.step and HCD.low. The first part of the workflow uses HCD.step spectra for confirming the correctness of the peptide sequence and the *N-*glycan composition within the gPSM suggested by Byonic. In the second part, HCD.low fragment ion spectra are used to confirm the proposed *N-*glycan composition. Information about the *N-*glycan structure is manually added, for example, to clarify if a fucose is linked to the core or to the antenna of an *N-*glycan. After acknowledging errors, such as missed *N-*glycan compositions, the third part of the workflow focuses on correcting gPSM that were classified as uncertain identifications by *N-*glycan *de novo* sequencing or performing additional glycoproteomic searches.

The first part of the data analysis workflow is conducted according to the decision tree shown in **Figure 4A**. With this decision strategy, a total of 7,867 gPSM were classified in three main categories: “True”, “Uncertain”, and “False”. A “True” gPSM has evidence on three aspects: 1) it is an *N-*glycopeptide, 2) the mass of the peptide moiety is consistent with the Y ion [M_peptide_+H]^+^, and 3) the oxonium ions observed are coherent with the *N-*glycan composition suggested. An “Uncertain” gPSM meets the first requirement but fails in any of both others. A “False” gPSM does not fulfill the first requirement since it corresponds to a non-glycosylated peptide or an *O-*glycopeptide. The latter is recognized by the ratio between the intensity of HexNAc_1_ [M-H_2_O+H]^+^ and HexNAc_1_ [M+H]^+^ oxonium ions [26]. In an *N-*glycopeptide-derived MS^2^ spectrum (HCD.step) the intensity of HexNAc_1_ [M+H]^+^/204.0867 will be between three to ten times the intensity of HexNAc_1_ [M-H_2_O+H]^+^/186.0761 [26]. As described in **Figure 4A**, each main category acquires a more specific classification according to the evidence observed. Each of the subcategories is exemplified and described in **Supplementary Figures S4-S14**. For example, the MS^2^ spectra from a gPSM classified as “True-Evidence” contain at least three characteristic Y ions and oxonium ions consistent with the *N-*glycan composition presented in the gPSM, yet the MS^2^ spectra might lack peptide b and y ions. The category “Uncertain-change glycopeptide” is also an *N-*glycopeptide identification, but the mass of the characteristic Y ions disagrees with the one expected from the gPSM suggested by the software.

**Figure 4.**
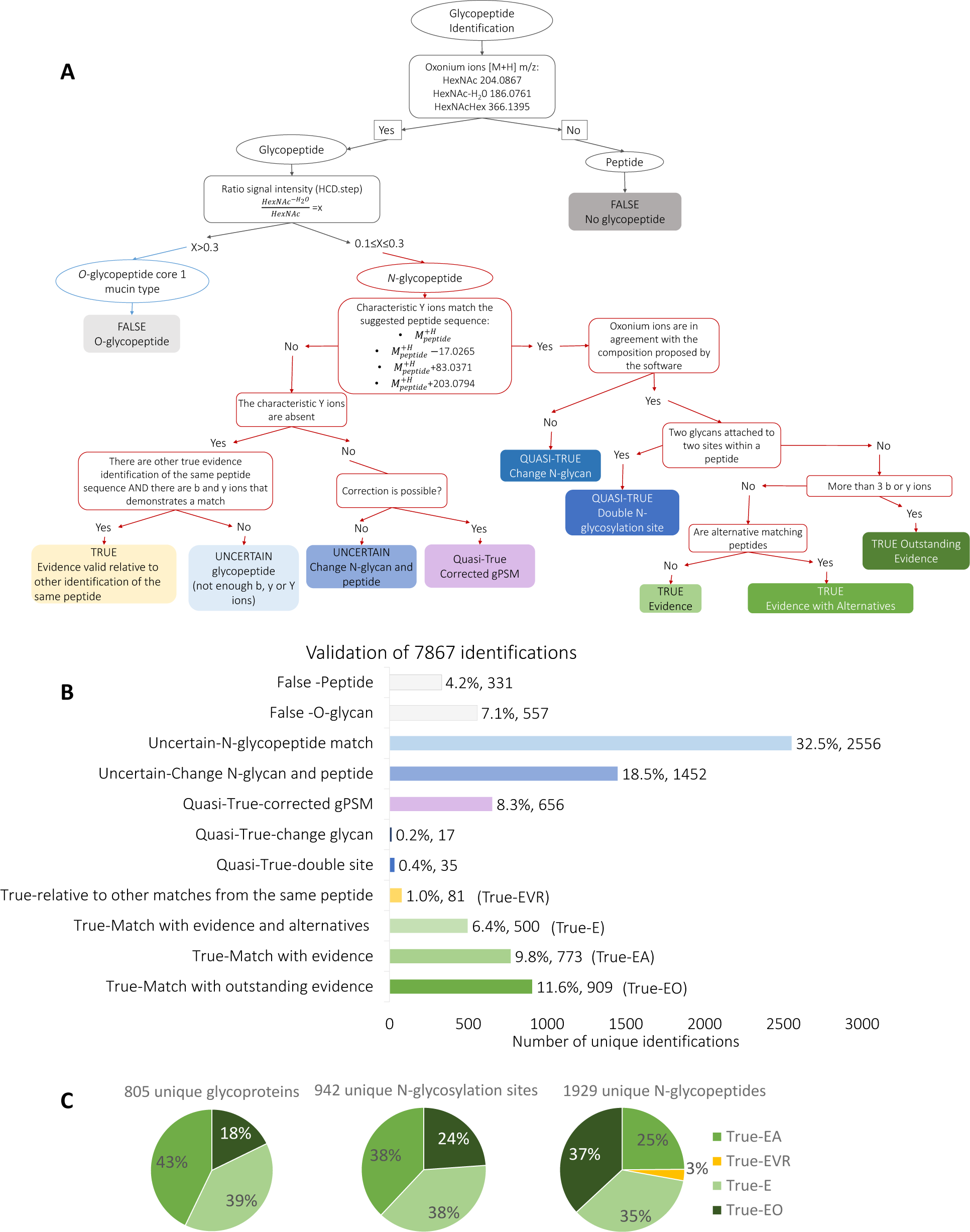
Validation of *N-*glycopeptide identifications. (**A**) Decision tree depicting the process for validating the gPSMs identified in the HCD.step-HBP list. (**B**) Number of gPSM assigned to each validation category. (**C**) Proportion of unique glycoproteins, *N-*glycosylation sites, and *N-*glycopeptides obtained from the true gPSM. T-EO: True Outstanding evidence. T-E: True-Evidence. T-EA: True-Evidence and Alternatives. T-EVR-True Evidence Valid Relative to other identification of the same peptide.

### 3.3. Validated N-glycopeptide identifications in human blood plasma

Figure 4B shows the results of the manual validation of the HCD.step-HBP list, where 88.7% of the gPSM are *N-*glycopeptide identifications. The “False” cases (non *N-*glycopeptides) represent only 11% of the dataset. The results from the validation of HCD.step-HBP are listed in **Supplementary Table S7**. From the total of 7,867 gPSMs, 2263 were correct concerning peptide and *N-*glycan composition (27.9% of the total gPSMs). The total “True” gPSMs are spread across four subcategories: 909 “True matches with outstanding evidence” (T-EO), 773 “True matches with evidence” (T-E), 500 “True matches with evidence and alternative identifications” (T-EA) and 81 “True matches with evidence relative to other matches from the same peptide sequence” (T-EVR). One third of the total identifications correspond to “Uncertain-no-evidence” identifications, in which gPSM feature MS^2^ spectra with a poor number of fragment ions. Another 17% of all gPSMs are incorrect matches (category “Uncertain-change glycopeptide”), which MS^2^ spectra might require different glycoproteomic searches. Some “Uncertain-change glycopeptide” were corrected by comparing their MS^2^ spectra with the ones from “True” gPSM with similar precursor ion mass or retention time. Thus, 656 manually corrected gPSM resulted in the “Quasi-true corrected gPSM” subcategory. Other “Quasi-true” categories, “Change glycan” and “Double site”, represents only 0.6% of the total gPSMs and are cases in which the peptide moiety is correct, but the *N-*glycan cannot be confirmed. As it is displayed in Figure 4C, overall 942 different *N-*glycosites belonging to 805 human glycoproteins where found. Focusing on the different glycoforms per glycosylation site, the total number of gPSM is condensed to 1,929 disregarding the peptide modifications or missed cleavages.

### 3.4. Annotating structural features of bisecting, fucosylated, and LacNAc extended N-glycans using HCD.low spectra

Only the 2263 “True” gPSM were applied on the second part of the workflow for the annotation of *N-*glycan structural information in the HCD.low-HBP list (**Supplementary Table S8**). *N-*glycan structural evidence was screened in HCD.low-spectra corresponding to the validated gPSM, by using the *N-*glycan marker ions listed in **Table 1**. In this table, Y ions are considered in different charge state (z=1–3^+^). This is the second part of the data analysis workflow, which enables the differentiation between antenna and core fucose, multi-antennary *N-*glycans and repeated LacNAc units, or antenna HexNAc and bisecting HexNAc [8, 12, 14–18]. The detection of the Y-ion peptide+HexNAc_3_Hex_1_ ion, for instance, suggests a bisecting *N-*glycan. In total, 16 bisected *N-*glycopeptides were identified in 11 glycoproteins. Most of these bisected *N-*glycans are linked to the proteins IgA2 (heavy chain), IgG1 (heavy chain), IgG2 (heavy chain), and synaptojanin-1. In case of a LacNAc repeat unit, the observation of the oxonium-ion HexNAc_2_Hex_2_NeuAc_1_ (B-ion [M+H]+/1022.3671) was detected and annotated in the HCD.low-HBP list [5]. Of note, the oxonium ion HexNAc_2_Hex_2_ (B-ion [M+H]+/731.2717) was not applied because this fragment ion is also generated by complex-type and bisected *N-*glycans. Therefore, the identification of non-sialylated LacNAc repeat units was not included. It was found that apolipoprotein D harbors three different *N-*glycans with a sialylated LacNAc repeat unit at glycosylation site N_65_, being the protein with highest frequency of this type of *N-*glycan. Three additional *N-*glycopeptides, belonging to kininogen-1, β-2-glycoprotein 1, and kallistatin, were also found to have LacNAc repeat units.

**Table 1.**
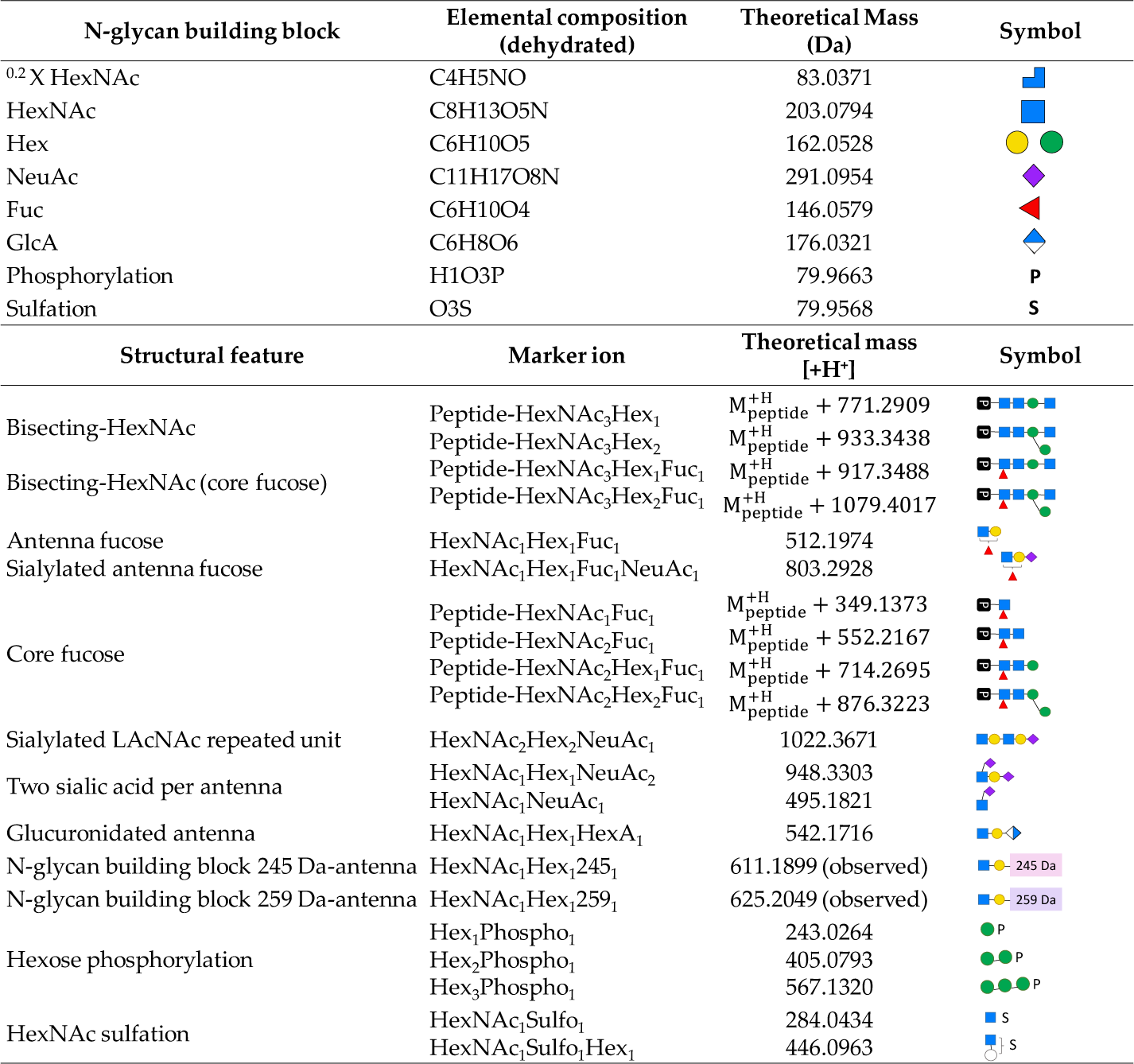
*N*-Glycan-derived marker ions. B and Y ions applied in this study.

| N-glycan building block | Elemental composition (dehydrated) | Theoretical Mass (Da) | Symbol |
| --- | --- | --- | --- |
| <sup>0.2</sup>X HexNAc | C<sub>4</sub>H<sub>5</sub>NO | 83.0371 |  |
| HexNAc | C<sub>8</sub>H<sub>13</sub>O<sub>5</sub>N | 203.0794 |  |
| Hex | C<sub>6</sub>H<sub>10</sub>O<sub>5</sub> | 162.0528 |  |
| NeuAc | C<sub>11</sub>H<sub>17</sub>O<sub>8</sub>N | 291.0954 |  |
| Fuc | C<sub>6</sub>H<sub>10</sub>O<sub>4</sub> | 146.0579 |  |
| GlcA | C<sub>6</sub>H<sub>8</sub>O<sub>6</sub> | 176.0321 |  |
| Phosphorylation | H<sub>1</sub>O<sub>3</sub>P | 79.9663 | P |
| Sulfation | O<sub>3</sub>S | 79.9568 | S |
| Structural feature | Marker ion | Theoretical mass [H<sup>+</sup>] | Symbol |
| Bisecting-HexNAc | Peptide-HexNAc<sub>3</sub>Hex<sub>1</sub> | M<sub>peptide</sub><sup>H</sup> + 771.2909 |  |
|  | Peptide-HexNAc<sub>3</sub>Hex<sub>2</sub> | M<sub>peptide</sub><sup>H</sup> + 933.3438 |  |
| Bisecting-HexNAc (core fucose) | Peptide-HexNAc<sub>3</sub>Hex<sub>1</sub>Fuc<sub>1</sub> | M<sub>peptide</sub><sup>H</sup> + 917.3488 |  |
|  | Peptide-HexNAc<sub>3</sub>Hex<sub>2</sub>Fuc<sub>1</sub> | M<sub>peptide</sub><sup>H</sup> + 1079.4017 |  |
| Antenna fucose | HexNAc<sub>1</sub>Hex<sub>1</sub>Fuc<sub>1</sub> | 512.1974 |  |
| Sialylated antenna fucose | HexNAc<sub>1</sub>Hex<sub>1</sub>Fuc<sub>1</sub>NeuAc<sub>1</sub> | 803.2928 |  |
| Core fucose | Peptide-HexNAc<sub>1</sub>Fuc<sub>1</sub> | M<sub>peptide</sub><sup>H</sup> + 349.1373 |  |
|  | Peptide-HexNAc<sub>2</sub>Fuc<sub>1</sub> | M<sub>peptide</sub><sup>H</sup> + 552.2167 |  |
|  | Peptide-HexNAc<sub>2</sub>Hex<sub>1</sub>Fuc<sub>1</sub> | M<sub>peptide</sub><sup>H</sup> + 714.2695 |  |
|  | Peptide-HexNAc<sub>2</sub>Hex<sub>2</sub>Fuc<sub>1</sub> | M<sub>peptide</sub><sup>H</sup> + 876.3223 |  |
| Sialylated LacNAc repeated unit | HexNAc<sub>2</sub>Hex<sub>2</sub>NeuAc<sub>1</sub> | 1022.3671 |  |
| Two sialic acid per antenna | HexNAc<sub>1</sub>Hex<sub>1</sub>NeuAc<sub>2</sub> | 948.3303 |  |
|  | HexNAc<sub>1</sub>NeuAc<sub>1</sub> | 495.1821 |  |
| Glucuronidated antenna | HexNAc<sub>1</sub>Hex<sub>1</sub>HexA<sub>1</sub> | 542.1716 |  |
| N-glycan building block 245 Da-antenna | HexNAc<sub>1</sub>Hex<sub>1</sub>245<sub>1</sub> | 611.1899 (observed) |  |
| N-glycan building block 259 Da-antenna | HexNAc<sub>1</sub>Hex<sub>1</sub>259<sub>1</sub> | 625.2049 (observed) |  |
| Hexose phosphorylation | Hex<sub>1</sub>Phospho<sub>1</sub> | 243.0264 |  |
|  | Hex<sub>2</sub>Phospho<sub>1</sub> | 405.0793 |  |
|  | Hex<sub>3</sub>Phospho<sub>1</sub> | 567.1320 |  |
| HexNAc sulfation | HexNAc<sub>1</sub>Sulfo<sub>1</sub> | 284.0434 |  |
|  | HexNAc<sub>1</sub>Sulfo<sub>1</sub>Hex<sub>1</sub> | 446.0963 |  |

In our HCD.step-HBP list, 679 unique fucosylated *N-*glycopeptides were further reviewed for deducing the position of fucose in the respective *N-*glycans (**Supplementary Table S7**). The presence of the following marker ions was registered in the HCD.step and HCD.low-HBP lists: for core-fucose, peptide+HexNAc_1_Fuc_1_ [M+H]^+^ (Y-ion); and for antenna fucose, HexNAc_1_Hex_1_Fuc_1_[M+H]^+^ (oxonium ion, B-ion). In total, we found 350 unique fucosylated *N-*glycopeptides having MS^2^ spectra with either a core or an antenna fucose marker ion (**Supplementary Table S9**). Ambiguous cases occurred when only one fucose was suggested in the *N-*glycopeptide identification but both classes of marker ions, antenna- and core-fucose, were observed. Since fucose transfer (fucose rearrangement) in the gas-phase can occur from core to antenna generating “false” antenna fucose marker ions, the *N-*glycopeptides including one fucose but presenting both antenna- and core-fucose ion were set as core-fucose only [42]. In the case of *N-*glycan compositions with more than one fucose presenting both antenna- and core-fucose ions, both core and antenna fucosylation were granted. For *N-*glycopeptides where core-fucose ions were not observed but antenna-fucose ion was detected, only antenna fucosylation was accepted. As a result, we found 242 core-fucosylated *N-*glycopeptides, 47 core- and antenna-fucosylated *N-*glycopeptides, and 61 *N-*glycopeptides presenting only one or more antenna-fucose. The proteins with the highest frequency on antenna-fucosylation are zinc-α-2-glycoprotein, α-2-HS-glycoprotein, β-2-glycoprotein 1, clusterin and ceruloplasmin. In Figure 5, all gPSM including fucose(s) in the *N-*glycan composition were plotted, in order to analyze the frequency of fucosylated *N-*glycopeptides with and without fucose marker ion evidence. It was observed that a high number of difucosylated gPSMs, which lack of any kind of fucose ion, presented complex-type diantennary monosialylated or triantennary disialylated *N-*glycans. A prominent example of a difucosylated gPSMs is shown in Figure 6A, where no antenna-fucose oxonium ions are observed in the annotation done by the software. By analyzing the isotopic pattern of the triply charged precursor of this example (Figure 6B), it is noticeable that there is an initial isotopic peak with an m/z difference of 0.3331 with respect to the monoisotopic peak assigned by the software. This presumes that the mass of the precursor ion deduced by the software is one dalton bigger than the real mass of such a precursor ion (Δm/z=0.3331, z=3, Δm=0.9993 Da). Then, the mass difference between assigning 2xFucoses instead of 1xNeuAc is one dalton (2xFuc=292.1158 Da, 1xNeuAc=291.0954 Da, Δ +1.0204 Da). Hence, we hypothesized that the software misrepresented the MS^1^ isotopic pattern, since it must start from a previous isotopic peak (Figure 6D). This would explain the mass difference of one dalton, which leads to incorrectly assign two fucoses instead of one neuraminic acid. To confirm this, the Figure 6C shows the manual *de novo* sequencing of the *N-*glycan composition reflected in the HCD.low spectrum from this precursor ion. As fucose ion evidence was not observed, we searched here for neutral losses corresponding to fucose. The result shows only a delta mass corresponding to a doubly charged NeuAc and a neutral loss from a second NeuAc at the end of the m/z range. This demonstrates that this *N-*glycan ion contains two NeuAc instead of one NeuAc and two fucoses. Other examples in which the potential real monoisotopic peak of the precursor ion is ignored in gPSM with multiple fucoses are shown in **Supplementary Figure S15**.

**Figure 5.**
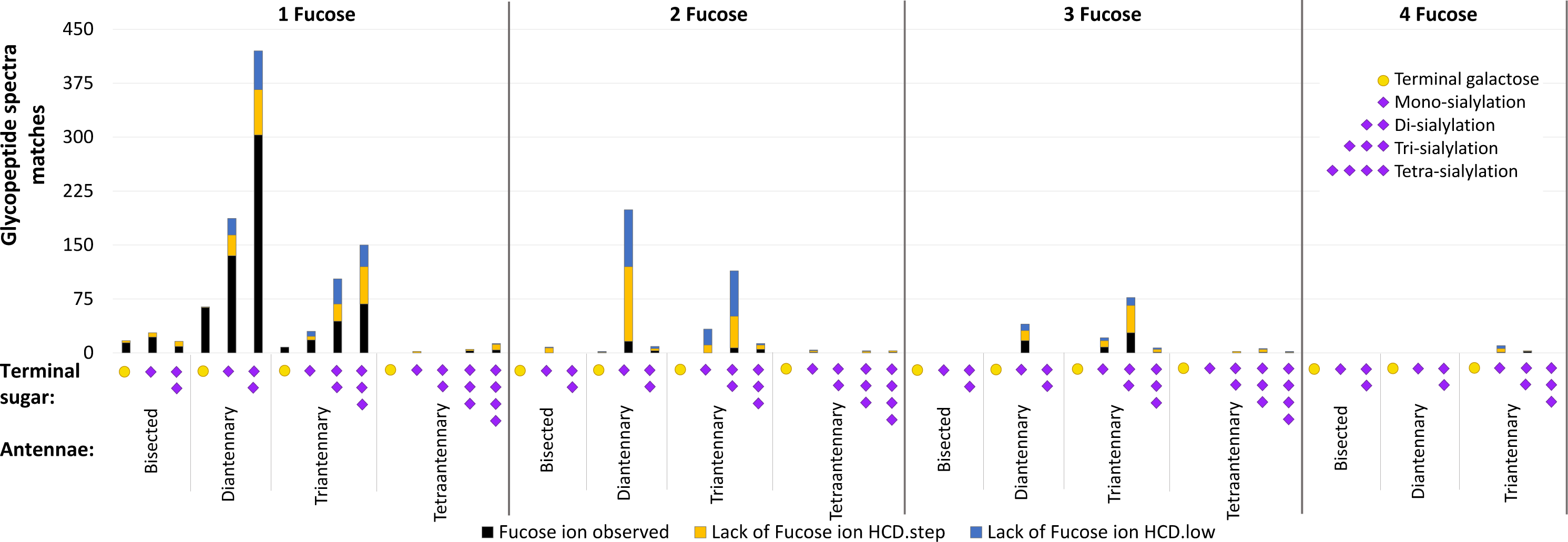
Evaluation of evidence present in gPSMs including fucose. Distribution of all gPSM including fucose identified by Byonic software in both HCD.step and HCD.low-HBP lists (with and without fucose ion evidence observed).

**Figure 6.**
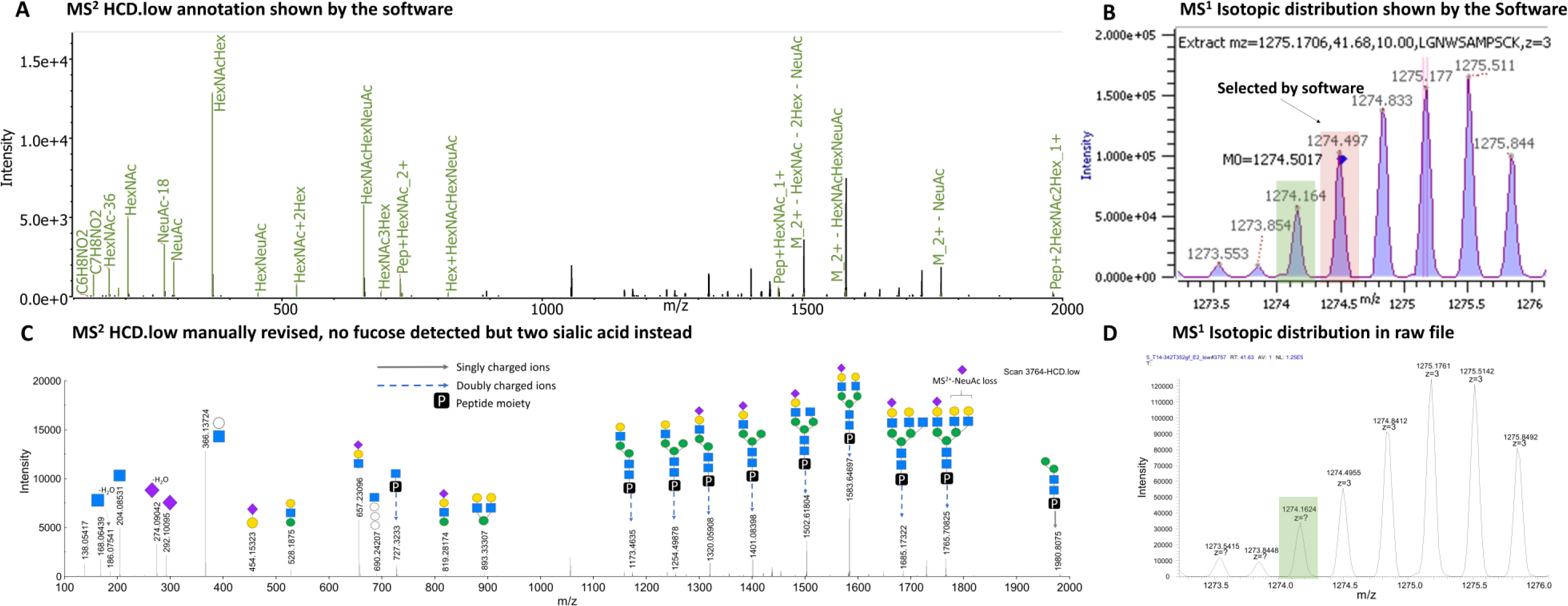
Manual revision of an incorrect identification for β-2-glycoprotein 1 (peptide K.LGnWSAMPSCK.A) containing a difucosylated complex-type *N-*glycan. (**A**) MS^2^ HCD.low annotation proposed by Byonic software assigning an incorrect *N-*glycan HexNAc(5)Hex(6)Fuc(2)NeuAc(1) *N-*glycan mass: 2570.9250 Da. (**B**) MS^1^ Isotopic distribution with monoisotopic peak selected by Byonic software (red). Precursor ion m/z 1274.5017, z=3, observed mass 3820.4789 Da. Correct precursor ion highlighted in green. (**C**) MS^2^ HCD.low manual annotation assigning the correct *N-*glycan HexNAc(5)Hex(6)NeuAc(2) *N-*glycan mass: 2569.9046 Da. (**D**) MS^1^ Isotopic distribution observed in the raw file. Correct precursor ion highlighted in green. Precursor ion suggested m/z 1274.1624, z=3, calculated mass 3819.4872 Da.

### 3.5. Detection of sulfated and phosphorylated N-glycopeptides in common human blood plasma glycoproteins

*N-*Glycopeptides with sulfated and phosphorylated *N-*glycans are difficult to detect and locate not only due to their low-abundance, but also due to their unstable behavior during MS analysis [15,43]. Still, by integration of a dedicated diagnostic search using Thermo Xcalibur Qual Browser, we were able to detect *N-*glycopeptides with phosphorylated and sulfated *N-*glycan compositions in the human blood plasma (**Supplementary Figures S1 and S2**). A collection of all possible phosphorylated and sulfated *N-*glycan compositions was added to the Byonic *N-*glycan composition database. This database was then used for all *N-*glycopeptide searches. Additionally, during manual validation, both the HCD.step-and HCD.low-HBP lists were revised with regard to marker ions related to sulfation or phosphorylation. The example of Figure 7 shows that the HCD.low MS^2^ spectrum of a sulfated *N-*glycopeptide identification contained the HexNAc_1_Hex_1_Sulfo_1_ [M+H]^+^ oxonium marker ion, while the HCD.step MS^2^ spectrum of the same *N-*glycopeptide displays the HexNAc_1_Sulfo_1_ [M+H]^+^ oxonium marker ion (software annotation **Supplementary Figure S16**). Interestingly, in none of the acquired MS² spectra the Hex_1_Sulfo_1_ [M+H]^+^ oxonium ion was detected. With regard to phosphorylated *N-*glycopeptides (Figure 8), the HCD.low MS^2^ spectrum mainly shows the Hex_1_Phospho_1_ [M+H]^+^ and Hex_2_Phospho_1_ [M+H]^+^ oxonium marker ions, while the HCD.step MS^2^ spectra dominantly produce the Hex_1_Phospho_1_ [M+H]^+^ oxonium marker ion (software annotation in **Supplementary Figure S17**). Among the *N-*glycopeptide identifications from the HCD.step-HBP list, 65 contain sulfated *N-*glycans but only 10 of them contain sulfation marker ions (**Supplementary Table S10**). In the HCD.step-HBP list, 15 *N-*glycopeptides presented phosphorylated *N-*glycans, and in contrast to the sulfated *N-*glycopeptides, all the MS^2^ spectra contain marker ion(s) for phosphorylated hexose (**Supplementary Table S11**).

**Figure 7.**
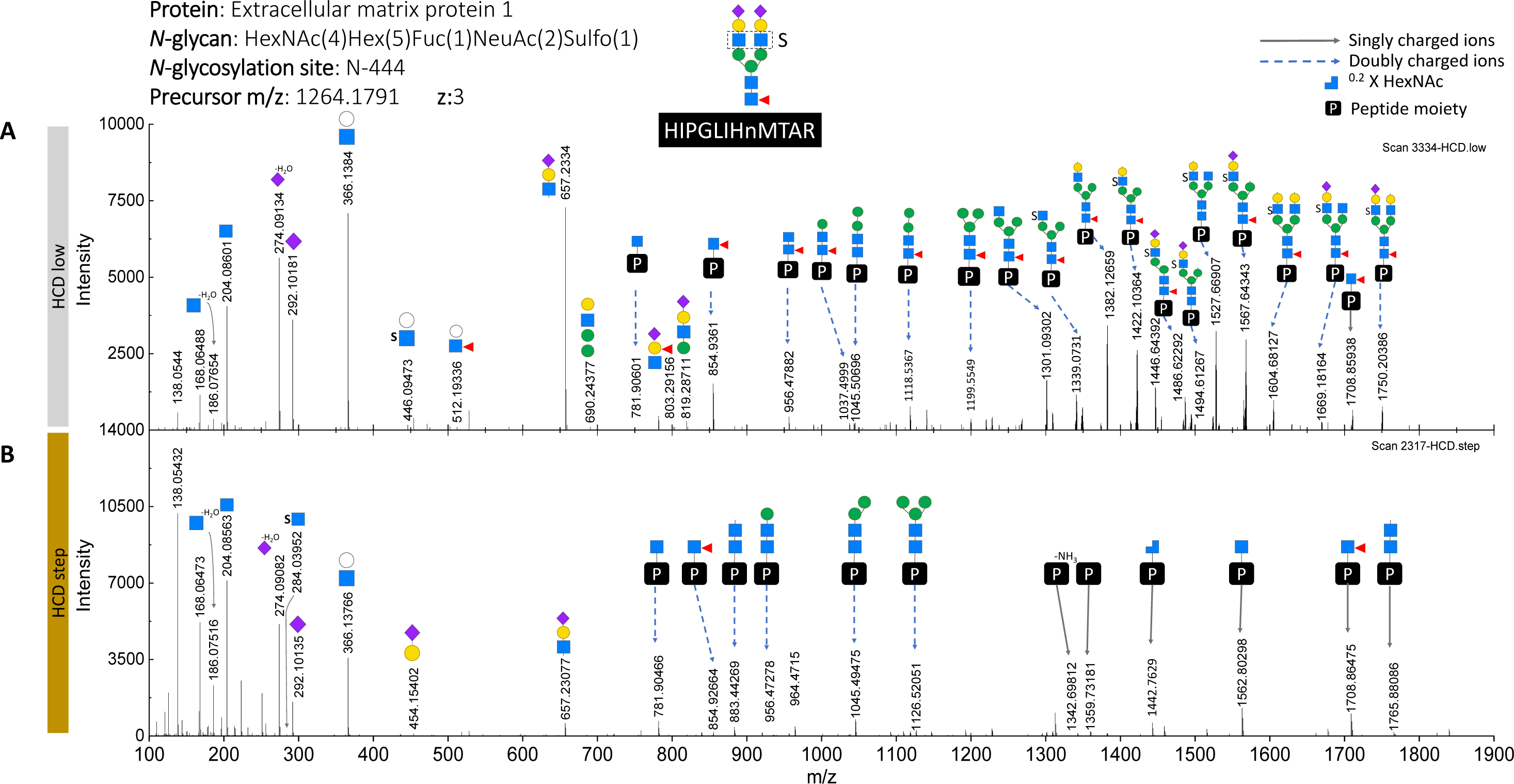
Annotation of fragment ions released from sulfated *N-*glycopeptide. (**A**) HCD.low fragment ion spectrum showing sulfation position. (**B**) HCD.low fragment ion spectrum showing oxonium ion evidence related to HexNAc sulfation.

**Figure 8.**
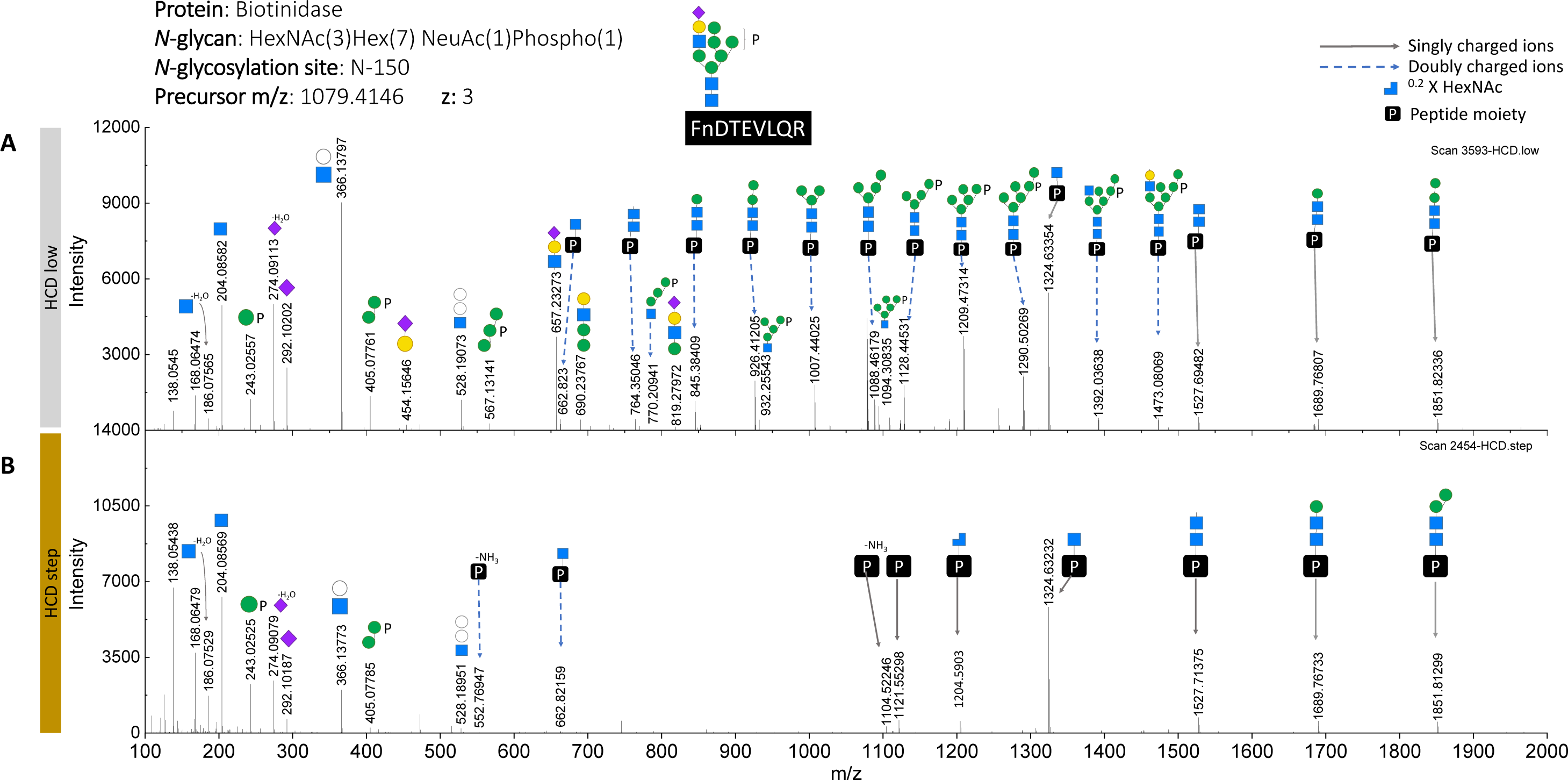
Annotation of fragment ions released from phosphorylated *N-*glycopeptide. (**A**) HCD.low fragment ion spectrum showing phosphorylation position. (**B**) HCD.low fragment ion spectrum showing oxonium ion evidence related to hexose phosphorylation.

As the examples in Figure 7 and **8** show, through *de novo* sequencing of the HCD.low MS^2^ spectra, sulfation was detected on the antenna-HexNAc and phosphorylation after the fourth mannose of the hybrid-type *N-*glycan. In the lists of validated gPSMs containing sulfation or phosphorylation (**Supplementary Tables S10 and S11**), it is observed that the sulfated *N-*glycan compositions are related to complex-type *N-*glycans, while phosphorylated *N-*glycans are related to hybrid-type or oligomannose-type *N-*glycan. The *N-*glycosylation micro-heterogeneity of some proteins, like ceruloplasmin, hemopexin, heparin cofactor 2 and zinc α-2-glycoprotein, reflected several *N-*glycosylation sites harboring a sulfated *N-*glycan (**Supplementary Table S10**). The proteins ceruloplasmin, hemopexin and zinc α-2-glycoprotein also contained a higher frequency of phosphorylated *N-*glycans (**Supplementary Table S11**).

Glycoproteomic analysis of the untreated blood plasma did not achieved the detection of phosphorylated nor of sulfated *N-*glycopeptides at all (**Supplementary Figure S18**). After the analysis of the top 14-HAP depleted sample we detected one sulfated *N-*glycan attached to N_762_ from ceruloplasmin and one phosphorylated hybrid *N-*glycan linked to N_187_ of hemopexin. In contrast, by combining the immunoaffinity depletion of top 14-HAP plus the fractionation by protein size, our sample preparation workflow reached the detection of phosphorylated *N-*glycans in 11 glycoproteins and sulfated *N-*glycans in 54 glycoproteins. Moreover, it allowed the identification of a phosphorylated *N-*glycan in a very-low abundant protein cysteine-rich secretory protein LCCL domain-containing 1 (8.2 pg/mL)[44,45]. Even though sulfated *N-*glycopeptides tend to lose the sulfated marker ions due to their lability, we confirmed 10 glycoproteins holding sulfated-HexNAc *N-*glycans. This demonstrates the power of the developed workflow for the site-specific identification of modified *N-*glycans.

### 3.6. Rare N-glycans in human blood plasma proteins

As displayed in the Figure 9, three rare *N-*glycan building blocks with the masses 176.0314 Da, 245.0524 Da and 259.0672 Da were found during a process of *de novo* sequencing of HCD.low MS^2^ spectra of previously incorrect gPSM. As described in the following paragraphs, it is hypothesized that the first mass corresponds to glucuronic acid, while the two last possibly resemble other types of sialic acid. All the identifications with these masses show complex diantennary monosialylated *N-*glycans. While one antenna is capped by NeuAc, the second is capped by any of the rare *N-*glycan building block (Figure 9). This is supported by the presence of the oxonium ions generated in each case: Hex_1_HexNAc_1_+176.0314 [M+H]^+^/542.1694, Hex_1_HexNAc_1_+245.0524 [M+H]^+^/611.1899, and Hex_1_HexNAc_1_+259.0672 [M+H]^+^/625.2049. Furthermore, site-specific identification of these *N-*glycans revealed their occupancy on N_121_ from prothrombin (**Supplementary Figure S19**).

**Figure 9.**
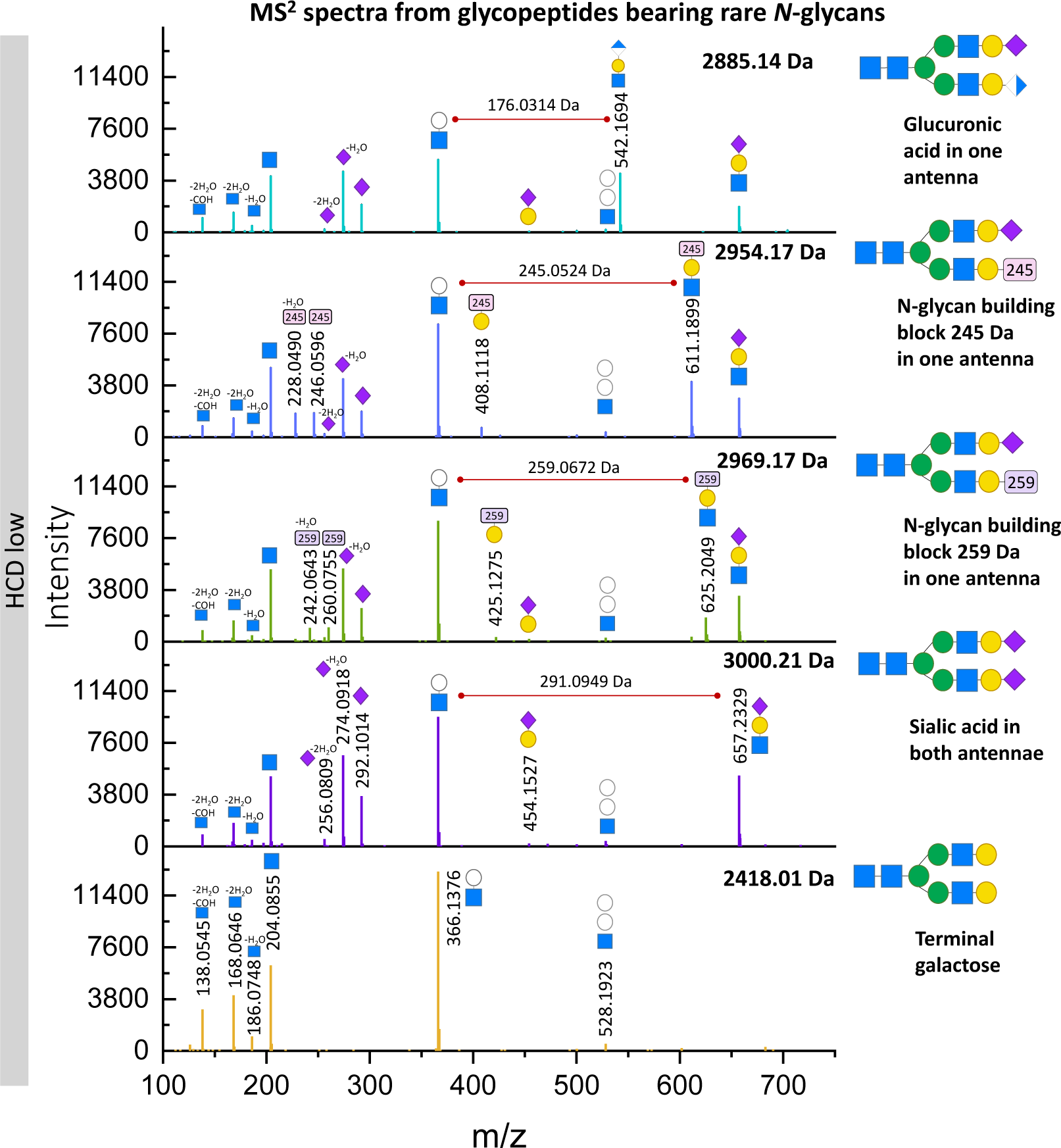
Annotation of fragment ions released from rare *N-*glycans attached to N121 (peptide GHVNITR).

The composition of the oxonium ion: Hex_1_HexNAc_1_GlcA_1_ [M+H]^+^ (542.1716 [M+H]^+^, where GlcA refers to glucuronic acid) matches the observed m/z 542.1694 [M+H]^+^ (mass error −4.06 ppm). The ion Hex_1_HexNAc_1_GlcA_1_ [M+H]^+^ has been reported as a fragment derived from 3-sulfonated glucuronidated *N-*glycans (HNK-1) common in human brain [46]. In our case, the existence of the HNK-1 glycoepitope is rejected, since no fragment ion nor neutral loss representing GlcA sulfation was detected. Moreover, the mass of the elemental composition of GlcA (C6H8O6 after water loss) is 176.0321 Da and the observed neutral mass is 176.0314 Da. This results in an error of −3.98 ppm between both masses, which favors the hypothesis that the first *N-*glycan building blocks identified is glucuronic acid. In order to find other proteins harboring glucuronidated *N-*glycans, a new Byonic wildcard search was performed (detailed in Materials and Methods section). As a result, ten glucuronidated *N-*glycopeptides derived from β-2-glycoprotein 1, α-2-HS-glycoprotein, kallikrein, histidine-rich glycoprotein, prothrombin, hemopexin and complement factor H were identified (**Supplementary Table S12**). No glucuronidated *N-*glycopeptides were detected by the analysis of the untreated human blood plasma (**Supplementary Figure S18**). After the analysis of top 14 HAP-depleted sample only the *N-*glycosylation site N_453_ from hemopexin was found to harbor a glucuronidated complex *N-*glycan. These comparisons demonstrates that the here developed sample preparation workflow supports the identification of considerably rarer *N-*glycopeptides.

To our knowledge, the terminal *N-*glycan building blocks 245.0524 and 259.0672 Da have not been described so far. Assuming that these molecules might be a variant of sialic acid, the molecules were searched in PubChem, proposing their hydrated masses (263.20 and 277.23 Da, respectively). The candidates with the highest similarity to Neu5Ac (C11H19NO9, 309.27 Da [47]) were selected and submitted to CFM-ID 4.0, an online tool to predict fragment ion spectra [48]. The software predicted the fragment ion spectrum using NCE 10 and 20 V (NCE 20 V applied here for HCD.low measurements). After comparing the predicted and the average observed fragment ions, the two candidate molecules shown in **Supplementary Figure S19** were selected. The first candidate (C9H13NO8, 263.0641 Da [49]) theoretically generates the B ions 246.0608 [M+H]^+^ and 228.0503 [M+H-H_2_O]^+^, which are also observed in the MS^2^ spectra that contains the building block “245 Da” (oxonium ions 246.0596 and 228.0490 [M+H]^+^). This results in an error of −4.99 ppm and −5.53 ppm, respectively. The second candidate (C10H15NO8, 277.0798 Da [50]) produces the oxonium ions 260.0765 [M+H]^+^ and 242.0659 [M+H-H_2_O]^+^. These B ions are observed in the MS^2^ spectra where the mass “259 Da” is present (oxonium ions 260.0755 and 242.0643 [M+H]^+^). Comparing the error between the theoretical and observed fragment ions m/z for the second candidate, it results in −3.77 ppm and −6.65 ppm, respectively. The low error supports the hypothesis that the *N-*glycan building blocks found might be two molecules similar to NeuAc, such as the ones proposed in **Supplementary Figure S19** (molecules drawn in PubChem Sketcher[51]). Both *N-*glycan building blocks were only found on N_121_ of prothrombin (**Supplementary Figure S20** and **Supplementary Table S13**) and have not been reported in this nor in other proteins before.

During manual annotation of *N-*glycan structural features in the HCD.low-HBP list, we found disialo-antennary *N-*glycan structures as shown in Figure 10. The surprising presence of one antenna holding two sialic acids was evident by the B ion HexNAc_1_Hex_1_NeuAc_2_ [M+H]^+^/948.3303. Typically, sialic acid is linked to galactose, while the second sialic acid might be linked to this first sialic acid or to the antenna-HexNAc. In order to describe this disialo-antennary structure, the MS^2^ HCD.low spectra from two *N-*glycopeptides, corresponding to N_762_ ceruloplasmin and N_65_ apolipoprotein D, were *de novo* sequenced. In both MS^2^ spectra the linkage of sialic acid to the HexNAc is demonstrated by the presence of the peptide-HexNAc_3_Hex_3_NeuAc_1_ or the peptide-HexNAc_4_Hex_4_NeuAc_2_ fragment ions ([M+H]^2+^ or [M+H]^3+^). In addition, the fragment ion spectra from apolipoprotein D show the oxonium marker ion HexNAc_1_NeuAc_1_ [M+H]^+^, supporting the existence of the unexpected HexNAc-NeuAc linkage. Overall, ten proteins presented a disialo-antennary *N-*glycan structure including ceruloplasmin, α-1-antichymotripsin, apolipoprotein D and β-2-glycoprotein 1. All the *N-*glycopeptides presenting disialo-antennary *N-*glycans have the compositions: HexNAc(5)Hex(6)NeuAc(3) or HexNAc(5)Hex(6)Fuc(1)NeuAc(3). Interestingly, Figure 10 also describes the *N-*glycan structure of a disialo-antennary *N-*glycan attached to β-2-glycoprotein 1. This *N-*glycopeptide not only shows one disialo-antenna but also a LAcNAc repeat unit on the other antenna. The aforementioned oxonium marker ions are absent in the HCD.step fragment ion spectra annotated by the software and displayed in the **Supplementary Figure S21**. This demonstrates that only by including HCD.low fragmentation, more *N-*glycan features are collected to detect relevant glycoepitopes.

**Figure 10.**
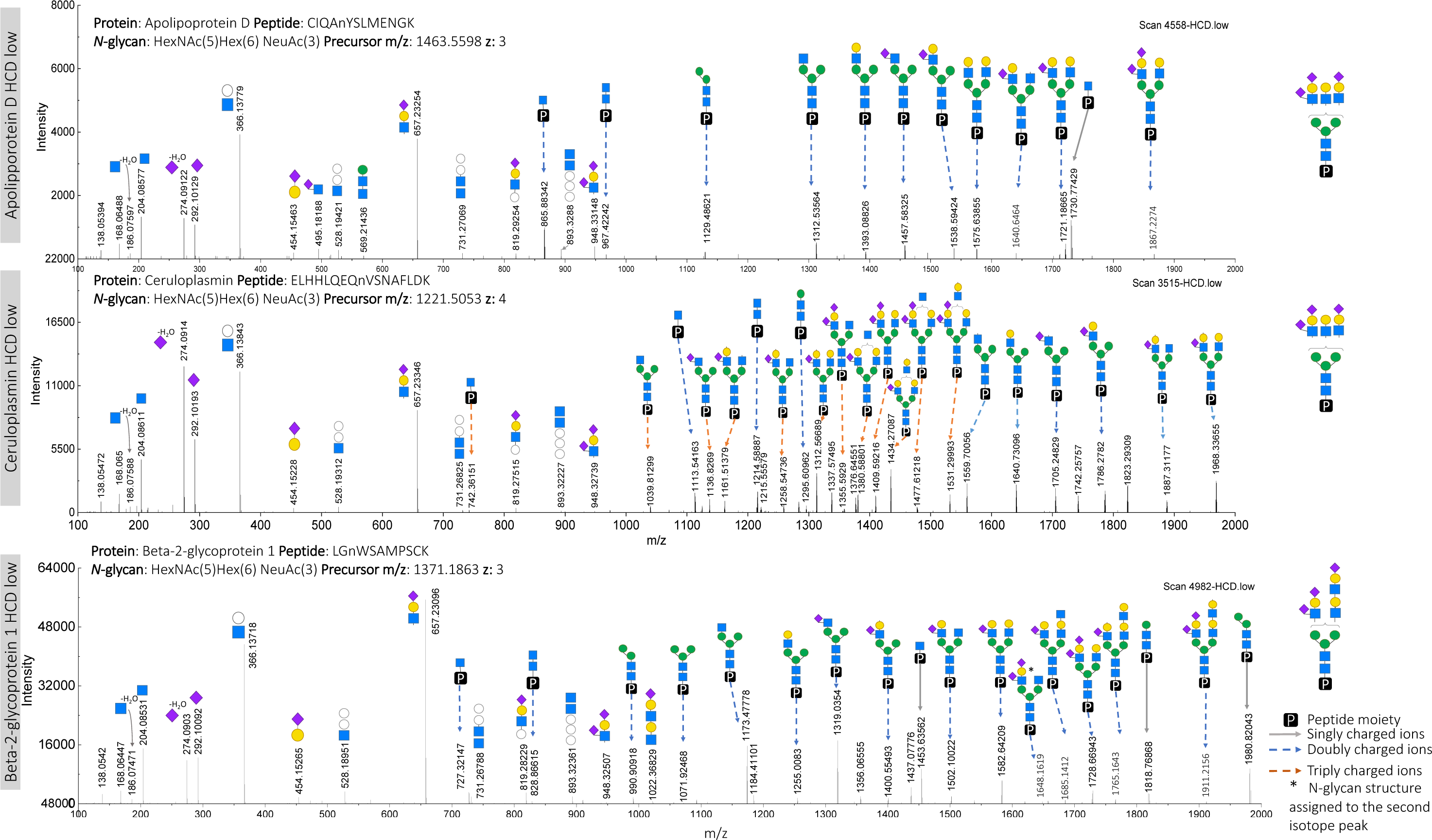
*N-*Glycan de novo sequencing of *N-*glycopeptides with disialo-antennary *N-*glycan structure confirming HexNAc-NeuAc linkage

## 4. Discussion

Here, we developed and applied both a sample preparation and a data analysis workflow for an in-depth analysis of the blood plasma intact *N-*glycopeptides. The internal steps in the workflows are connected as follows: 1) conducting blood plasma top 14-HAP depletion, followed by a protein size fractionation, tryptic digestion and intact glycopeptide enrichment, 2) measuring glycopeptide enriched fractions twice by nanoLC-ESI OT-OT MS/MS HCD.low and HCD.step fragmentation methods, 3) searching for *N-*glycopeptides using software, 4) manually validating *N-*glycopeptides using the newly established decision tree, 5) manually annotating structural *N-*glycan features using HCD.low fragment ion spectra and 6) conducting especial glycoproteomic searches to incorporate missed rare *N-*glycan compositions. The main achievement of this in-depth analysis is the capacity to delve even deeper into the *N-*glycan micro-heterogeneity along with a significantly extended protein concentration range (10^9^–10^1^ pg/mL). This is demonstrated by the observation of low-abundant glycans like phosphorylated *N-*glycans in very low-abundant proteins such as cysteine-rich secretory protein LCCL domain-containing 1 (8.2 pg/mL) [44,45]. Another example is the reliable identification of sulfated *N-*glycans in middle-abundant proteins (e.g. extracellular matrix protein 1, 0.78 μg/mL [40]) through the detection of sulfated fragment ions, typically hard-to-acquire [43]. In addition, the manual validation of all unique *N-*glycopeptide identifications computed by Byonic, led to the detection of glucuronidated and other rare *N-*glycans attached to blood plasma proteins expressed by the liver. Finally, the manual annotation of *N-*glycan structural features revealed the advantages of using HCD.low spectra for recognizing relevant structures or glycoepitopes. After searching through GlyCosmos, UniCarb, and GlyConnect databases, we conclude that there is not yet a site-specific description of the glycoproteins harboring some of the sulfated, phosphorylated, glucuronidated, and disialylated-antenna *N-*glycan structures as identified in our study (listed in **Supplementary Tables S14 and S15**) [52–58]. Then, by performing a proteomic analysis on all fractions after *N-*glycopeptide enrichment (FDR <1%), we demonstrated an expanded detection limit of blood plasma proteins within a concentration range from 1·10^9^ down to 1·10^1^ pg/mL. This evaluation not only demonstrates that our workflow is comparable to other in-depth proteomic workflows [20,59], but also that it has significant advantages for glycoproteomics in blood plasma. In contrast, the study from Wessels H. *et al.*, presented a strategy for the diagnostic of congenital disorders of glycosylation (CDG) via the direct glycoproteomic analysis of untreated blood plasma [60]. Their method resulted in the site-specific profile of 34 proteins, which satisfied the evaluation of the CDG cases selected. Our workflow offers an alternative for evaluating site-specific profiles of low-abundant proteins and other *N-*glycan types, like rare *N-*glycan structures that might be involved in different CDGs. So far, structural *N-*glycan details require glycomics or exoglycosidases and lectins. However, here, we show a new avenue for moving towards structural glycoproteomics. This study aimed to address the limitations and pitfalls caused by the blind spots existing during an *N-*glycoproteomic analysis. By acknowledging these aspects, glycoproteomic bioinformatics tools can be refined to obtain results that are more accurate and comprehensive.

In the recent years, many studies revealed substantial information about the blood plasma *N-*glycoproteome based on the analysis of intact *N-*glycopeptides [20,59,61]. These analyses not only applied multi-step sample preparation workflows but also powerful bioinformatics tools. The biggest study was performed by Shu *et al.*, achieving the identification of 1,036 *N-*glycosites containing 738 *N-*glycans, resulting in 22,677 unique *N-*glycopeptides derived from 526 glycoproteins using pMatchGlyco software (FDR 1%) [59]. In our study, we obtained 7,867 *N-*glycopeptide identifications (no decoys, no cuts on FDR) from which 1,929 were manually validated as “True” identifications revealing 942 different *N-*glycosites, and 805 human glycoproteins. The higher number of *N-*glycopeptide identifications of Shu *et al.,* might result from a library of de-*N-*glycosylated peptide identifications, created by cumulative searches including semi-tryptic digestion and 16 variable peptide modifications. In addition, they explored atypical *N-*glycosylation consensus sequences (N-X-S/T/V/C, X≠P) and a bigger *N-*glycan database (739 *N-*glycans). In contrast, our study resulted from one standard search applying full specific tryptic digestion, four variable peptide modifications, common *N-*glycosylation consensus sequence (N-X-S/T, X≠P), and 288 *N-*glycan compositions. In regard of sample material, Shu *et al.,* used a pooled serum sample from 50 healthy individuals, which might increase the opportunity to accumulate the proteins secreted in blood plasma in different abundances. Additionally, the authors included the high-abundant protein depleted fraction in their study. In comparison, our analysis resulted from a blood plasma sample (pooled from a smaller number of healthy donors) from which the top 14 high-abundant proteins were depleted and later discarded using a single use immunoaffinity column. Analysis of the high-abundant protein fraction was not in the scope of our study, since *N-*glycosylation of these proteins is well studied. Another important limitation in our study was the relatively long measurement time of the instrument used for acquiring data (Orbitrap Elite-Velos, scan speed 4 Hz), compared to Orbitrap Q Exactive mass spectrometer (with three times faster scan speed of 12 Hz [62]) used by Shu *et al*.

Software and parameters for the search are another factor that can lead to different results regarding the interpretation of *N-*glycopeptides. This is shown in the study from Kawahara *et al.*, where two glycoproteomic spectra files derived from human serum were provided to 22 expert-groups in glycoproteomics to evaluate the impact of applying different bioinformatics strategies on the identification of intact *O-* and *N-*glycopeptides [18]. The results showed an enormous variability of glycopeptide identifications, glycoproteins and glycan compositions. The study attributed this inconsistency mainly to the filters applied after the search, such as score threshold or FDR cut. In our study, we have excluded post-search filter interference by disallowing decoys, and FDR cuts. Instead, we conducted a manual validation after the *N-*glycopeptide search. We are aware that this approach could result in incorrect identifications. Nevertheless, our scope was to identify hitches and opportunities given during intact *N-*glycopeptide analysis, especially for incorrect *N-*glycopeptide identifications with high quality-score and ambiguous gPSM. All the identifications were manually scrutinized, evidence supporting the trueness of the identification was annotated and incorrect matches were revised. Thus, a validation subcategory, related to the variability observed by Kawahara *et al*., was the subcategory “True evidence with alternatives”. This subcategory lists *N-*glycopeptides which alternative peptide matches are named in the column “comments” of the table HCD.step-HBP list in **Supplementary Table S7**. In our dataset, 35% of the “True” *N-*glycopeptides are classified within this subcategory and two common features were observed: 1) a low number of b and y ions in the MS^2^ spectra and 2) a low abundance of these observed proteins in blood plasma. The first issue relates to low MS^2^ spectra quality, which is associated to several factors such as precursor ions with poor ionization efficiency, fragment ion losses, and low-abundant precursor ions. The second issue depends on the protein concentration distribution of the sample, where low-abundant proteins are reflected as low-abundant precursor ions in the final spectra. Regarding the second feature, Kreimer *et al*., propose the implementation of algorithms designed for smart selection and acquisition of the typically suppressed precursor ions to reduce the proportion of spectra with poor-quality [63]. Then, these algorithms would reduce the proportion of spectra that lead to the variability of glycopeptide identifications by improving spectra quality during the MS/MS measurement.

Another source of variability among glycoproteomic analyses is fucosylation. After manual validation, we observed a high frequency of identifications presenting multiple-fucose moieties without any fucose-related fragment ion. The occurrence is higher in di-tri-or tetra-antennary *N-*glycopeptides with an incomplete number of capping sialic acid in the antennae. Kawahara *et al.*, reported a similar observation, where the high frequency of *N-*glycopeptide identifications with multiple-fucose moieties, reported by many participants, did not correlate with the results obtained from typical *N-*glycomic analysis of human blood plasma. We observed that the assignment of multiple-fucose moieties instead of one sialic acid is caused by an incorrect detection of the isotopic pattern for the corresponding precursor ion. Lee *et al*. also observed this problem and reported that it might be influenced by the precursor-picking default settings [22]. Hence, special attention must be given to this phenomenon since this problem could also induce the incorrect assignation of other not fucose-related quasi-isobaric *N-*glycan masses, such as HexNAc(5)Hex(6)Fuc(1)NeuAc(3) / 3007.0580 Da and HexNAc(7)Hex(6)NeuGc(2) / 3008.0532 Da.

Determination of fucose position on the *N-*glycan is an additional difficulty when deciphering the structure of a fucosylated *N-*glycan. Hexose rearrangement is a reaction often found in MS where an internal hexose migrates to a different position within the *N-*glycan, producing “false” *N-*glycan structures [64]. Fucose rearrangement occurs in the gas-phase, resulting in a fucose transfer between two antennae or between core and – most likely – α6-Man-linked antenna (due to its flexibility) [42]. In consequence, this reaction generates misleading fragment ions such as HexNAc_1_Hex_1_Fuc_1_ in the MS^2^ HCD.step spectra of only core fucosylated *N-*glycopeptides. Acs *et al.*, show that core-fucose linkage is robust at high NCE energies. On the other hand, Acs *et al*. and Wuhrer *et al*., also acknowledge that a transfer of fucose from the antenna to the core can be induced, since they observed the Y-ion (peptide-HexNAc_2_Hex_3_Fuc_1_ [M+H]^+^). However, in our study, that Y ion was not used for confirming core fucosylation but primarily the peptide-HexNAc_1_Fuc_1_ [M+H]^+^ Y ion.

Diverse studies reported that core and antenna fucose-linkage stability improves by using NCE 20 instead of higher collisional energies, even though, fragments from fucose rearrangement might still be produced in lower abundance [26,65]. In agreement with these studies, we observed the ambiguous generation of fucose fragment ions, e.g. detection of both peptide+HexNAc_1_Fuc_1_ (Y-ion) and HexNAc_1_Hex_1_Fuc_1_ [M+H]^+^ in the MS^2^ HCD.step spectra of monofucosylated *N-*glycopeptides. In this case, only core fucosylation was granted as it was assumed that the superior stability of core fucosylation would most likely lead to the generation of an antenna-marker ion instead of the opposite rearrangement. However, to describe multiple-fucosylated *N-*glycan structures with a high confidence, it is necessary to check more details. For example, the presence of Y ions, neutral losses and the MS^1^ isotopic pattern distribution using the spectra acquired at low collisional energies, but for a large number of *N-*glycopeptide identifications, manual annotation is not feasible.

Over the last years, special emphasis was given towards describing not only *N-*glycan structural information but also their localization within the protein [17]. Even though glycomic analysis has successfully added on the description of structural groups in *N-*glycans (diLAcNAc unit, sialyl Lewis X, phosphorylation, etc.), only MS allows to precisely determine the original position of such *N-*glycans. With the aim to integrate *N-*glycan structural information using MS, Shen *et al.*, conducted an *N-*glycoproteomic analysis of mouse brain tissue using the new software “StrucGP”. By acquiring and combining two complementary fragmentation energies (HCD.low for glycan and HCD.step for peptide moiety), this software was able to interpret *N-*glycan structural groups on intact *N-*glycopeptides [29]. Similarly, we coupled information from both HCD.step and HCD.low MS/MS analysis, and manually annotated the oxonium marker ions that support the identification of special *N-*glycan structural features and modifications, as previously described by our group [26]. From the observation of fragment ions, *N-*glycan isoforms such as bisecting *N-*glycans and repeated LAcNAc structure were identified. These identifications were searched in GlyConnect to find their corresponding reported information [52,53]. The bisecting *N-*glycans observed in immunoglobulins proteins and plasma protease C1 inhibitor are in agreement with information reported in Glyconnect [52,53]. However, for the rest of the identified *N-*glycopeptides with bisecting GlcNAc no reports were found (**Supplementary Table S15**). Even though the *N-*glycosylation sites harboring LAcNAc structure have been reported in terms of *N-*glycan composition, evidence of sialylated diLacNAc structure was not described before for the *N-*glycosylation sites identified in this work. An interesting disialo-antennary *N-*glycan was observed in the N_253_ of β-2-glycoprotein 1. A similar *N-*glycan is reported for the site N_162_ of this protein [53]. The disialylated-antenna structure found in our study resembles the epitope disialyl Lewis C NeuAcα2-3Galβ1-3(NeuAcα2-6)GlcNAc, reported in α-2-HS-glycoprotein from *bos taurus* [66]. Disialic acid in one antenna was also found by Saraswat *et al.* in ten blood plasma proteins with a linkage between two sialic acids (except for clusterin protein, where the linkage is also to a HexNAc). However, there were no *N-*glycosylation sites in common between Saraswat *et al.* and our study. Only the protein α-1-antichymotrypsin was identified in both studies with a disialylated antennary group in different *N-*glycosites [61]. Sturiale *et al*. compared the *N-*glycomes from a patient and two controls (parents), identifying the NeuAcα2-3Galβ1-3(NeuAcα2-6)GlcNAc epitope in isolated proteins (transferrin, α-1-antitrypsin, Immunoglobulin G and α-1-acid glycoprotein) and serum *N-*glycome [67]. The production of GlcNAc-NeuAc linkage in human *N-*glycans has been studied previously [68,69]. Our study provides evidence that proposes the existence of this glycoepitope in the low-abundant blood plasma *N-*glycoproteome.

Sulfation and phosphorylation are two post-glycosylational modifications that add a negative charge to the *N-*glycan. The *N-*glycopeptides holding this type of glycan modifications (as well as sialylated *N-*glycans) are better detected by methods that enrich anionic molecules [70,71]. Nevertheless, our study showed that, while none of these modified *N-*glycopeptides were detected through the analysis of the untreated human blood plasma, their detection is possible after applying a multi-step sample preparation workflow. *N-*Glycopeptides featuring sulfated and phosphorylated *N-*glycans were detected for proteins with reported concentrations ranging from 4.17·10^8^ down to 1.55·10^5^ pg/mL (PPD, plasma proteome database) [40]. Our results show that sulfated *N-*glycopeptides are more frequently present than phosphorylated *N-*glycopeptides. However, *N-*glycopeptide identifications holding phosphorylated *N-*glycans always showed corresponding oxonium marker ions –which was not the case for sulfated *N-*glycopeptides. It seems that phosphate-containing fragment ions are more stable than sulfate-containing fragment ions. This observation is in agreement with Zhang, T. *et al.*, who found that the abundance of Hex_1_Phospho_1_ [M-H_2_O+H]^+^ oxonium ions was higher than Hex_1_Sulfo_1_ [M+H]^+^ using low fragmentation energy [72]. From the sulfated *N-*glycopeptide identifications observed, only 17% showed the oxonium ion HexNAc_1_Sulfo_1_ [M+H]^+^ within their MS^2^ spectra, probably due to the short lifetime of sulfated fragment ions in MS analysis [43]. Spurious sulfated *N-*glycopeptide identifications might be caused by the assignation of sulfation plus two HexNAc instead of three Hexoses (Hex_3_=486.1585 Da and HexNAc_2_Sulfo_1_=486.1156 Da). Shu *et al*., detected phosphorylated and sulfated *N-*glycans in many blood plasma proteins, including some proteins also found here. In contrast, all the sulfated *N-*glycans identified in their study show hexose sulfation instead of a HexNAc sulfation [59]. Previous studies showed that *N-*glycans holding a sulfated-galactose (like Gal-3-sulfate) do not generate the ion Hex_1_Sulfo_1_ [M+H]^+^; however these sulfated *N-*glycans generate HexNAc_1_Hex_1_Sulfo_1_ [M+H]^+^ (scarcely) and a sulfate neutral loss between the Y ion including galactose and the consecutive sulfated-Gal Y ion [30]. This suggests that some of our sulfated *N-*glycopeptide identifications lacking of sulfated fragment ions might correspond to galactose-sulfated *N-*glycopeptides, however it is more difficult to collect this evidence for all identifications, since it is required to annotate manually the Y ions observed in each HCD.low MS^2^ spectra. Nevertheless, during the manual validation of our data in the HCD.step MS² spectra, no HexNAc_1_Hex_1_Sulfo_1_ [M+H]^+^ oxonium ions were detected but only HexNAc_1_Sulfo_1_ [M+H]^+^. After *de novo* sequencing in HCD.low MS^2^ spectra, the HexNAc sulfation was confirmed in the antennae, most likely corresponding to sulfated-6-GlcNAc (Figure 7). The majority of the sulfated *N-*glycopeptides found here, feature diantennary, mono-or disialylated, and sometimes core-fucosylated *N-*glycans with LacNAc extensions, resembling the LacNAc sulfated glycoepitope: 6-sulfo sialyl Lewis X in its defucosylated form. The *N-*glycopeptide identifications containing confirmed sulfated *N-*glycans derive from glycoproteins involved in blood coagulation, fibrinolysis, hemostasis, inflammatory response, mineral balance, osteogenesis, complement pathway, apoptosis, and innate immunity [73].

LacNAc sulfated *N-*glycans have various implications in immunology. On the one hand, 6-sulfo sialyl Lewis X (with GlcNAc-6-SO4) is a ligand for L-selectin, a cell adhesion molecule for tethering and trafficking of lymphocytes through the peripheral nodes [33]. On the other hand, the structural isomer, 6’-sulfo sialyl Lewis X (with Gal-6-SO4), is a primary ligand of Siglec-8. The cross linkage between 6’-sulfo sialyl Lewis X and Siglec-8, in mast cells, induces histamine and prostanglandin D2, whereas it induces apoptosis in eosinophils [31,32]. A glycomic analysis performed by Yamada *et al.*, on the serum of patients with pancreatic cancer showed that the abundance of sulfated *N-*glycans is increased compared to healthy controls [15]. On the contrary, the study showed that the amount of phosphorylated *N-*glycans remained stable, stressing the importance of reliably discriminating both *N-*glycan modifications in pathophysiology. Sleat *et al.*, detected for first time Man-6-P in high-abundant plasma glycoproteins. They calculated the relative fraction of proteins with phosphorylated *N-*glycans in blood plasma and compared it with the fraction of proteins with phosphorylated *N-*glycans from lysosome, concluding that phosphorylated *N-*glycans in blood plasma proteins exist rather as traces [34]. It is known that Man-6-P is recognized by transporters (Man-6-P receptors) that carry the Man-6-P modified glycoprotein to the lysosomes [34]. This receptor, expressed in all human cells and tissues, is not only present intracellularly in Golgi apparatus and endosomes but also extracellularly, in cell membranes [35]. In the extracellular context, Man-6-P receptors like CD222 are involved in protein internalization, trafficking, lysosomal biogenesis, apoptosis, cell migration, and regulation of cell growth [35]. Overall, this suggests that interactions between mannose-phosphorylated *N-*glycans and binding molecules like CD222 (a receptor from the P-lectin family) are essential in physiology [35,36].

Another *N-*glycan modification not regularly explored in the clinical context is glucuronidation. *N-*glycopeptides containing glucuronic acid were observed after the manual curation of incorrectly assigned *N-*glycopeptide identifications. The incorrect matches might have resulted from the assignation of two NeuAc instead of two HexNAc plus one GlcA residue, which have in total the same atomic composition (2NeuAc=2[C11H17O8N] = C22H34O16N2, and 2HexNAc+1GlcAc = 2[C8H13O5N]+1[C6H8O6] = C22H34O16N2 = 582.1908 Da). Huffman, *et al*. observed glucuronidated *N-*glycans in the blood plasma *N-*glycome of different European populations [16]. Yamada, *et al.*, reported a reduced relative abundance of glucuronidated *N-*glycans from the serum of patients with pancreatic cancer, compared to healthy controls [15]. Sulfated glucuronic acid is more common in brain *N-*glycans as a key component of the glycoepitope HNK-1 (SO4-3GlcAβ1-3Galβ1-4GlcNAc). HNK-1 influences neuronal functions like adhesion, cell recognition, migration, preferential motor-re-innervation, synaptic plasticity, and post-trauma regeneration in the peripheral and central nervous systems [74,75]. Laminins and cadherin-2 are HNK-1 binding proteins [75]. Nevertheless, it is unclear if this is relevant to the biological role of the non-sulfated HNK-1.

Two new *N-*glycan building blocks with the masses 245.0524 and 259.0672 Da attached to complex di-antennary monosialylated *N-*glycans were identified. During our literature search, we could not find any reported residues that resemble such *N-*glycan building blocks in human. A limitation for this observation is the lack of an orthogonal method to describe the molecule structure. Interestingly, both rare-*N-*glycan building blocks were found in prothrombin (glycosylation site N_121_). This *N-*glycosylation site corresponds to prothrombin fragment region 1 (also called kringle-1), which is important for calcium-mediated membrane-surface binding [76].

Bioinformatics tools able to provide a site-specific *N-*glycan description from the analysis of intact glycopeptides emerged 12 years ago [77]. Since then, software design has moved towards improving gPSM quality (pGlyco 2.0), visualization of results (pGlyco 2.0, glyXtoolMS, GPSeeker), and annotation of *N-*glycopeptide structural features (GPSeeker, glyXtoolMS, StrucGP) [21,29,78,79]. Nonetheless, challenges that prevent the accurate analysis of an *N-*glycoproteome to date are the lack of comprehensiveness, ambiguity on peptide composition or glycan structure, missing structural information, and false-positives [17]. Consequently, the detection of *N-*glycopeptides potentially useful as biomarkers or for therapy is hampered. Even though manual validation is an alternative for detecting pitfalls after a glycoproteomic search, it has discouraging aspects, such as the high time requirement and effort needed, plus the bias caused by each data reviewer. For future developments, it would be appropriate for *N-*glycoproteomic software to include an *N-*glycan diagnostic step plus the integration of *N-*glycan structural features during glycopeptide spectra matching. Missing *N-*glycan compositions, lack of information, or incorrect parameters are barriers against fully explaining an *N-*glycoproteomic spectra input. Our blood plasma *N-*glycoproteomic analysis aimed to find *N-*glycopeptides eclipsed between the obscure niche of the low-abundant *N-*glycoproteome and the MS^2^ spectra neglected due to insufficient search parameters. After manual validation, we acknowledged by the categories “Uncertain change *N-*glycopeptide” and “Uncertain glycopeptide”, that 51% of total gPSMs are still unexplained, for a greater part, due to the low quality of the fragment ion spectra. Nonetheless, we were able to identify several previously unknown and rare *N-*glycans plus various atypical *N-*glycan structures that are potentially useful for the design of biotherapeutics, clinical diagnostic (biomarker discovery), or exploration of *N-*glycan-protein interactions and functions.

## 5. Conclusions

Glycoproteomic sample preparation, LC-MS measurement, and data analysis software have further evolved in the last years. Nonetheless, two major limitations still need to be addressed: structural glycan elucidation and precision of gPSMs. Blood plasma analysis is a common and challenging task for exploring the potential of new *N-*glycoproteomic search strategies. Here we developed and applied a sample preparation workflow, which comprises fractionation of HAP-depleted blood plasma and glycopeptide enrichment followed by two LC-MS/MS measurements using the fragmentation energies HCD.step and HCD.low. The workflow enabled the detection of glycoproteins in a concentration range from 10^9^ down to 10^1^pg/mL, thus expanding the detection range by five orders of magnitude compared to the direct analysis of blood plasma. Validation and curation of the *N-*glycoproteomic search is based on a gPSM decision tree to critically assess mass spectral evidence on the peptide and *N-*glycan level. This approach allowed to reliably elucidate structural glycan features and to identify rare *N-*glycan compositions. This included, for instance, the presence of glucuronic acid and two rare *N-*glycan building blocks (245.0524 and 259.0672 Da). Furthermore, an atypical disialylated-antenna *N-*glycan structure containing a HexNAc-NeuAc linkage that resembles the Lewis C epitope NeuAcα2-3Galβ1-3(NeuAcα2-6)GlcNAc was primarily identified based on the presence of the oxonium ion HexNAc_1_NeuAc_1_ [M+H]^+^. Interestingly, without HCD.low spectra annotation, the NeuAc in those *N-*glycopeptides were erroneously assumed to be spread on different antennae. Other structural groups hard to discriminate, such as antenna versus core fucose, diLAcNAc versus multi-antennae, bisecting GlcNAc, and phosphorylation versus sulfation, were also reliably confirmed. In contrast to previous reports, no hexose sulfation but only HexNAc sulfation was detected with high certainty through oxonium marker ions. Making use of manual validation, we contribute to uncover problems, pitfalls, and opportunities in the field of glycoproteomics research applied to biomarker discovery, biotherapeutic products, and biochemistry. In the future, we aim to transfer the key features of our data analysis workflow to bioinformatics tools, in order to establish an efficient option to obtain an accurate picture of the *N-*glycoproteome of any sample.

## Supplementary Materials

**Figure S1:** Oxonium ion filter applied to identify oxonium ions present in sulfated glycopeptides; **Figure S2**: Oxonium ion filters applied to identify oxonium ions present in phosphorylated glycopeptides; **Figure S3**: Total *N-*glycopeptides identified in blood plasma after in-depth *N-*glycoproteomic analysis; **Figure S4**: Example of a gPSM with the features assigned for the validation category named as “True Match with Outstanding Evidence”; **Figure S5**: Example of a gPSM with the features assigned for the validation category named as “True Match with Evidence”; **Figure S6**: Example of a gPSM with the features assigned for the validation category named as “True Match with Evidence and Alternatives”; **Figure S7**: Example of a gPSM with the features assigned for the validation category named as “True Match with Evidence Relative to matches from the same peptide”; **Figure S8**: Example of a gPSM with the features assigned for the validation category named as “Quasi-true Match Double Site”; **Figure S9**: Example of a gPSM with the features assigned for the validation category named as “Quasi-true Match change glycan”; **Figure S10**: Example of a gPSM with the features assigned for the validation category named as “Quasi-true corrected Match”; **Figure S11**: Example of a gPSM with the features assigned for the validation category named as “Uncertain-Change glycopeptide match”; **Figure S12**: Example of a gPSM with the features assigned for the validation category named as “Uncertain glycopeptide”; **Figure S13**: Example of a gPSM with the features assigned for the validation category named as “False *O-*glycan”; **Figure S14**: Example of a gPSM with the features assigned for the validation category named as “False”; **Figure S15**: Examples of gPSM including multiple fucose in a complex *N-*glycan with a potential error in the assignation of the monoisotopic peak; **Figure S16**: Byonic software annotation of fragment ions released from sulfated *N-*glycopeptide; **Figure S17**: Byonic software annotation of fragment ions released from phosphorylated *N-*glycopeptide; **Figure S18**: Searching for rare *N-*glycopeptides in untreated blood plasma and top 14-HAP depleted sample; **Figure S19**: ESI-MS/MS prediction for two molecules proposed as the *N-*glycan building blocks of the rare *N-*glycopeptide identifications observed; **Figure S20**: Byonic software annotation of HCD.step fragment ion spectra containing rare *N-*glycans found in N121 from prothrombin; **Figure S21**: Byonic software annotation of HCD.step fragment ion spectra from *N-*glycopeptides with disialo-antennary structure confirming HexNAc-NeuAc linkage.

**Table S1**: Proteomic analysis of HILIC-fractions from untreated blood plasma sample; **Table S2**: Proteomic analysis of HILIC-fractions of top 14-HAP depleted sample; **Table S3**: Proteomic analysis of HILIC-fractions of top 14-HAP depleted sample plus fractionation by protein size (applying developed workflow); **Table S4**: Micro-heterogeneity of *N-*glycosylation observed in Zinc-alpha-2-glycoprotein (P25311) after each step of the sample preparation workflow; **Table S5**: *N-*glycoproteomic analysis of untreated blood plasma sample; **Table S6**: *N-*glycoproteomic analysis of top 14-HAP depleted sample; **Table S7**: *N-*glycoproteomic analysis of top 14-HAP depleted sample plus fractionation by protein size (HCD.step-HPB list); **Table S8**: *N-*glycoproteomic analysis of top 14-HAP depleted sample plus fractionation by protein size (HCD.low-HPB list); **Table S9**: Classified fucosylated *N-*glycan structures in *N-*glycopeptide identifications validated as true in the HCD.step-HBP list; **Table S10**: Sulfated *N-*glycopeptide identifications validated as true in the HCD.step-HBP list; **Table S11**: Phosphorylated *N-*glycopeptide identifications validated as true in the HCD.step-HBP list; **Table S12**: Glucuronidated *N-*glycopeptide identifications validated as true in the HCD.step-HBP list; **Table S13**: Rare *N-*glycopeptide identifications validated as true in the HCD.step-HBP list; **Table S14**: Cross referenced rare-compositions and modified *N-*glycans;

**Table S15**: Cross referenced especial *N-*glycan structural features.

## Author Approvals

All the authors approved the publication of this work. This work has not been accepted or published elsewhere.

## Author Contributions

Conceptualization, M.H., F.J.Z.B. and E.R.; methodology, M.H., F.J.Z.B. and E.R.; validation, M.H., F.J.Z.B. and E.R.; formal analysis, F.J.Z.B.; investigation, F.J.Z.B.; data curation, F.J.Z.B.; writing—original draft preparation, F.J.Z.B.; writing—review and editing, U.R., M.H. and E.R.; visualization, F.J.Z.B.; supervision, U.R., M.H. and E.R.; project administration, U.R. and E.R.; funding acquisition, U.R. and E.R. All authors have read and agreed to the published version of the manuscript.

## Data Availability Statement

All the raw data and search results produced in this study were deposited to the ProteomeXChange Consortium, dataset PXD042039 (https://proteomecentral.proteomexchange.org/cgi/GetDataset?ID=PXD042039), through MassIVE MSV000091870 (ftp://massive.ucsd.edu/MSV000091870/).

## Acknowledgments

The authors gratefully thank to Iva Budimir for creating visProteomics R package. Moreover, we thank Barbara Koehler and Lisa Fichtmueller for technical support.

