## Supplementary Figures for "New avenues for human blood plasma biomarker discovery via improved in-depth analysis of the low-abundant *N-*glycoproteome"

### List of Supplementary Figures


1. **Oxonium ion filter applied to identify oxonium ions present in sulfated glycopeptides.** Filter applied in Thermo Xcalibur Qual Browser software.
2. **Oxonium ion filters applied to identify oxonium ions present in phosphorylated glycopeptides.** Filter applied in Thermo Xcalibur Qual Browser software.
3. **Total *N*-glycopeptides identified in blood plasma after in-depth *N*-glycoproteomic analysis.** (**A**) Distribution of unique *N*-glycopeptides according to *N*-glycan attached. The x-axis in full range. (**B**) Closer overview of the distribution of glycopeptides bearing rare *N*-glycans.
4. **Example of a gPSM with the features assigned for the validation category named as "True Match with Outstanding Evidence".** (**A**) Validation category color code (**B**) Fragment ion spectrum annotated by Byonic software (**C**) Features description (**D**) Isotopic pattern.
5. **Example of a gPSM with the features assigned for the validation category named as "True Match with Evidence".** (**A**) Validation category color code (**B**) Fragment ion spectrum annotated by Byonic software (**C**) Features description (**D**) Isotopic pattern.
6. **Example of a gPSM with the features assigned for the validation category named as "True Match with Evidence and Alternatives".** (**A**) Validation category color code (**B**) Fragment ion spectrum annotated by Byonic software (**C**) Features description (**D**) Isotopic pattern.
7. **Example of a gPSM with the features assigned for the validation category named as "True Match with Evidence Relative to matches from the same peptide".** (**A**) Validation category color code (**B**) Fragment ion spectra annotated by Byonic software (**C**) Features description (**D**) Isotopic pattern.
8. **Example of a gPSM with the features assigned for the validation category named as "Quasi-true Match Double Site".** (**A**) Validation category color code (**B**) Fragment ion spectrum annotated by Byonic software (**C**) Features description (**D**) Isotopic pattern.
9. **Example of a gPSM with the features assigned for the validation category named as "Quasi-true Match Change Glycan".** (**A**) Validation category color code (**B**) Fragment ion spectrum annotated by Byonic software (**C**) Features description (**D**) Isotopic pattern.
10. **Example of a gPSM with the features assigned for the validation category named as "Quasi-true corrected Match".** (**A**) Validation category color code (**B**) Fragment ion spectra annotated by Byonic software (**C**) Features description (**D**) Isotopic pattern.
11. **Example of a gPSM with the features assigned for the validation category named as "Uncertain-Change glycopeptide match".** (**A**) Validation category color code (**B**) Fragment ion spectrum annotated by Byonic software (**C**) Features description (**D**) Isotopic pattern.
12. **Example of a gPSM with the features assigned for the validation category named as "Uncertain glycopeptide".** (**A**) Validation category color code (**B**) Fragment ion spectrum annotated by Byonic software (**C**) Features description (**D**) Isotopic pattern.
13. **Example of a gPSM with the features assigned for the validation category named as "False O-glycan".** (**A**) Validation category color code (**B**) Fragment ion spectrum annotated by Byonic software (**C**) Features description (**D**) Isotopic pattern.

### List of Supplementary Figures


14. **Example of a gPSM with the features assigned for the validation category named as "False".** (**A**) Validation category color code (**B**) Fragment ion spectrum annotated by Byonic software (**C**) Features description (**D**) Isotopic pattern.
15. **(1-2) Examples of gPSM including multiple fucose in a complex *N*-glycan with a potential error in the assignation of the monoisotopic peak.** (**A**) $MS^1$ Isotopic distributions. Monoisotopic peaks selected by Byonic software (red). (**B**) ) $MS^2$ HCD.low annotation proposed by the software assigning an incorrect *N*-glycan. Correct precursor ion highlighted in green.
16. **Byonic software annotation of fragment ions released from sulfated N-glycopeptide.** (**A**) HCD.low fragment ion spectrum (**B**) HCD.step fragment ion spectrum.
17. **Byonic  software annotation of fragment ions released from phosphorylated *N*-glycopeptide.** (**A**) HCD.low fragment ion spectrum (**B**) HCD.step fragment ion spectrum.
18. **Searching for rare *N*-glycopeptides in untreated blood plasma and top 14-HAP depleted sample.** (**A**) LC-MS/MS-HCD.step analysis of glycopeptide enriched blood plasma. No evidence of sulfated, phosphorylated or glucuronidated glycopeptide-derived $MS^2$ spectra. The signals observed in the glucuronidation marker ion-filter do not derive from glycopeptides. (**B**) LC-MS/MS-HCD.step analysis of glycopeptide enriched of top 14-HAP depleted blood plasma. Evidence of sulfated, phosphorylated or glucuronidated glycopeptide-derived $MS^2$ spectra. (**C**) Top 14-HAP depleted blood plasma Byonic search identified the *N*-glycopeptides holding sulfated, phosphorylated and glucuronidated *N*-glycans. The layout-filters containing glycan marker ions was created in Thermo Xcalibur Qual Browser (Version 2.2, Thermo Scientific) software.
19. **ESI-MS/MS prediction for two molecules proposed as the *N*-glycan building blocks of the rare *N*-glycopeptide identifications observed.** (**A**) Predicted fragmentation of molecule with the chemical composition C9H13NO8 suggested for *N*-glycan building block 245 Da. The molecule found in PubChem (CID 59861591) is shown below the spectra. (**B**) Predicted fragmentation of molecule with the chemical composition C10H15NO8 suggested for *N*-glycan building block 259 Da. The molecule found in PubChem (CID 59173067) is shown below the spectra. (**C**) Predicted fragmentation of Neu5Ac; structure shown below the spectra (PubChemCID 439197). Red triangles indicate the fragment ions observed in the fragment ion spectra containing the rare *N*-glycans and sialic acid. Predicted spectra computed by CFM-ID (https://cfmid.wishartlab.com/predict)  for two collisional energies (10 eV and 20 eV) in positive ion mode including [M+H]+ as adduct. Molecules drawn in PubChem Sketcher [47-51].
20. **Byonic  software annotation of HCD.step fragment ion spectra containing rare *N*-glycans found in N121 from prothrombin (UniProt P00734).** (**A**) N-glycopeptide annotation by the software. (**B**) $MS^1$ Isotopic distribution of precursor ions observed through HCD.low acquisition.
21. **Byonic software annotation of HCD.step fragment ion spectra from *N*-glycopeptides with disialo-antennary structure confirming HexNAc-NeuAc linkage.**

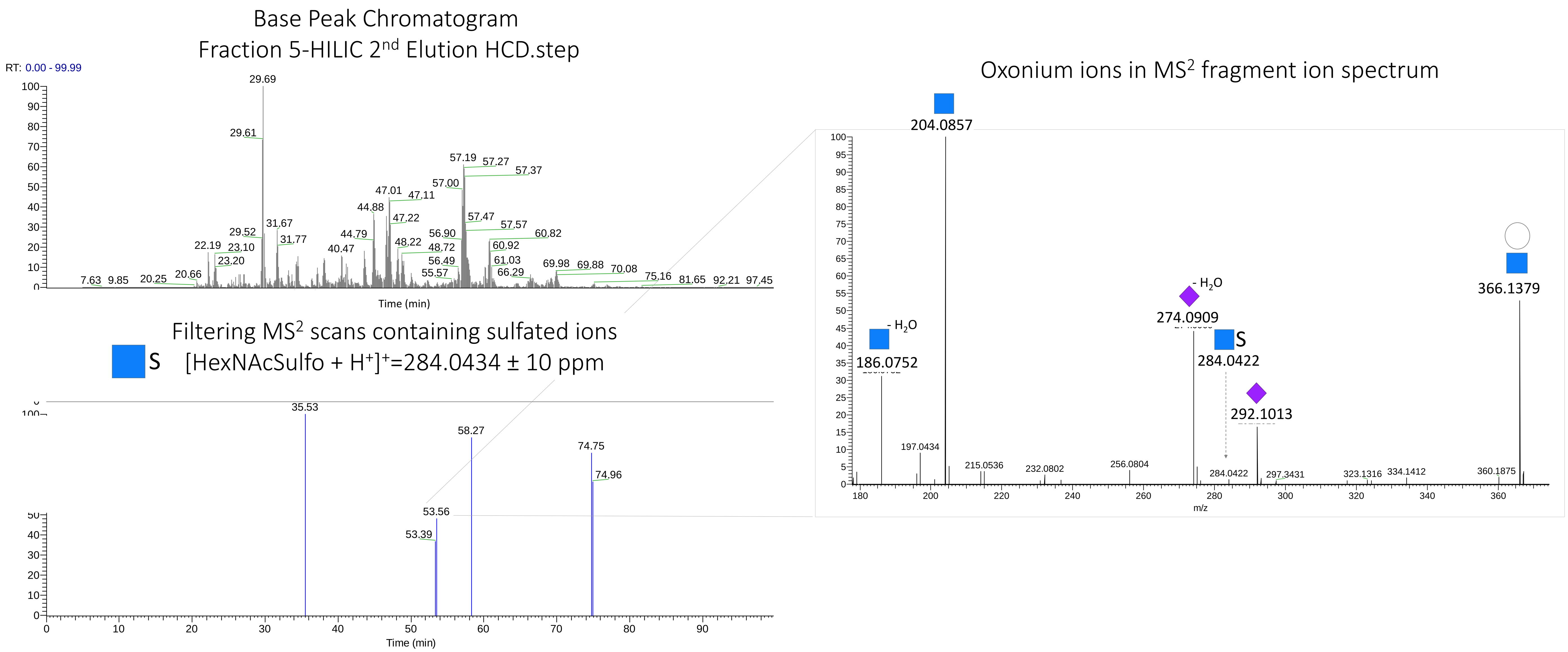


**Supplementary Figure S1. Oxonium ion filter applied to identify oxonium ions present in sulfated glycopeptides.** Filter applied in Thermo Xcalibur Qual Browser software.

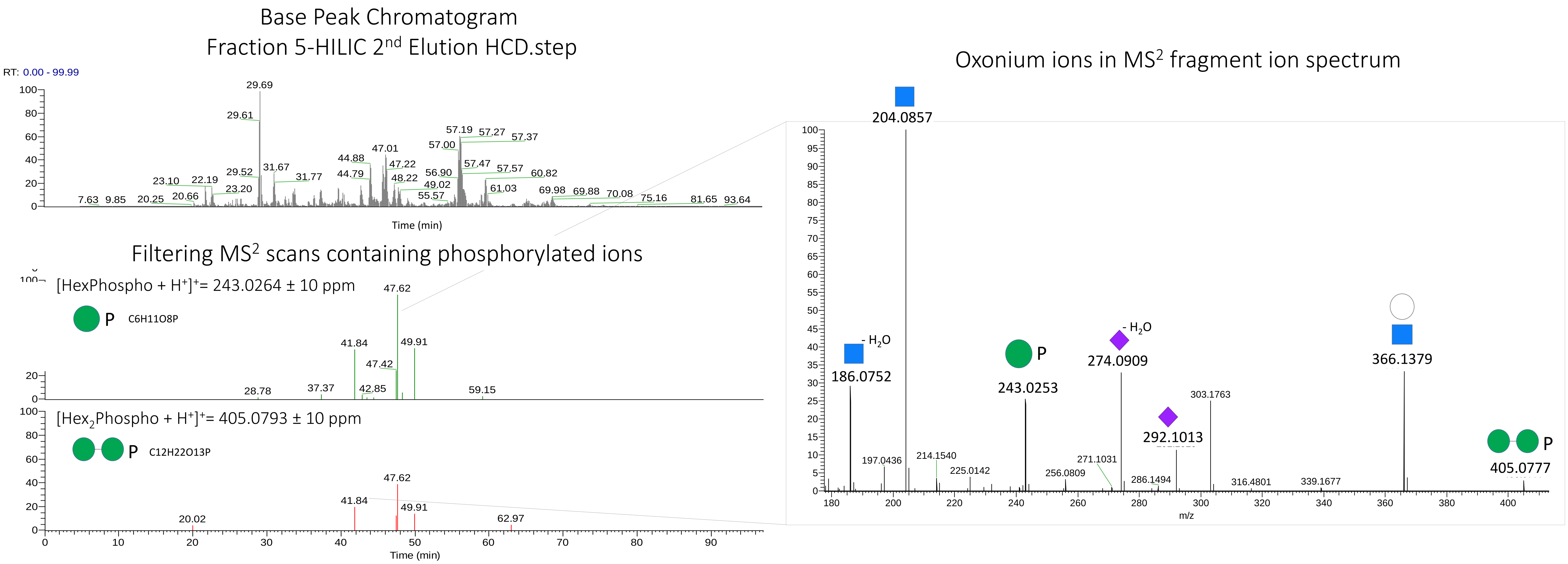


**Supplementary Figure S2. Oxonium ion filters applied to identify oxonium ions present in phosphorylated glycopeptides.** Filter applied in Thermo Xcalibur Qual Browser software.

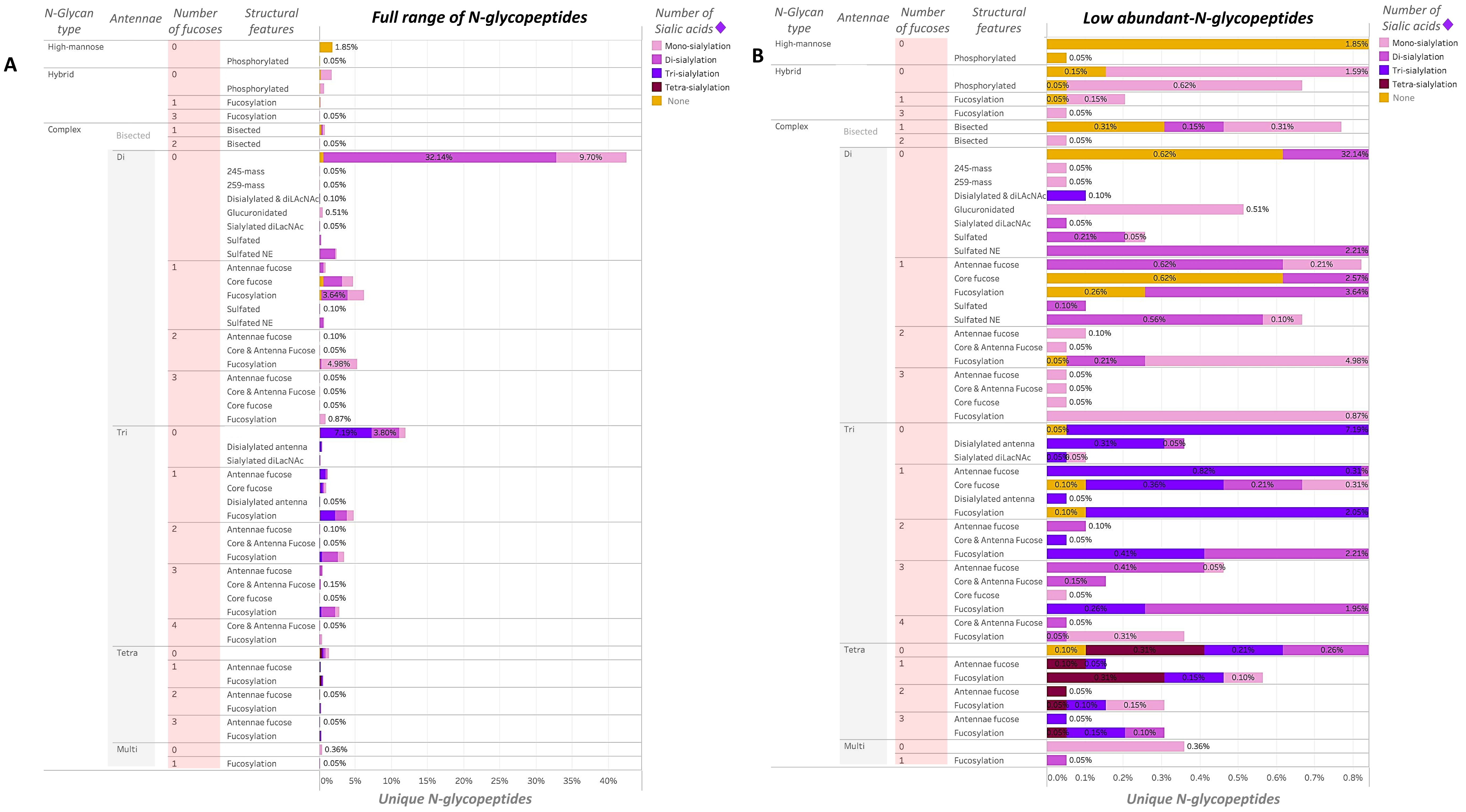


**Supplementary figure S3. Total *N*-glycopeptides identified in blood plasma after in-depth *N*-glycoproteomic analysis.** (**A**) Distribution of unique *N*-glycopeptides according to *N*-glycan attached. The x-axis in full range. (**B**) Closer overview of the distribution of glycopeptides bearing modified or rare *N*-glycans.

### True Match with Outstanding Evidence

A

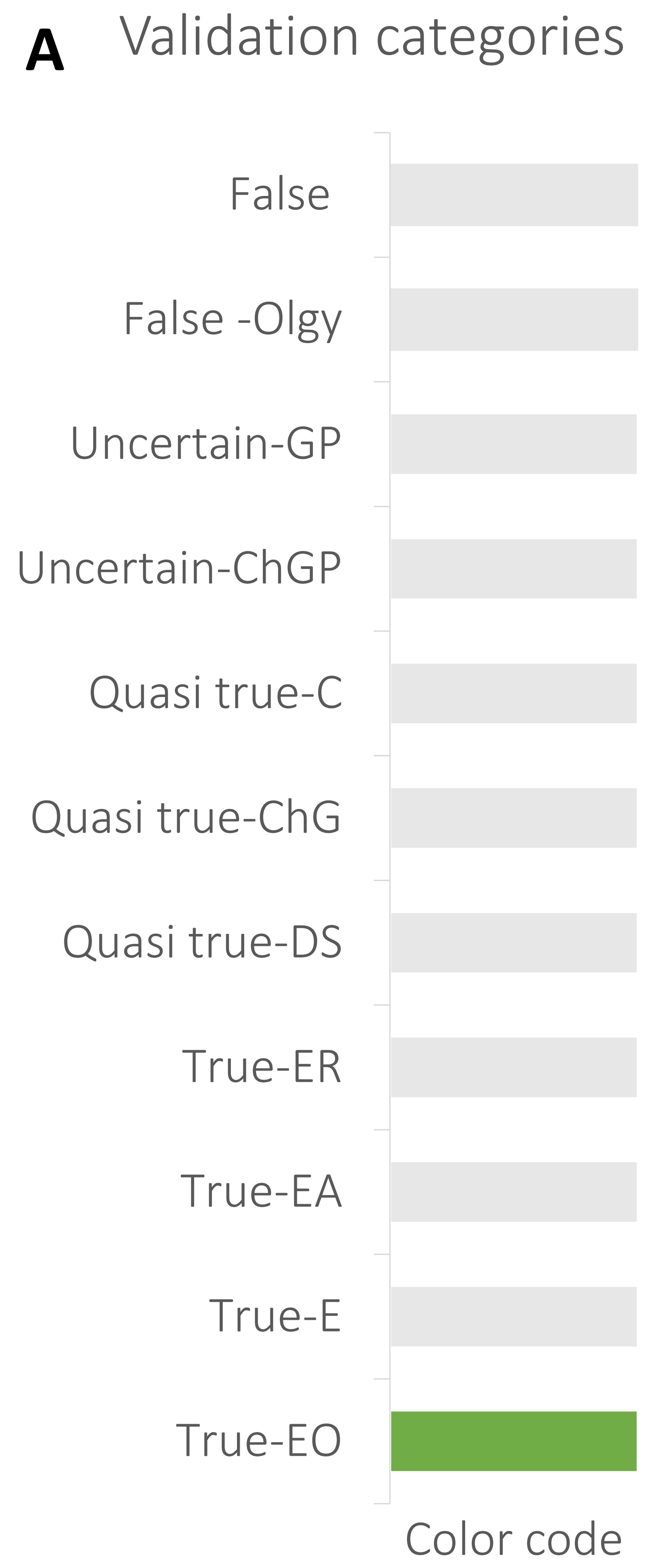


B

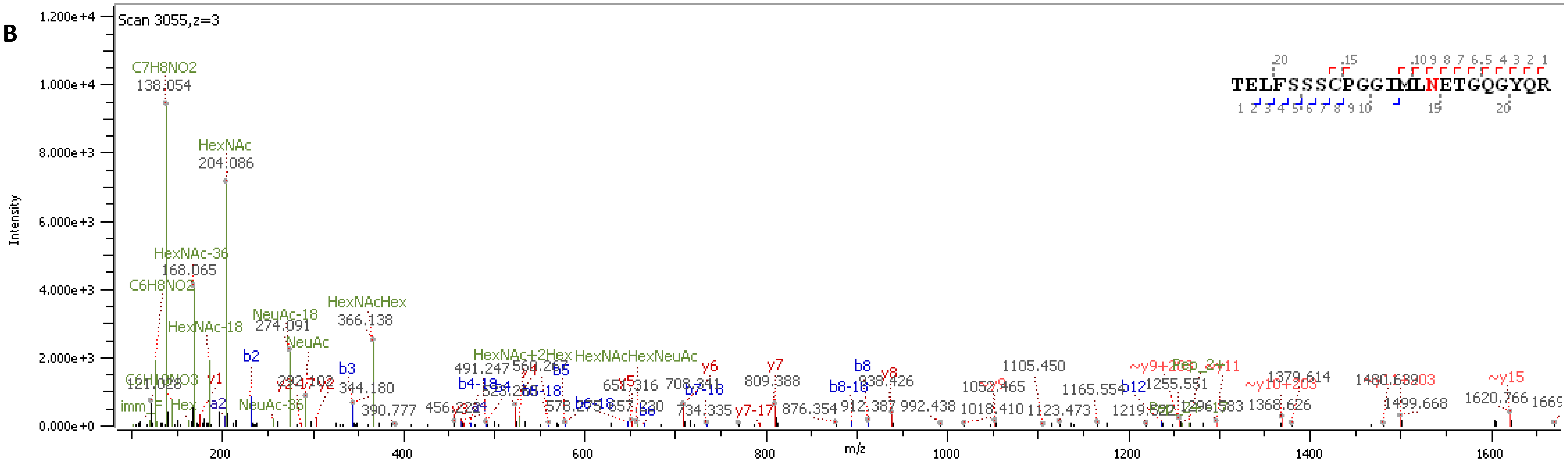


C

| | |
|---|---|
| Main validation category features | • Glycopeptide fragment ions<br>• High number of peptide fragment (b and y) ions |
| Software identification | • Protein: Apolipoprotein M<br>• Glycan: HexNAc(3)Hex(5)NeuAc(1) |

D

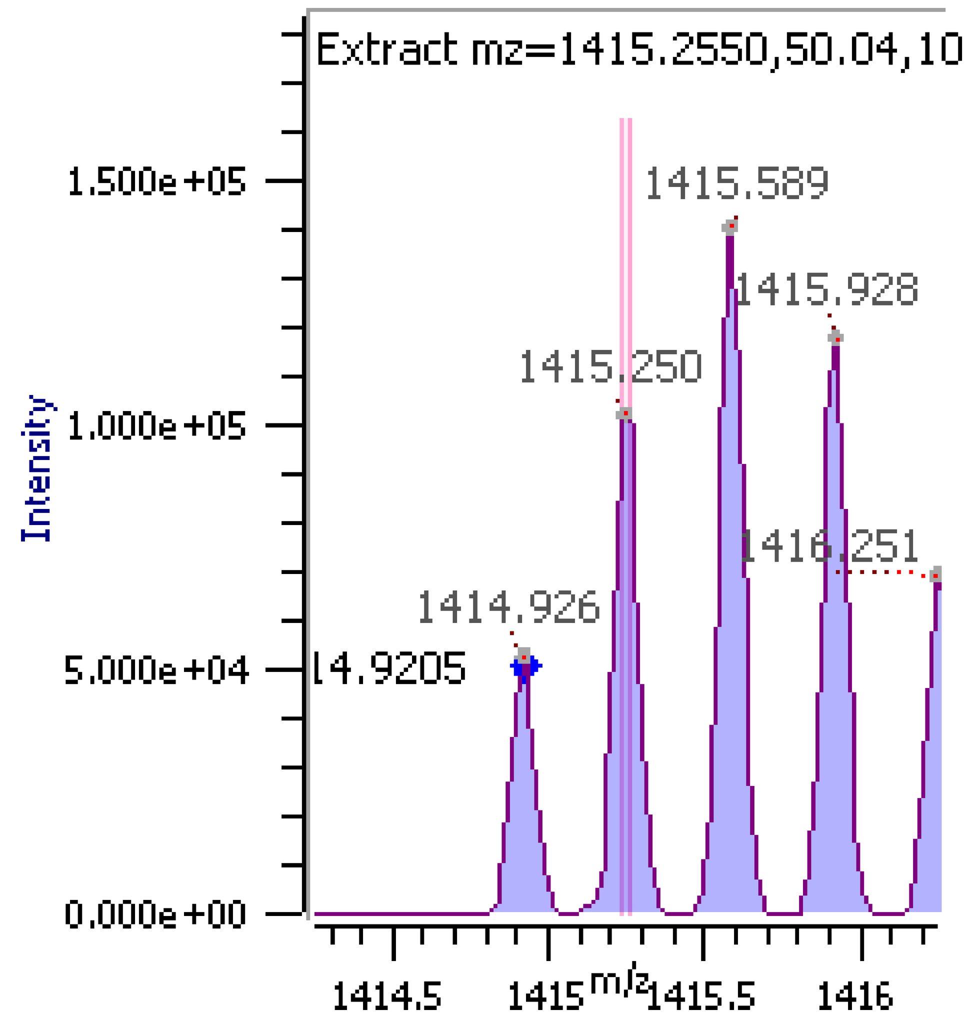


**Supplementary figure S4. Example of a gPSM with the features assigned for the validation category named as "True Match with Outstanding Evidence".** (**A**) Validation category color code (**B**) Fragment ion spectrum annotated by Byonic software (**C**) Features description (**D**) Isotopic pattern.

### True Match with Evidence

**A**

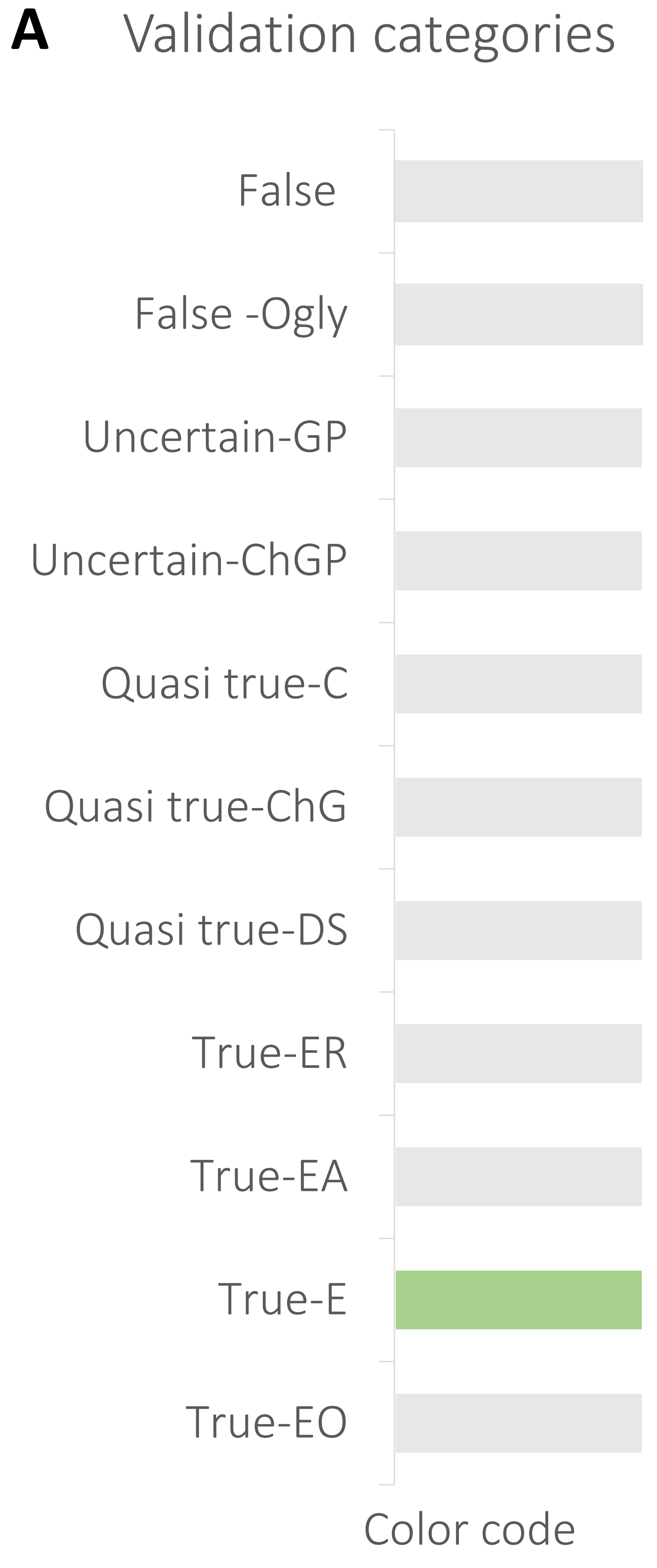


**B**

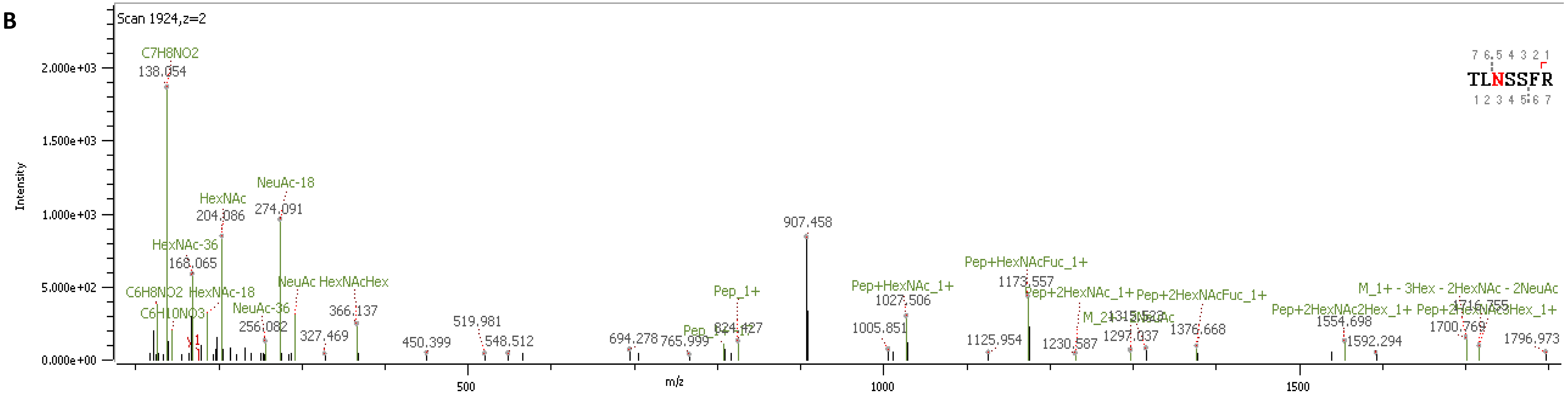


**C**

| | |
|---|---|
| Main validation category features | • Glycopeptide fragment ions<br>• Few or none peptide fragment ions (b and y) |
| Software identification | • Protein: FH2 domain-containing protein 1<br>• Glycan: HexNAc(4)Hex(5)Fuc(1)NeuAc(2 |

**D**

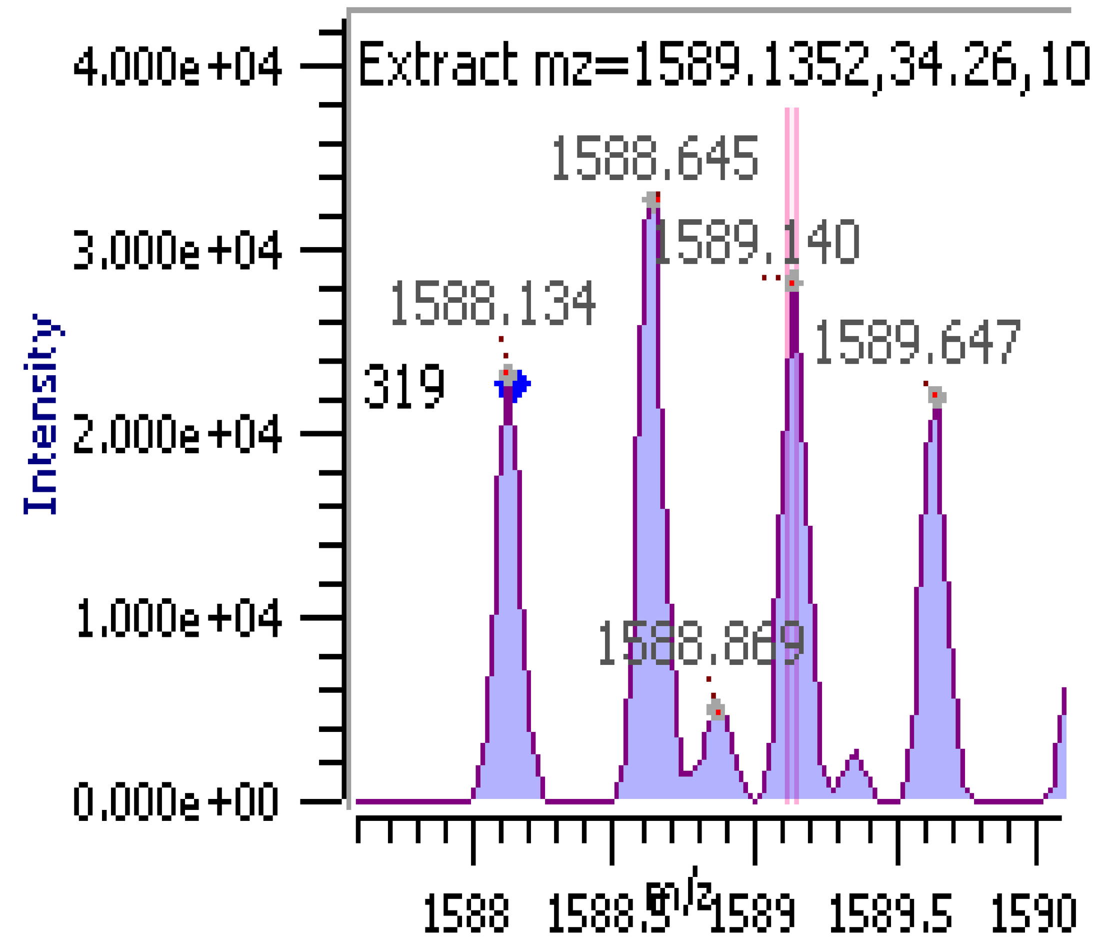


**Supplementary figure S5. Example of a gPSM with the features assigned for the validation category named as "True Match with Evidence".**
(**A**) Validation category color code (**B**) Fragment ion spectrum annotated by Byonic software (**C**) Features description (**D**) Isotopic pattern.

### True Match with Evidence and Alternatives

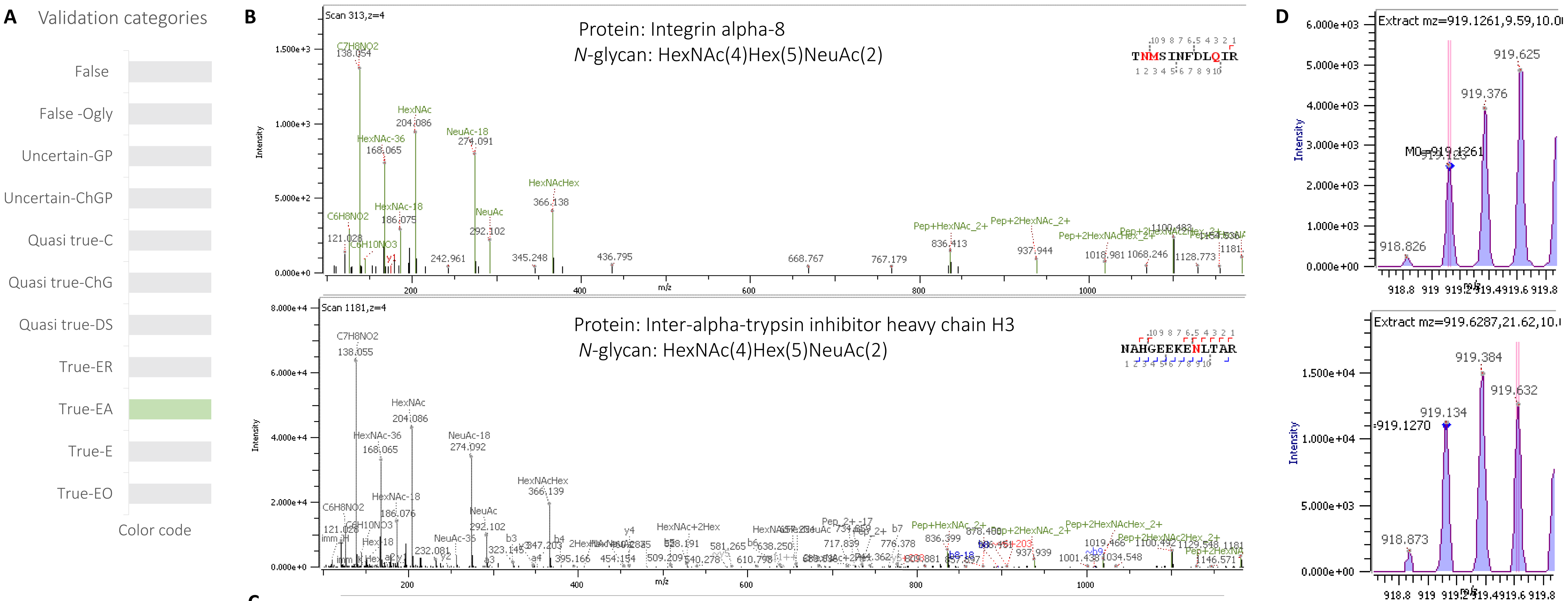


**C**

| Main validation category features | |
|---|---|
| | • Glycopeptide fragment ions<br>• Peptide fragment ions (b and y)<br>• Other glycoprotein(s) match some fragment ions |

**Supplementary figure S6. Example of a gPSM with the features assigned for the validation category named as "True Match with Evidence and Alternatives".** (**A**) Validation category color code (**B**) Fragment ion spectrum annotated by Byonic software (**C**) Features description (**D**) Isotopic pattern.

### True Match with Evidence Relative to matches from the same peptide

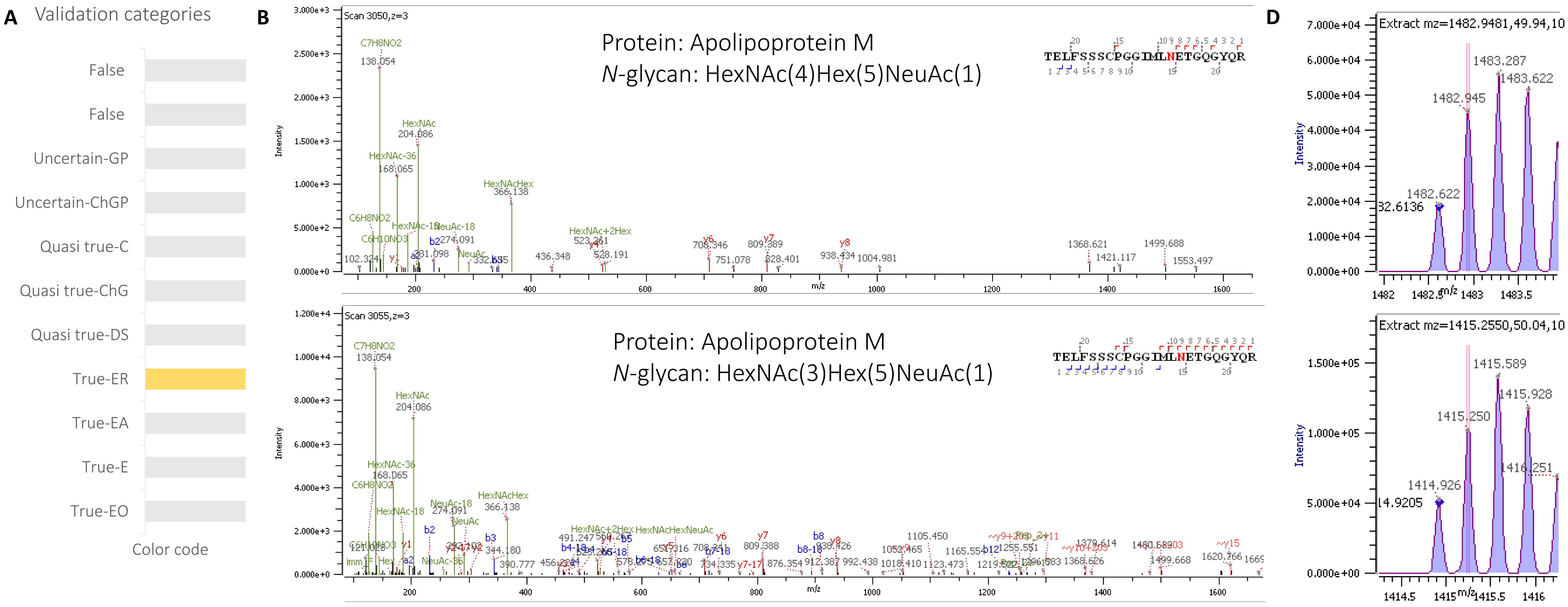


C

| Main validation category features | |
|---|---|
| | • No glycopeptide fragment ions<br>• Peptide fragment ions (b and y)<br>• Other matches from the same peptide |

**Supplementary figure S7. Example of a gPSM with the features assigned for the validation category named as "True Match with Evidence Relative to matches from the same peptide".** (**A**) Validation category color code (**B**) Fragment ion spectra annotated by Byonic software (**C**) Features description (**D**) Isotopic pattern.

### Quasi-true Match Double Site

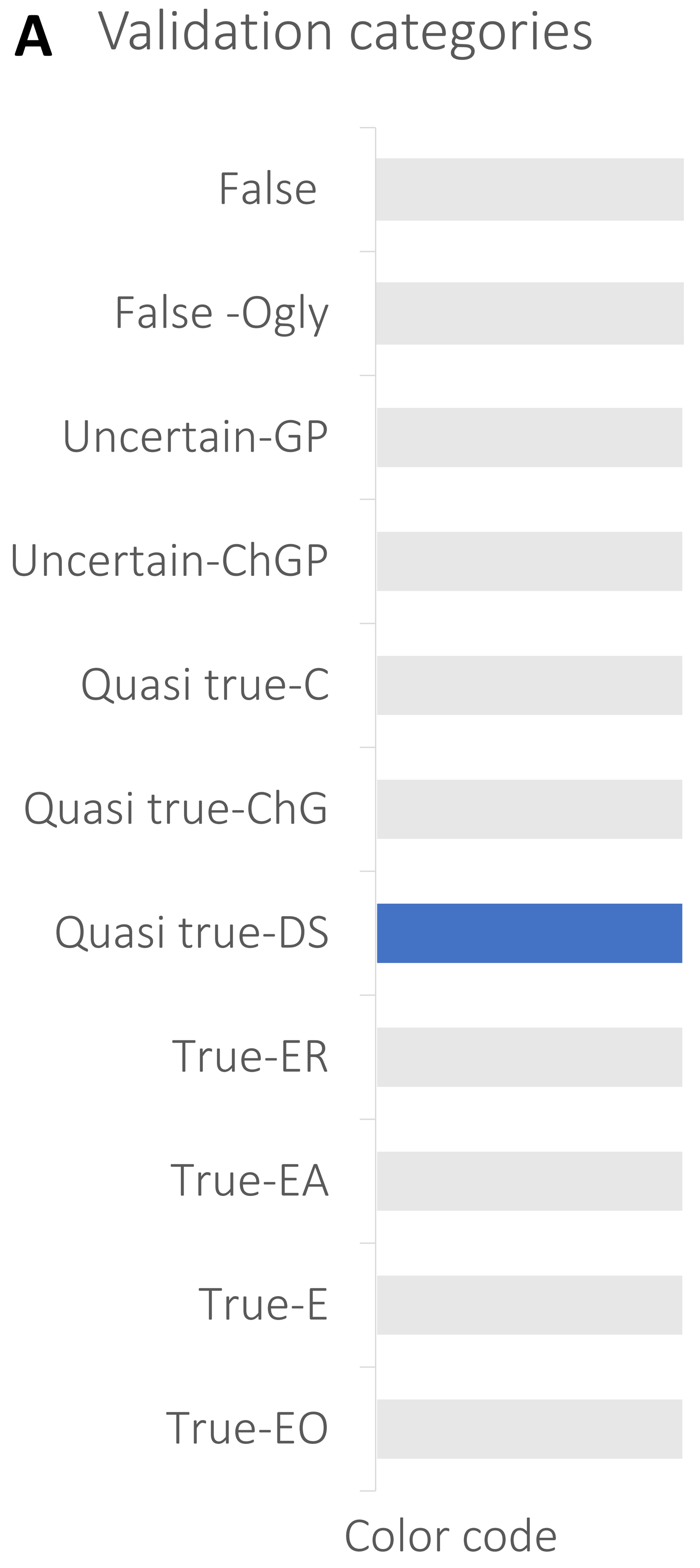


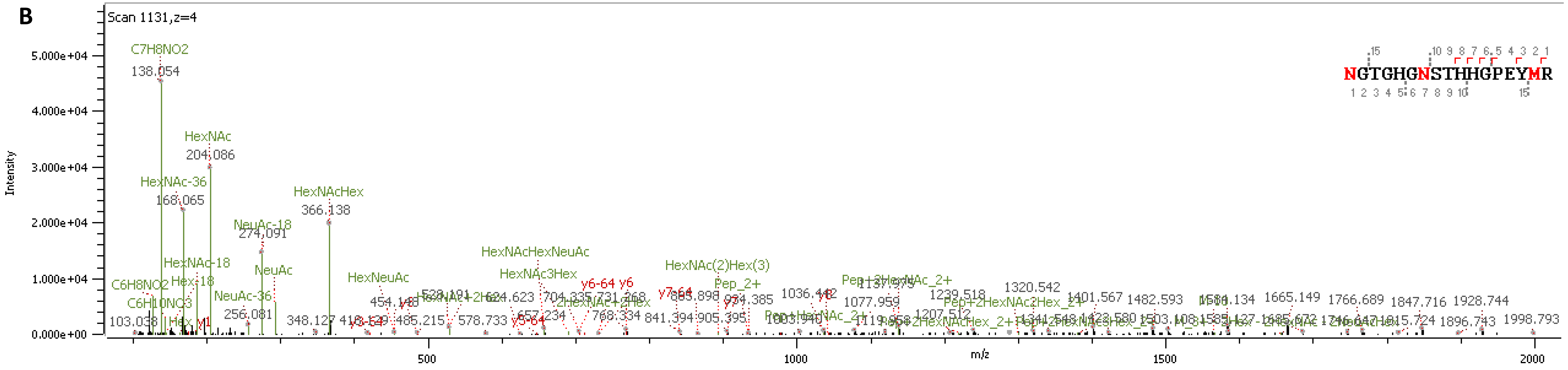


C

| | |
|---|---|
| Main validation category features | • Two *N*-glycosylation sites within the peptide<br>• Two *N*-glycans assigned. Not possible to verify the composition of each *N*-glycan |
| Software identification | **Protein:** Hemopexin<br>***N*-glycan:** HexNAc(2)Hex(3),HexNAc(6)Hex(7)NeuAc(4) |

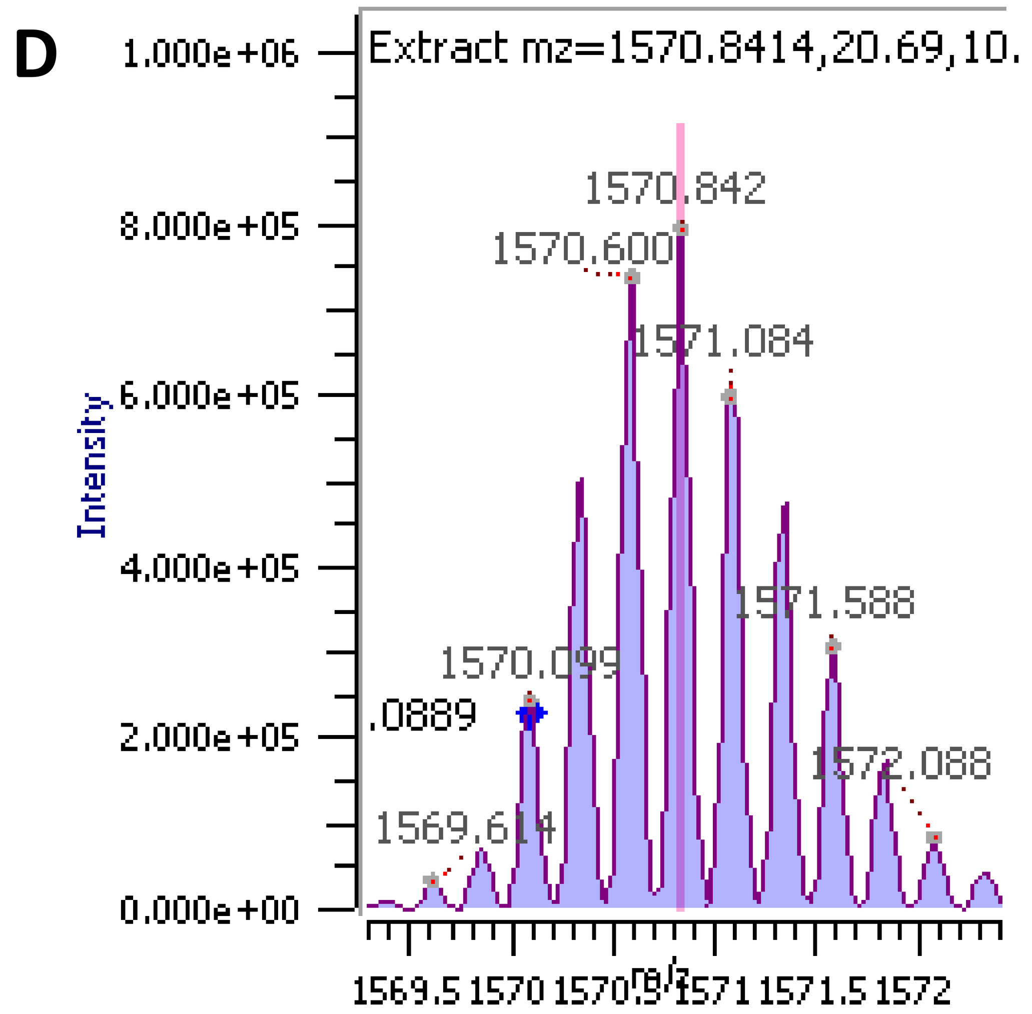


**Supplementary figure S8. Example of a gPSM with the features assigned for the validation category named as "Quasi-true Match Double Site".** (**A**) Validation category color code (**B**) Fragment ion spectrum annotated by Byonic software (**C**) Features description (**D**) Isotopic pattern.

### Quasi-true Match change glycan

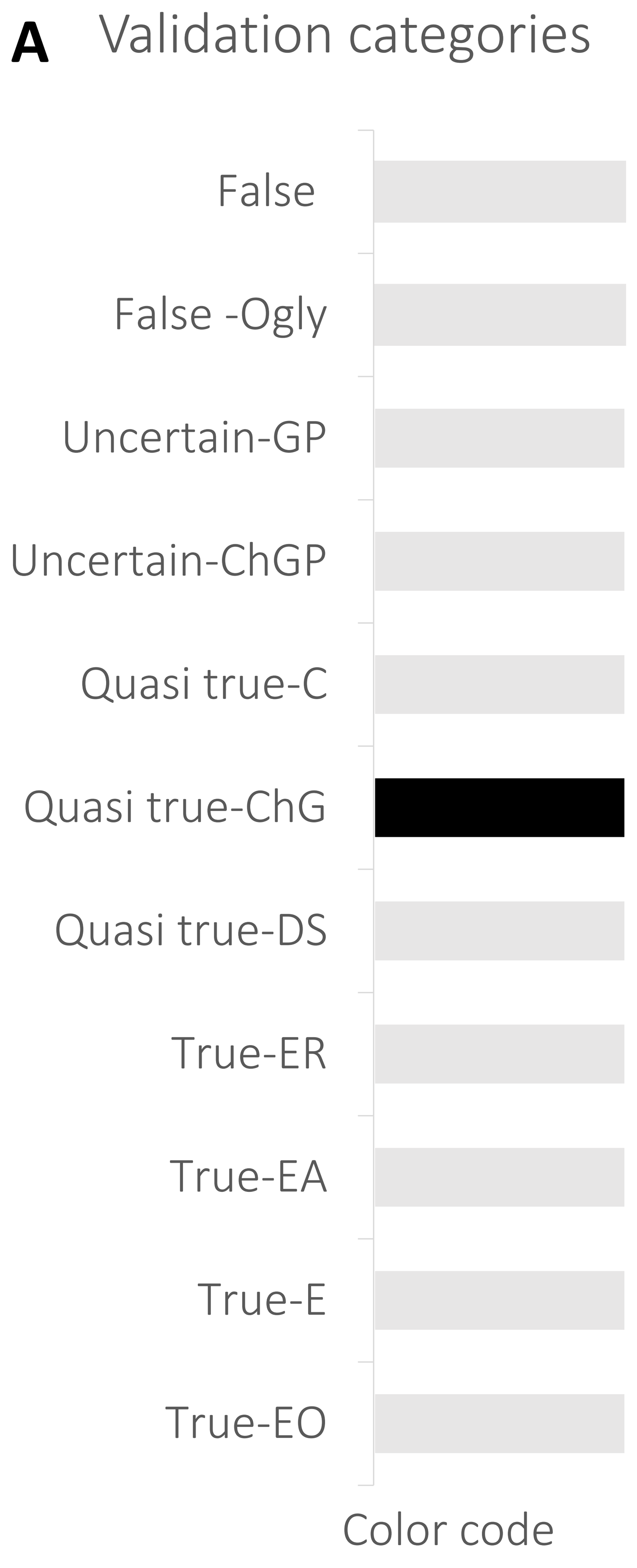


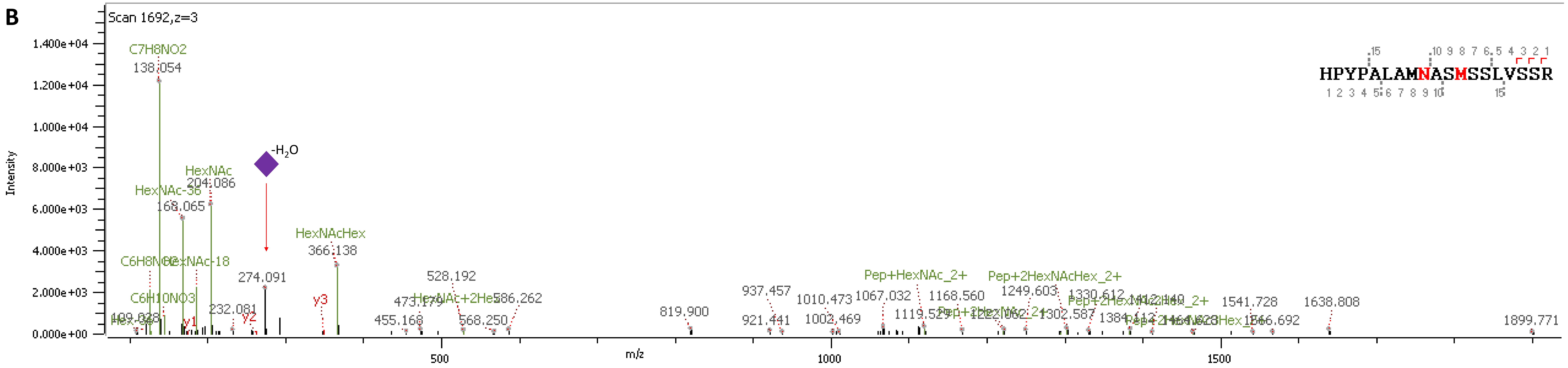


**C**

| | |
|---|---|
| Main validation category features | • Glycopeptide fragment ions<br>• High number of peptide fragment (b and y) ions<br>• Inconsistent evidence in regard to oxonium ions |
| Software identification | • Protein: Transcription factor 7-like 1<br>• *N*-glycan: HexNAc(4)Hex(5)Fuc(2), 1914.6974 Da |
| Expected glycan composition: | • HexNAc(4)Hex(5)NeuAc(1), 1913.6770 Da<br>• No fucose ion, missed NeuAc ion |

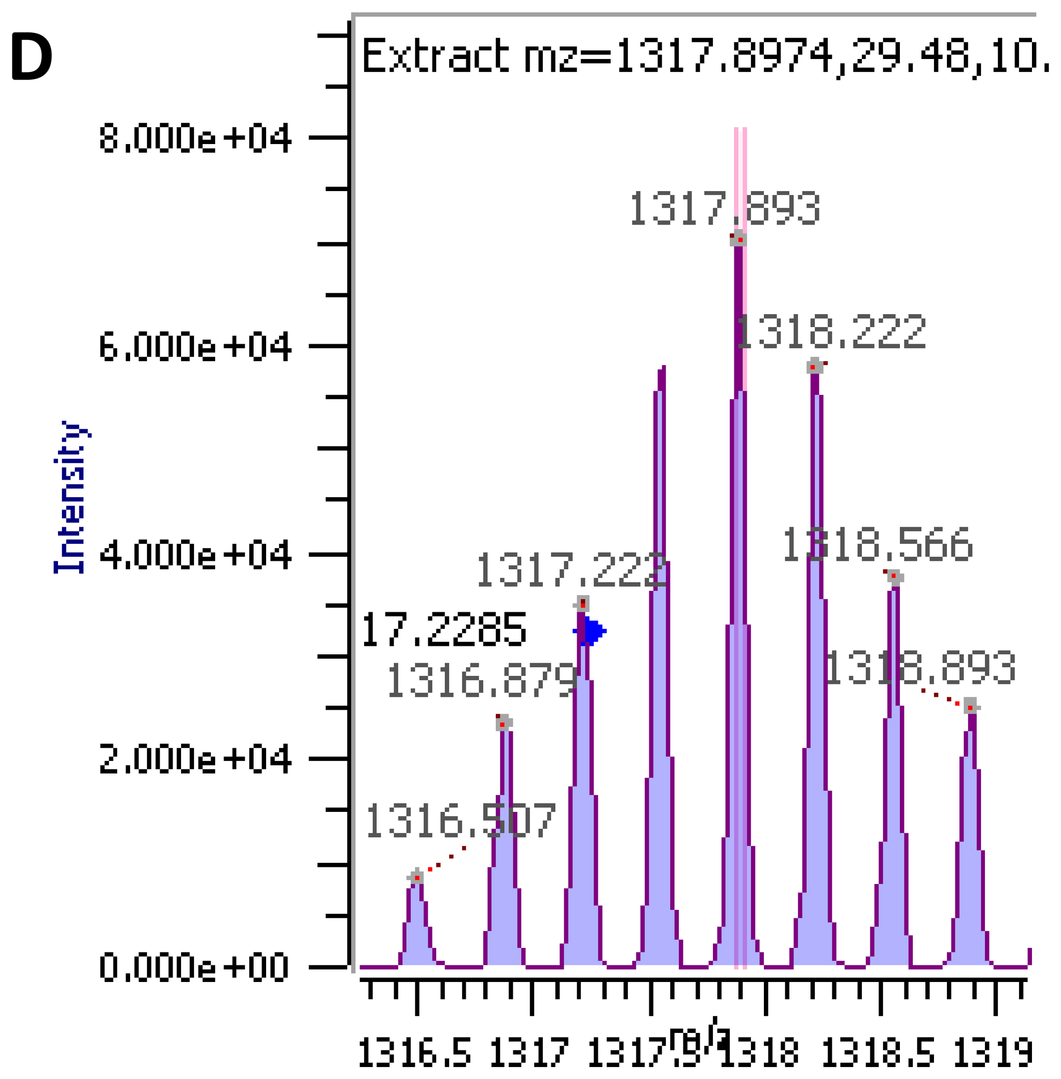


**Supplementary figure S9. Example of a gPSM with the features assigned for the validation category named as "Quasi-true Match change glycan".** (**A**) Validation category color code (**B**) Fragment ion spectrum annotated by Byonic software (**C**) Features description (**D**) Isotopic pattern.

### Quasi-true Corrected Match

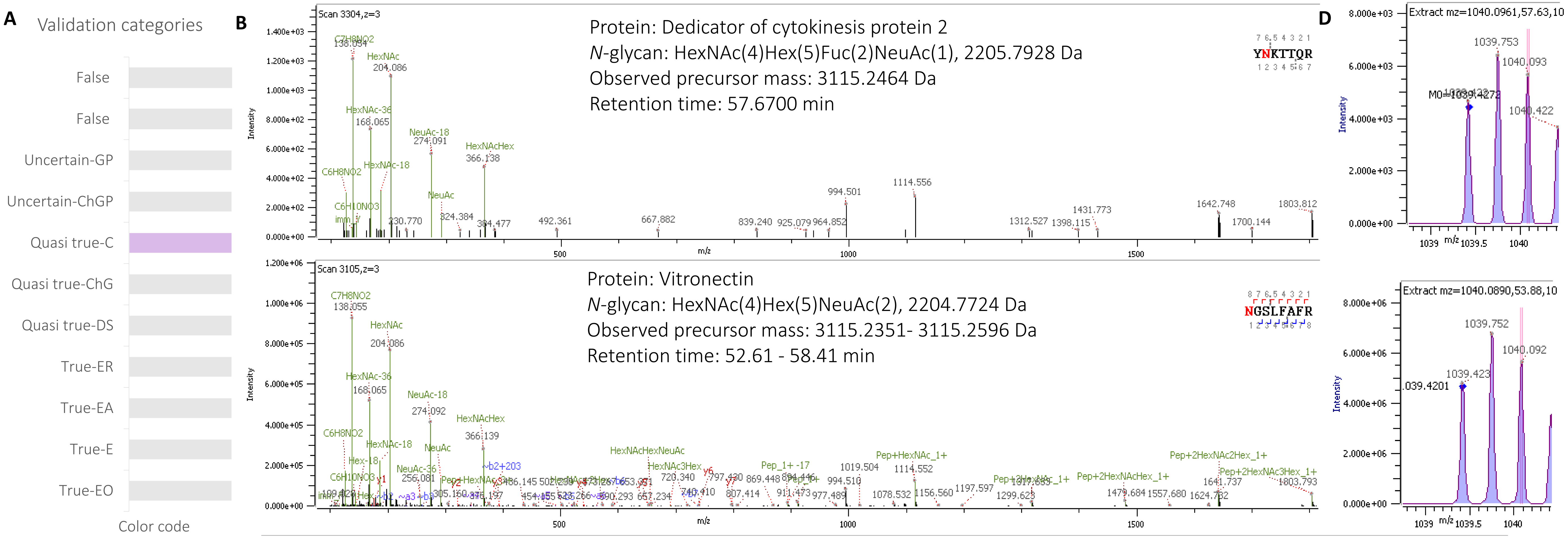


C

| | |
|---|---|
| Main validation category features | • No match with the characteristic glycopeptide fragment ions<br>• A true match from other peptide represents the observed glycopeptide fragment ions |
| Software identification | Dedicator of cytokinesis protein 2, HexNAc(4)Hex(5)Fuc(2)NeuAc(1) |
| Solution | Vitronectin, HexNAc(4)Hex(5)NeuAc(2) |

**Supplementary figure S10. Example of a gPSM with the features assigned for the validation category named as "Quasi-true corrected Match".** (**A**) Validation category color code (**B**) Fragment ion spectra annotated by Byonic software (**C**) Features description (**D**) Isotopic pattern.

### Uncertain-Change glycopeptide match

A

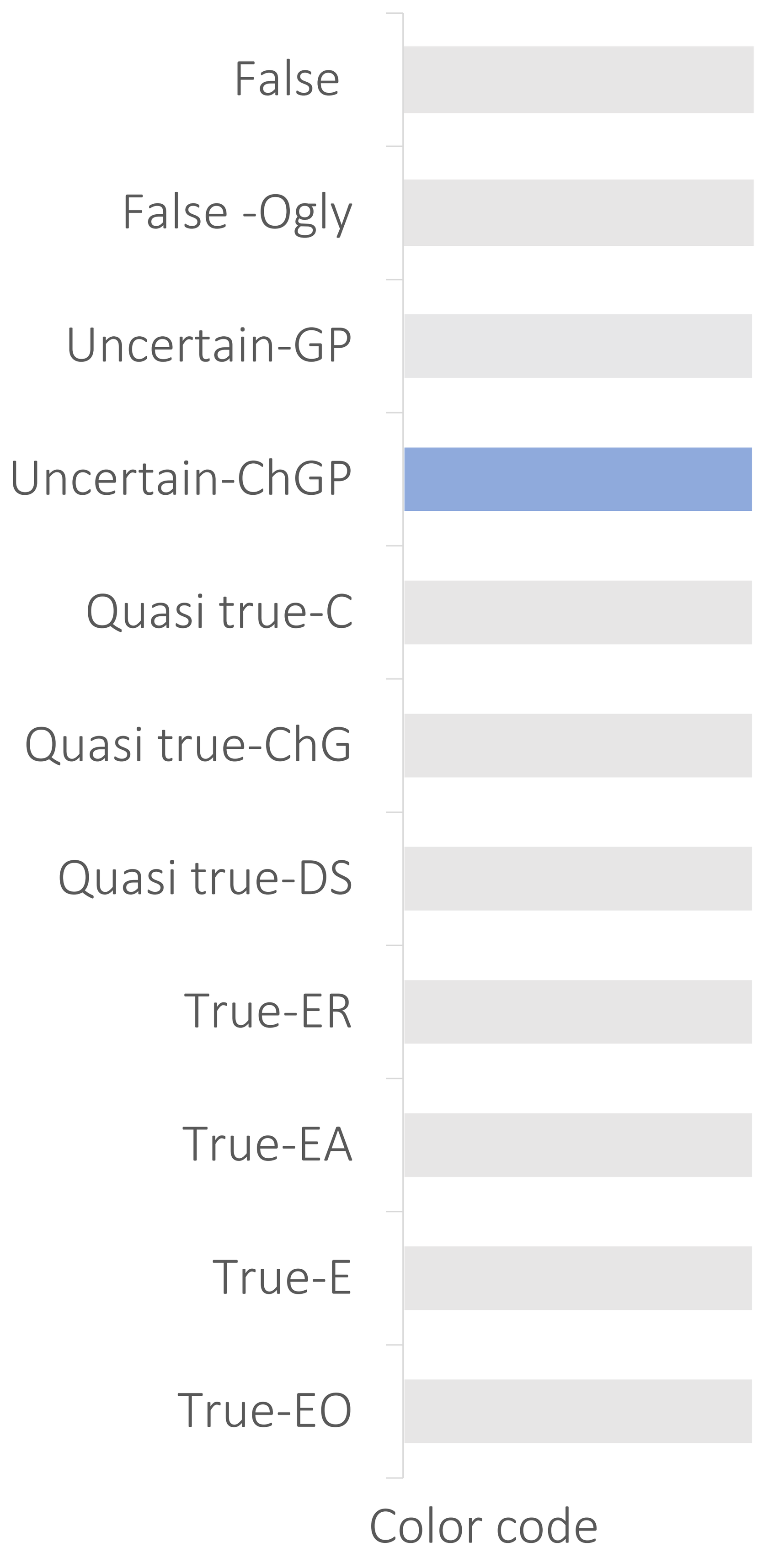


B

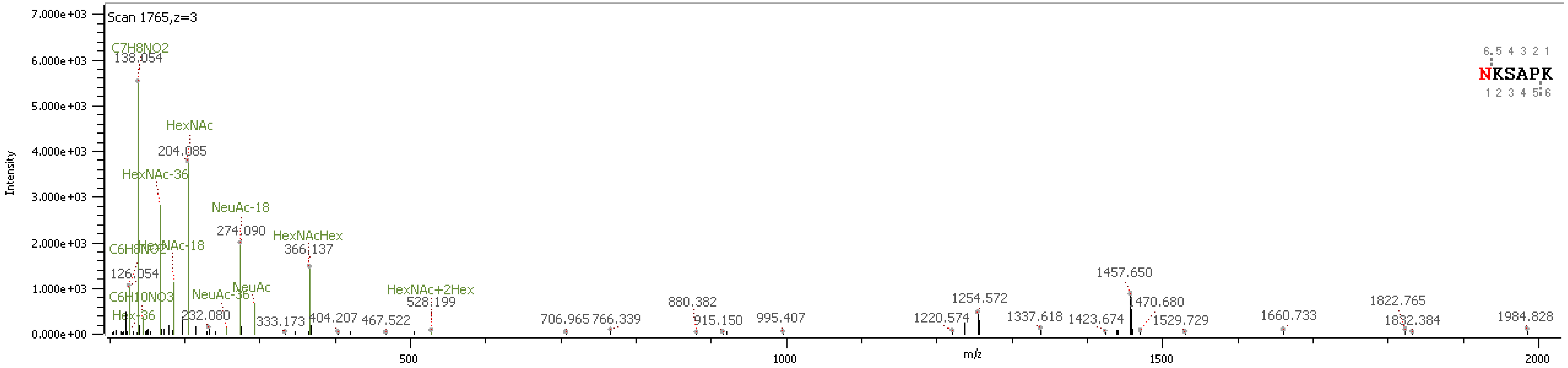


C

| | |
|---|---|
| Main validation category features | • Glycopeptide fragment ions<br>• High number of peptide fragment (b and y) ions<br>• Inconsistent evidence in regard to oxonium ions |
| Software identification | **Protein:** Xin actin-binding repeat-containing protein 2<br>***N*-glycan:** HexNAc(4)Hex(11)NeuAc(1), 2885.9940 Da |
| Expected glycan composition: | **Observed $M_{peptide}$ + H]+:** 1254.527<br>**Observed neutral mass**: 3529.3571 Da<br>**Expected glycan mass**: 2275.8371 Da |

D

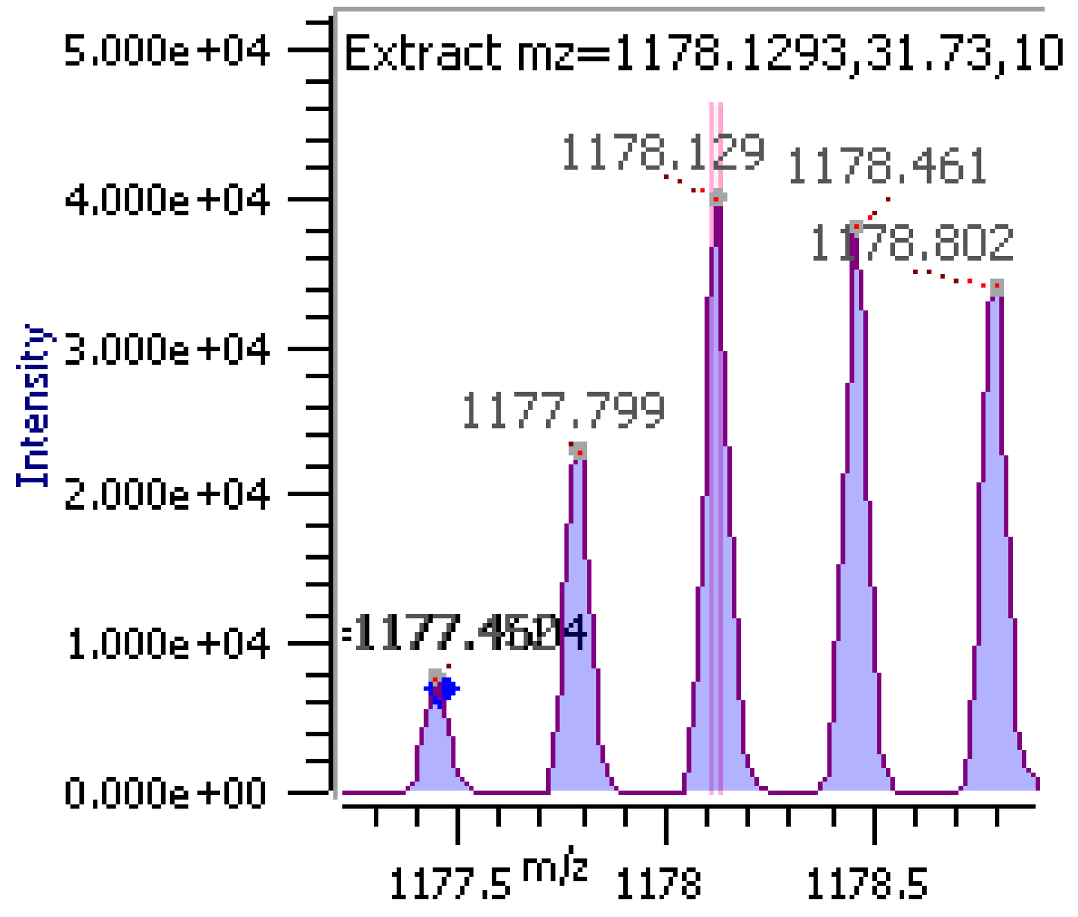


**Supplementary figure S11. Example of a gPSM with the features assigned for the validation category named as "Uncertain-Change glycopeptide match".** (**A**) Validation category color code (**B**) Fragment ion spectrum annotated by Byonic software (**C**) Features description (**D**) Isotopic pattern.

### Uncertain-glycopeptide

A

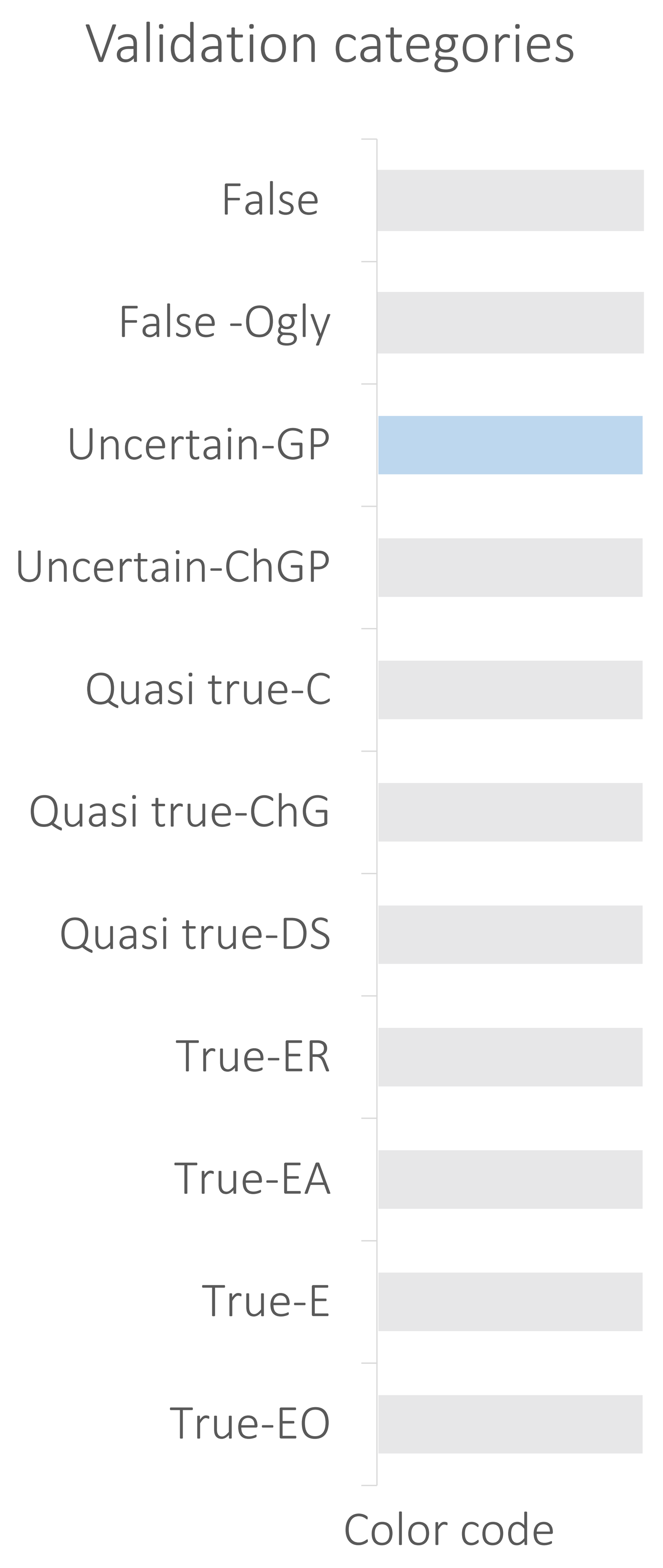


B

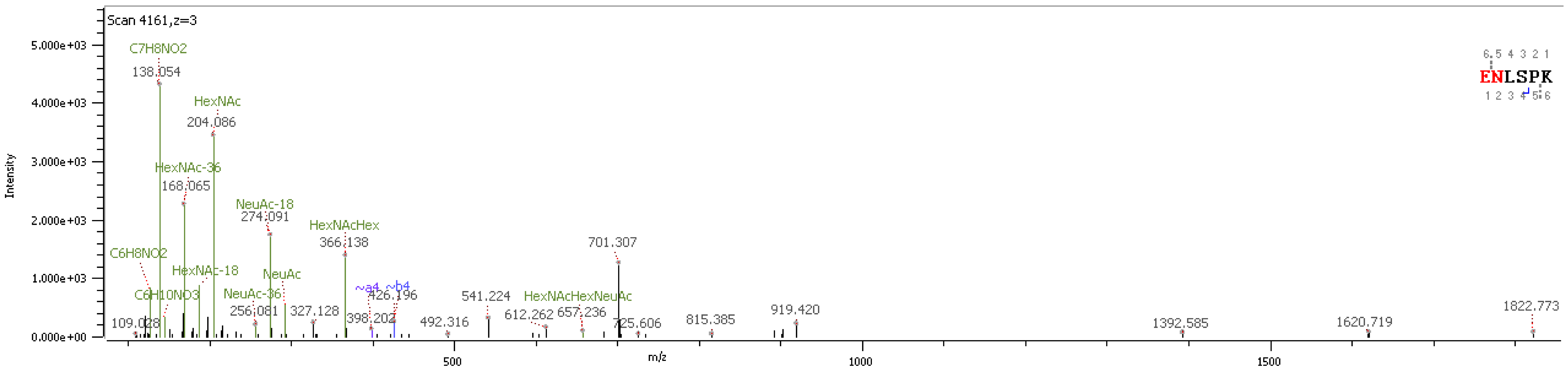


C

| | |
|---|---|
| Main validation category features | • No characteristic glycopeptide fragment ions<br>• No b and y ions |
| Software identification | **Protein:** Extended synaptotagmin-2<br>***N*-glycan:** HexNAc(7)Hex(8)Fuc(1)NeuAc(1), 3155.1315 Da<br>Peptide modification: Glu->pyro-Glu/-18.0106 |

D

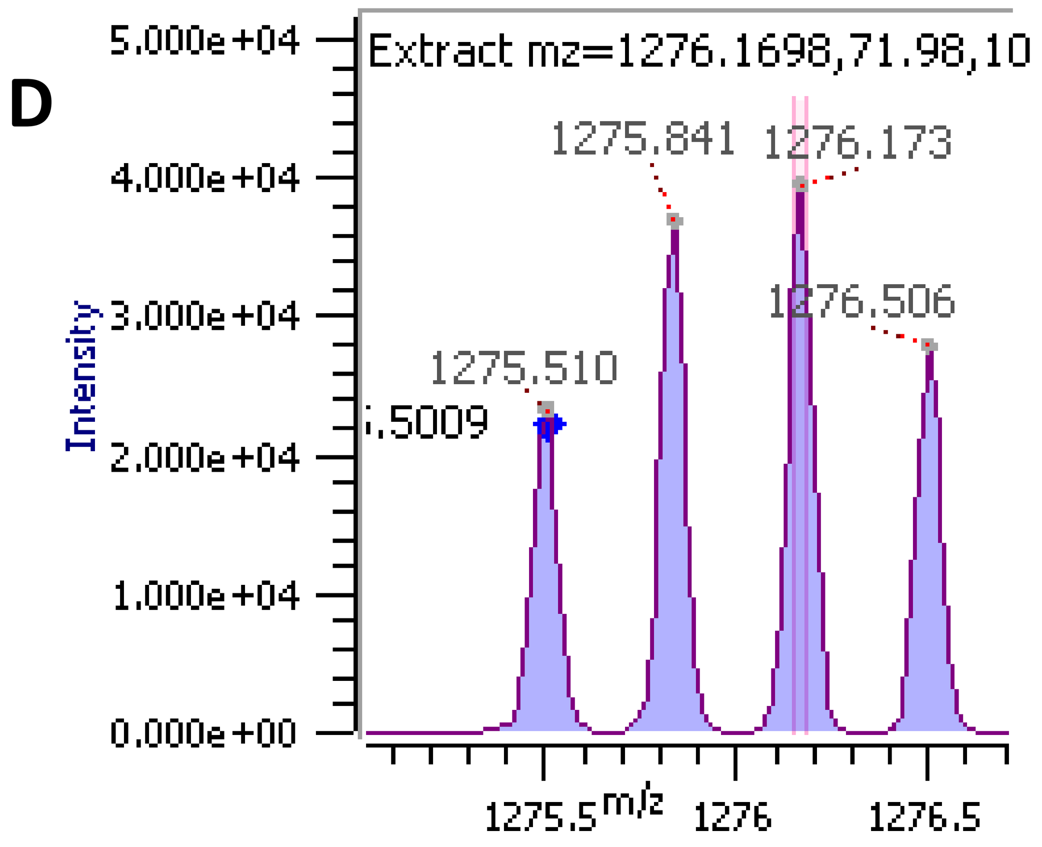


**Supplementary figure S12. Example of a gPSM with the features assigned for the validation category named as “Uncertain glycopeptide”.**
(**A**) Validation category color code (**B**) Fragment ion spectrum annotated by Byonic software (**C**) Features description (**D**) Isotopic pattern.

### False O-glycan

A

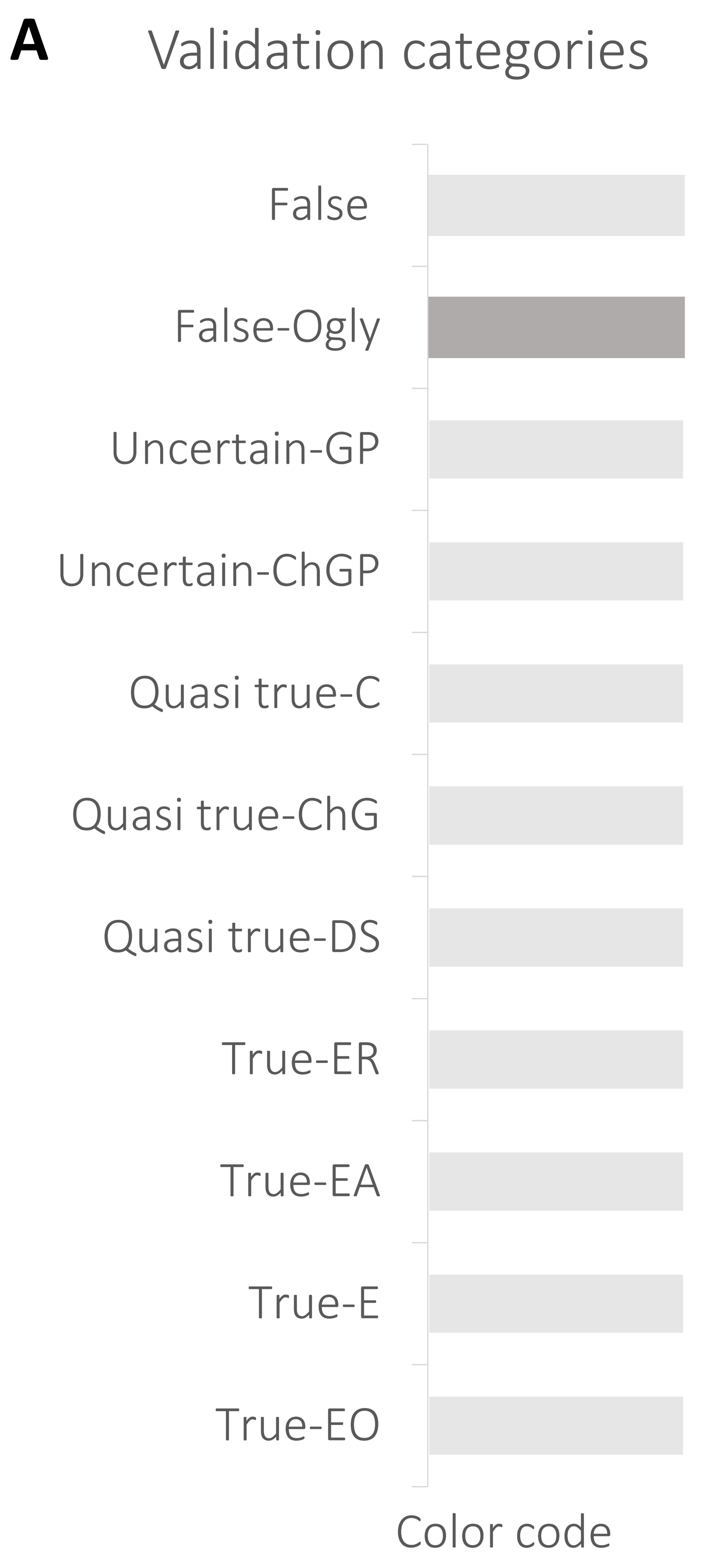


B

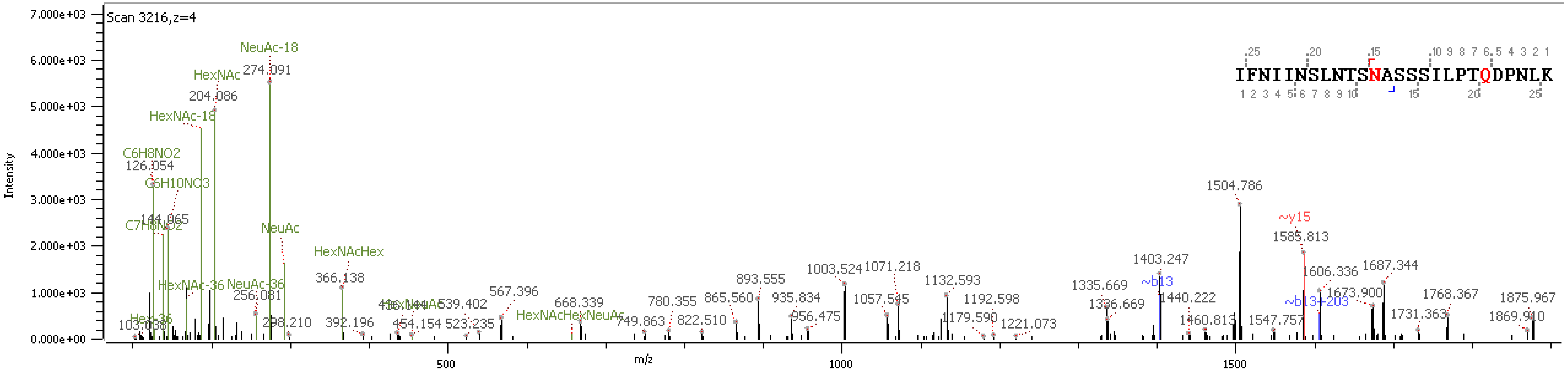


C

| Main validation category features | • Signal intensity (HCD.step) $\frac{HexNAc^{-H_2O}}{HexNAc} > 0.3$<br>• No characteristic glycopeptide fragment ions<br>• No b and y ions |
|---|---|
| Software® identification | Protein: Fer-1-like protein 5<br>*N*-glycan: HexNAc(5)Hex(5)Fuc(1)NeuAc(1) |

D

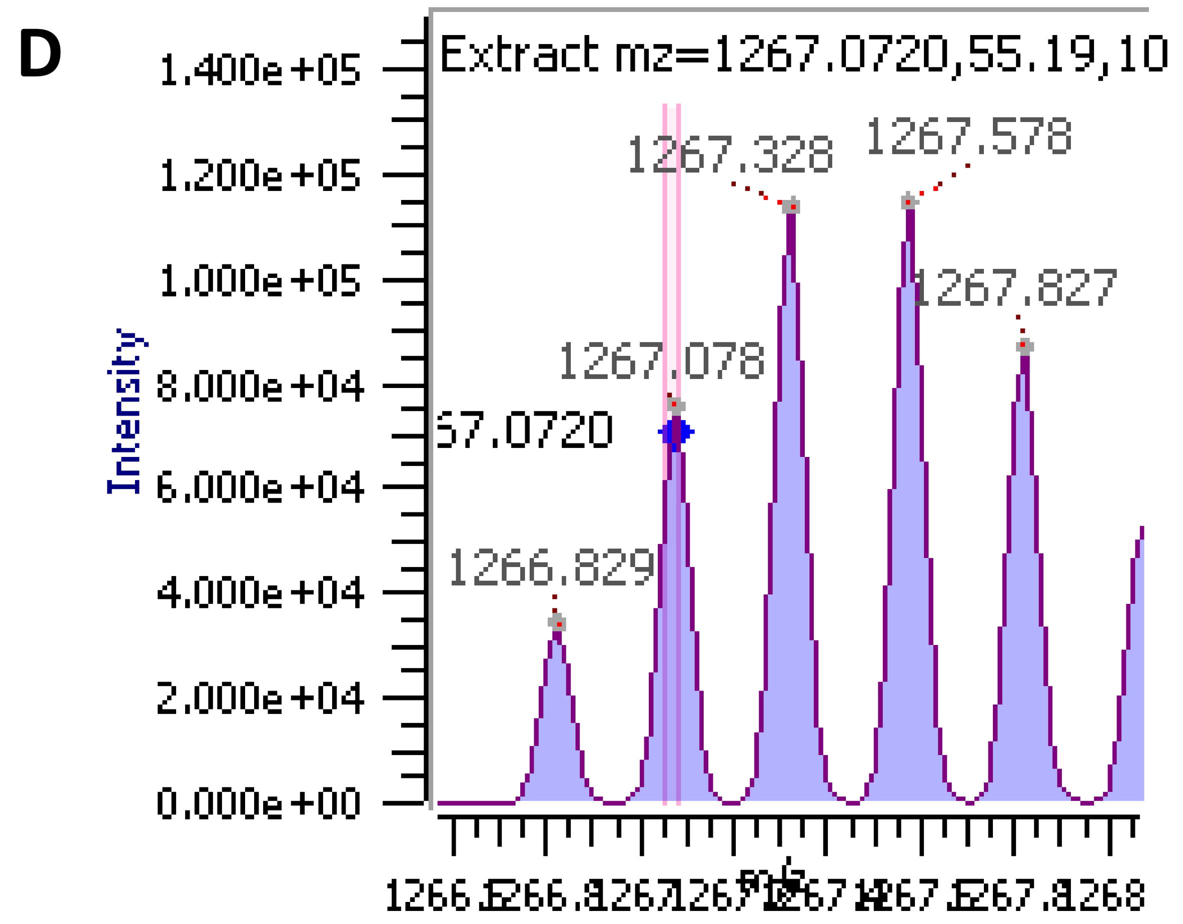


**Supplementary figure S13. Example of a gPSM with the features assigned for the validation category named as “False O-glycan”.** (**A**) Validation category color code (**B**) Fragment ion spectrum annotated by Byonic software (**C**) Features description (**D**) Isotopic pattern.

### False

**A**

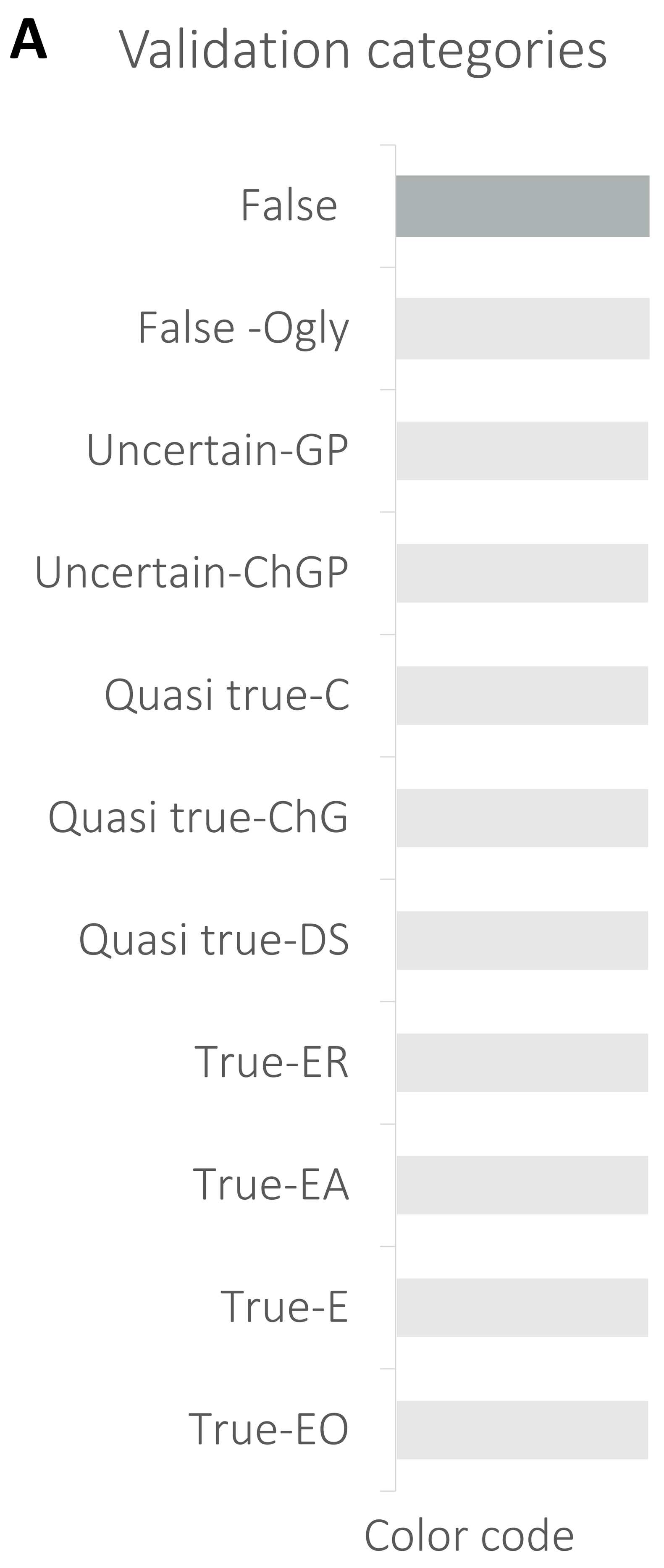


**B**

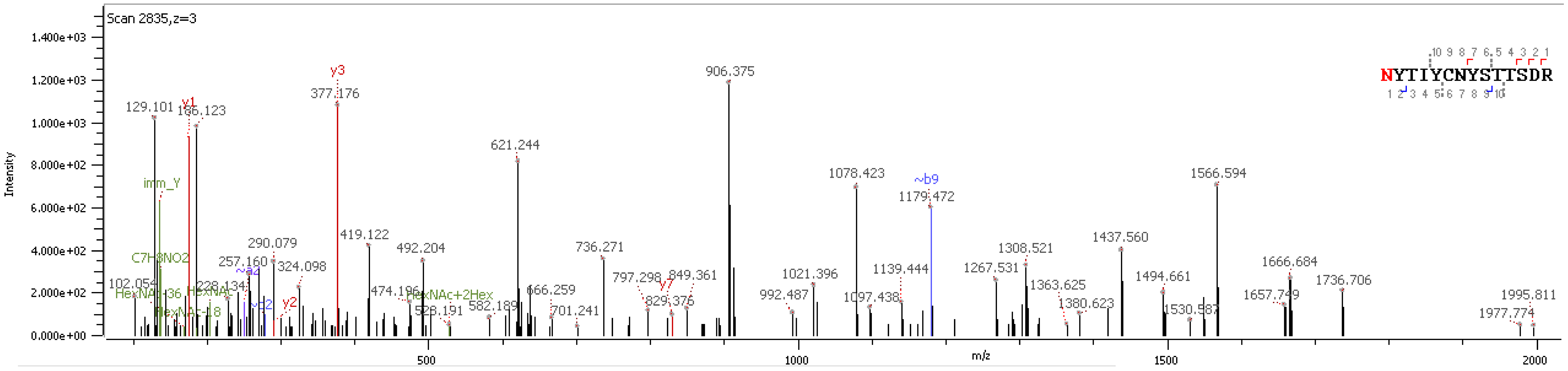


**C**

| | |
|---|---|
| Main validation category features | • No characteristic glycopeptide fragment ions<br>• No b and y ions |
| Software® identification | Protein: Extended synaptotagmin-2<br>*N*-glycan: HexNAc(7)Hex(8)Fuc(1)NeuAc(1), 3155.1315 Da<br>Peptide modification: Glu->pyro-Glu/-18.0106 |

**D**

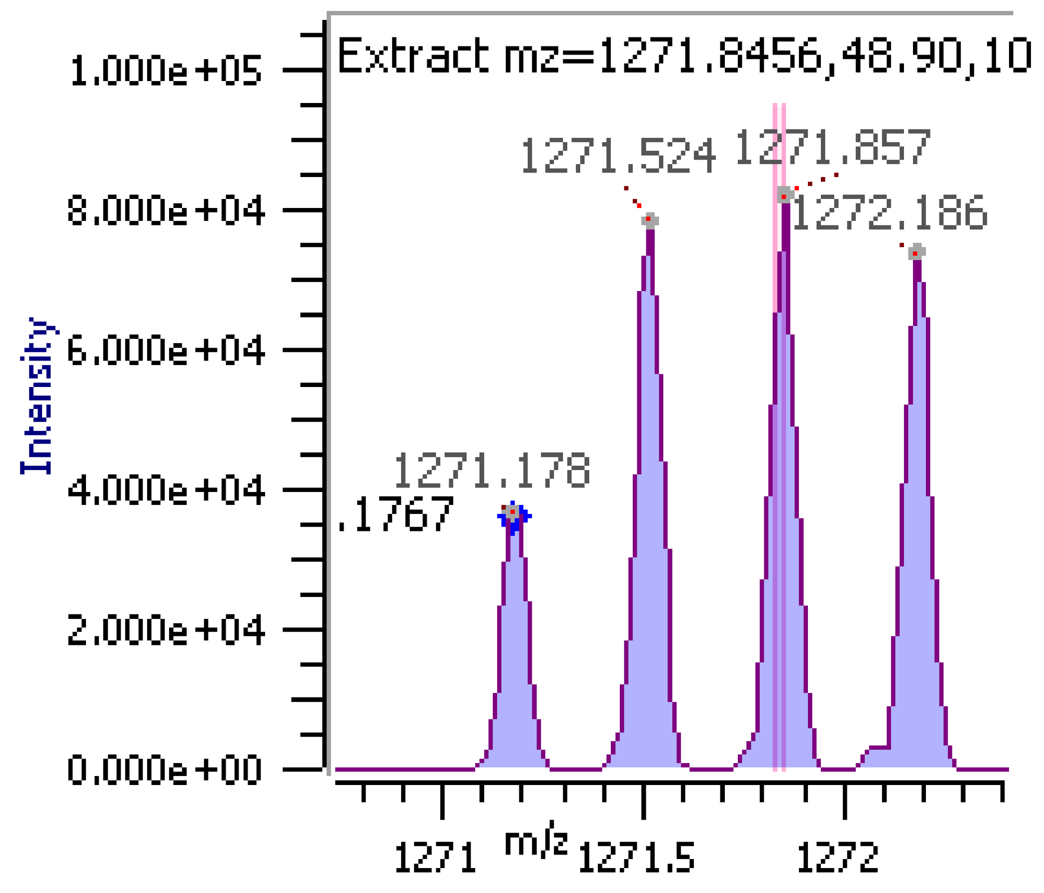


**Supplementary figure S14. Example of a gPSM with the features assigned for the validation category named as "False".** (**A**) Validation category color code (**B**) Fragment ion spectrum annotated by Byonic software (**C**) Features description (**D**) Isotopic pattern.

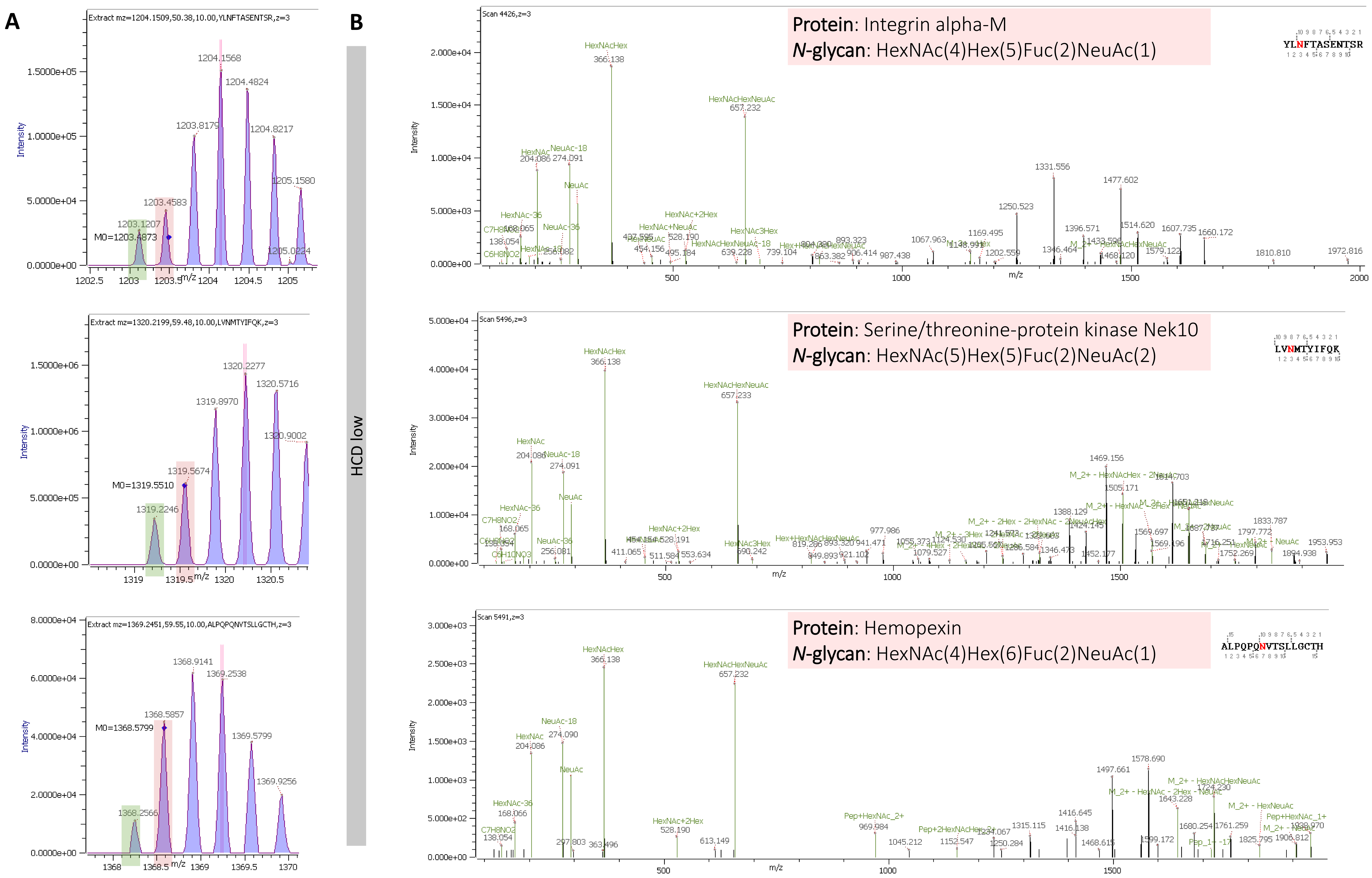


**Supplementary figure S15-1. Examples of gPSM including multiple fucose in a complex *N*-glycan with a potential error in the assignation of the monoisotopic peak.** (**A**) MS[1] Isotopic distributions. Monoisotopic peaks selected by Byonic software (red). (**B**) ) MS[2] HCD.low annotation proposed by the software assigning an incorrect *N*-glycan. Correct precursor ion highlighted in green.

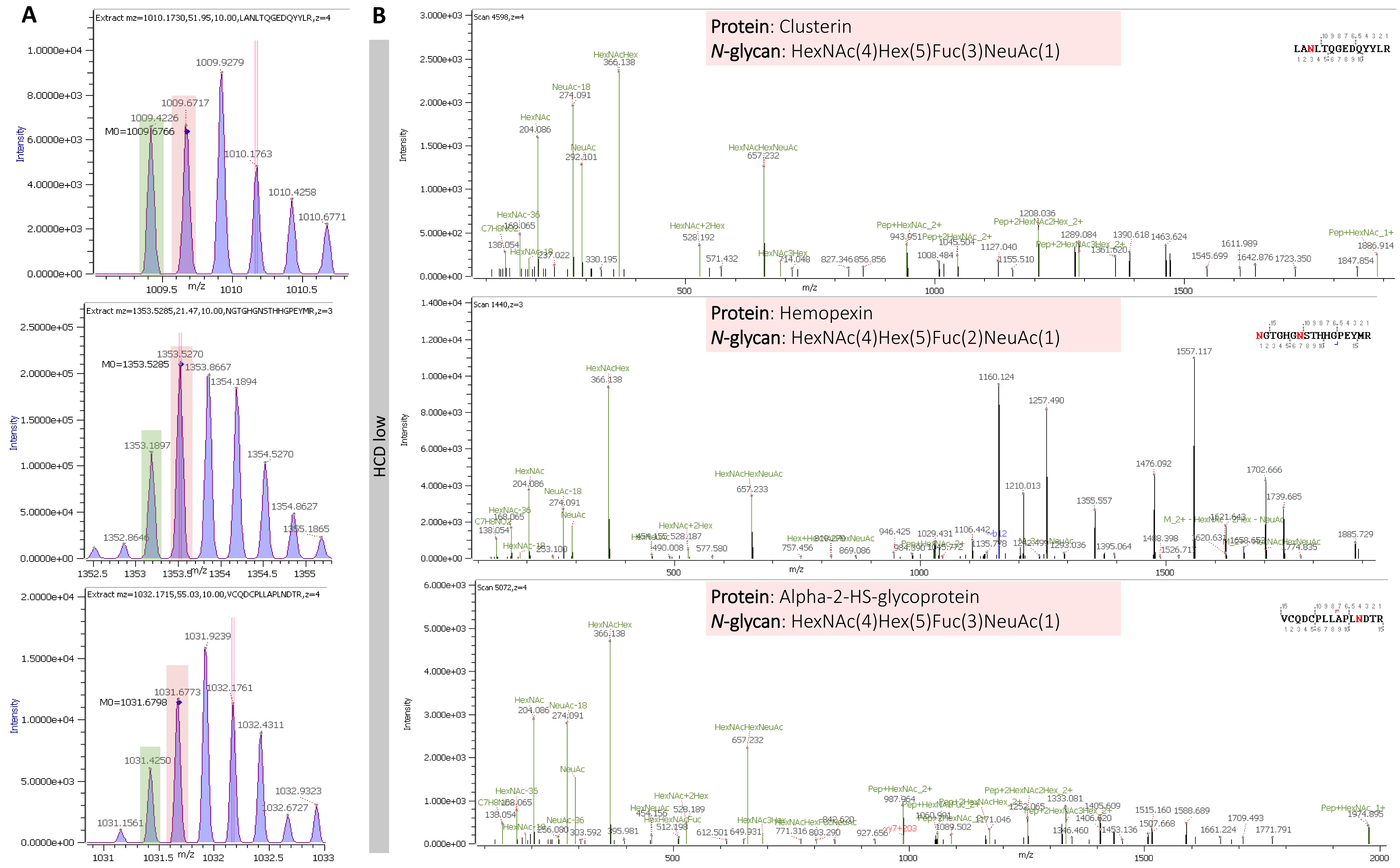


**Supplementary figure S15-2. Examples of gPSM including multiple fucose in a complex *N*-glycans with a potential error in the assignation of the monoisotopic peak.** (**A**) MS[1] Isotopic distributions. Monoisotopic peaks selected by Byonic software (red). (**B**) ) MS[2] HCD.low annotation proposed by the software assigning an incorrect *N*-glycan. Correct precursor ion highlighted in green.

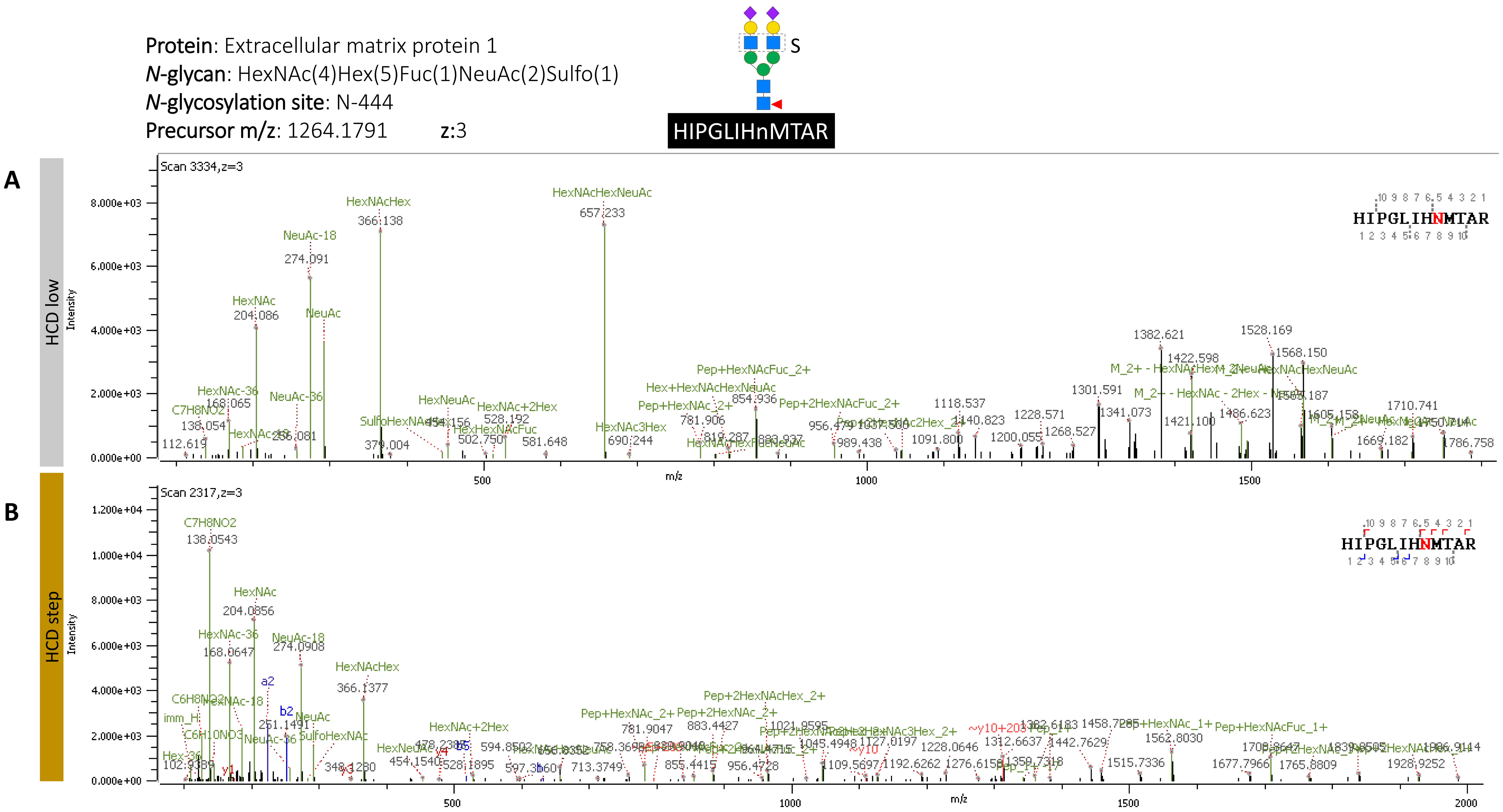


**Supplementary figure S16. Byonic software annotation of fragment ions released from sulfated N-glycopeptide.** (**A**) HCD.low fragment ion spectrum (**B**) HCD.step fragment ion spectrum.

Protein: Biotinidase
*N*-glycan: HexNAc(3)Hex(7) NeuAc(1)Phospho(1)
*N*-glycosylation site: N-150
Precursor m/z: 1079.4146 z: 3

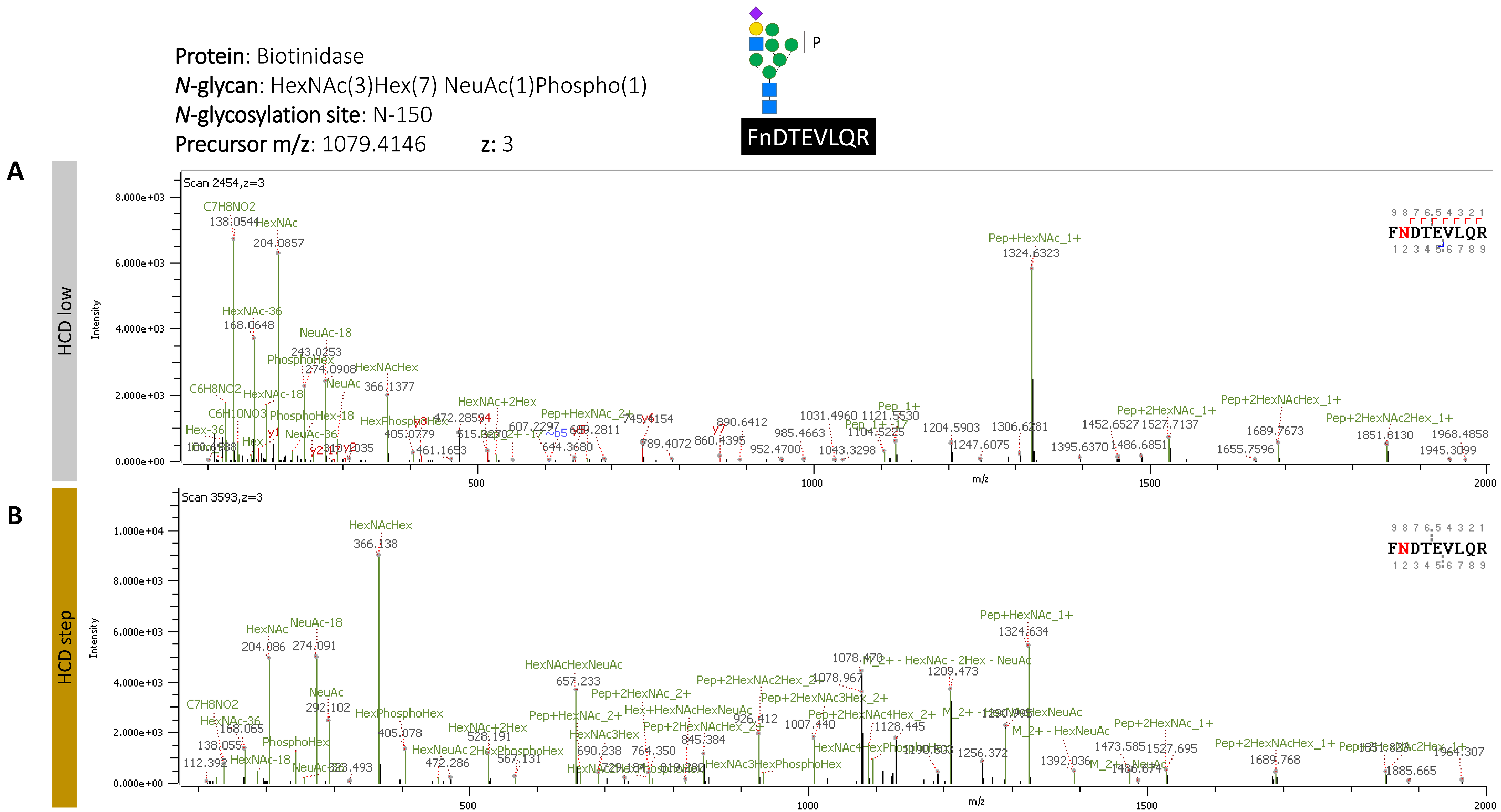


**Supplementary figure S17. Byonic software annotation of fragment ions released from phosphorylated *N*-glycopeptide.** (**A**) HCD.low fragment ion spectrum (**B**) HCD.step fragment ion spectrum.

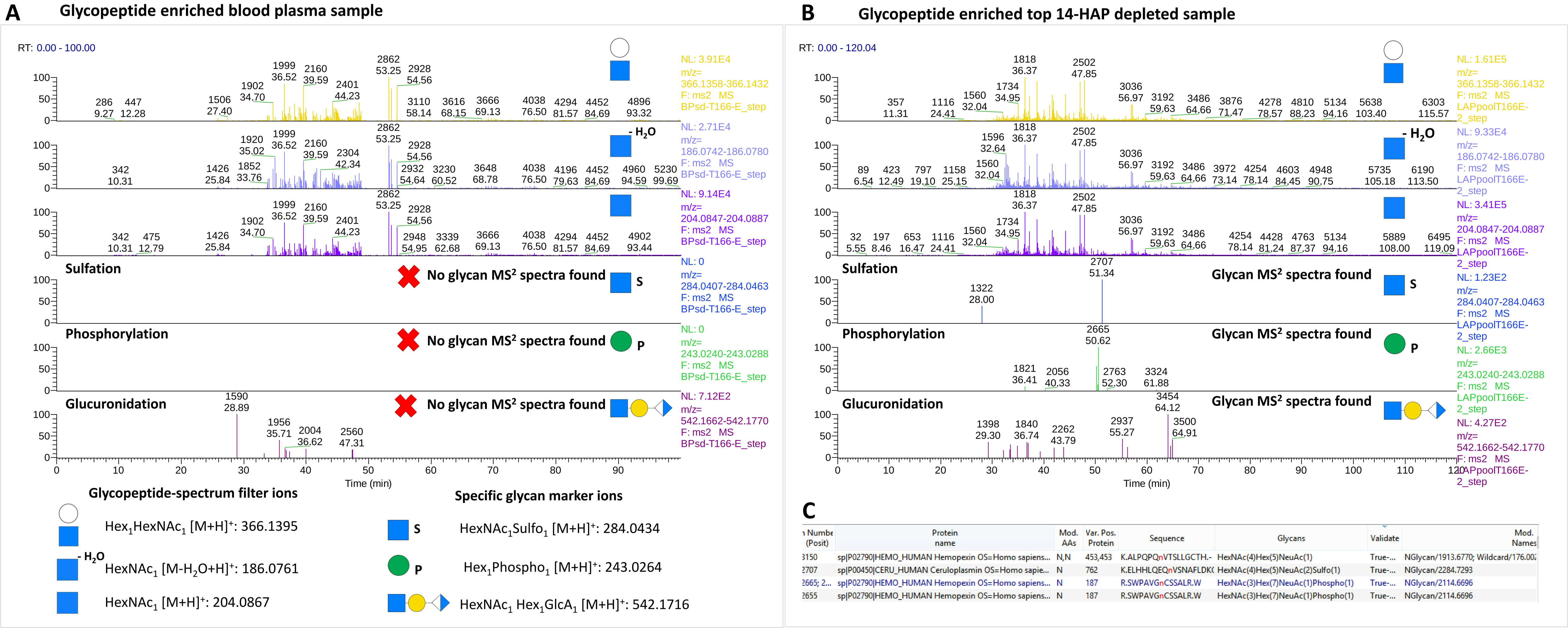


**Supplementary figure S18. Searching for rare *N*-glycopeptides in untreated blood plasma and top 14-HAP depleted sample.** (**A**) LC-MS/MS-HCD.step analysis of glycopeptide enriched blood plasma. No evidence of sulfated, phosphorylated or glucuronidated glycopeptide-derived MS$^2$ spectra. The signals observed in the glucuronidation marker ion-filter do not derive from glycopeptides. (**B**) LC-MS/MS-HCD.step analysis of glycopeptide enriched of top 14-HAP depleted blood plasma. Evidence of sulfated, phosphorylated or glucuronidated glycopeptide-derived MS$^2$ spectra. (**C**) Top 14-HAP depleted blood plasma Byonic search identified the *N*-glycopeptides holding sulfated, phosphorylated and glucuronidated *N*-glycans. The layout-filters containing glycan marker ions was created in Thermo Xcalibur Qual Browser (Version 2.2, Thermo Scientific) software.

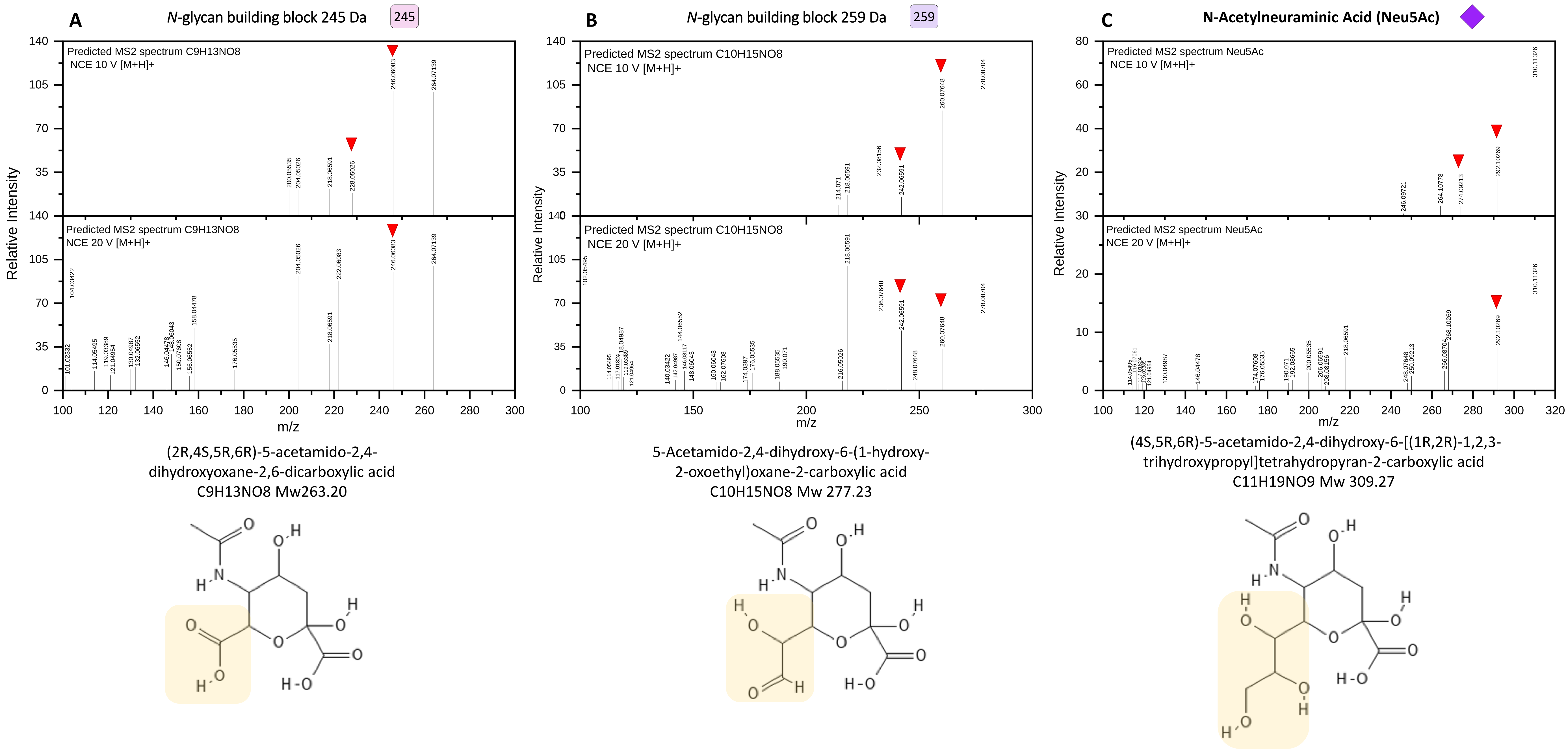


**Supplementary figure S19. ESI-MS/MS prediction for two molecules proposed as the *N*-glycan building blocks of the rare *N*-glycopeptide identifications observed.** (**A**) Predicted fragmentation of molecule with the chemical composition C9H13NO8 suggested for *N*-glycan building block 245 Da. The molecule found in PubChem (CID 59861591) is shown below the spectra. (**B**) Predicted fragmentation of molecule with the chemical composition C10H15NO8 suggested for *N*-glycan building block 259 Da. The molecule found in PubChem (CID 59173067) is shown below the spectra. (**C**) Predicted fragmentation of Neu5Ac; structure shown below the spectra (PubChemCID 439197). Red triangles indicate the fragment ions observed in the fragment ion spectra containing the rare *N*-glycans and sialic acid. Predicted spectra computed by CFM-ID (https://cfmid.wishartlab.com/predict) for two collisional energies (10 eV and 20 eV) in positive ion mode including [M+H]+ as adduct. Molecules drawn in PubChem Sketcher [47-51].

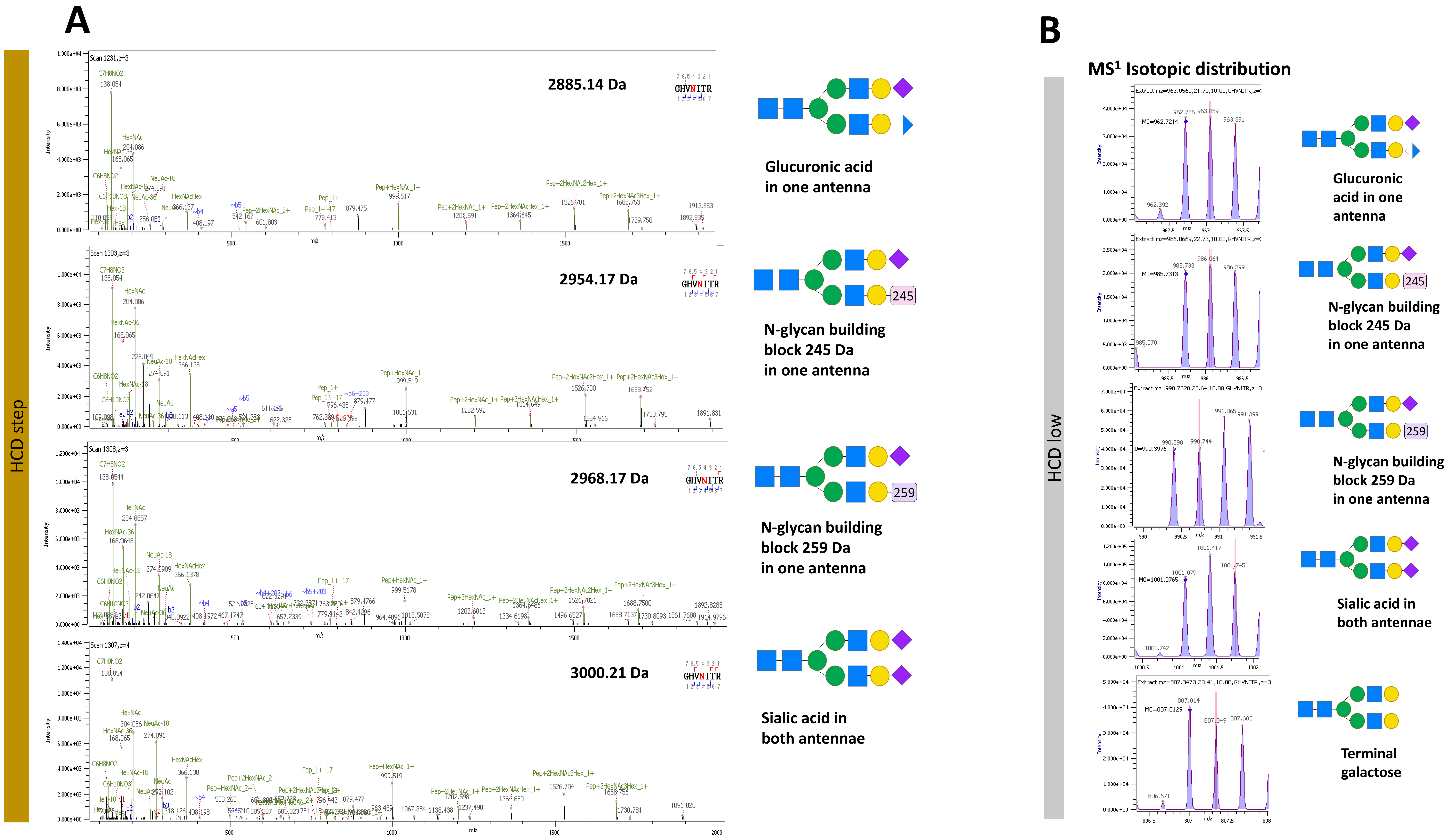


**Supplementary figure S20. Byonic software annotation of HCD.step fragment ion spectra containing rare *N*-glycans found in N121 from prothrombin (UniProt P00734).** (**A**) N-glycopeptide annotation by the software. (**B**) $MS^1$ Isotopic distribution of precursor ions observed through HCD.low acquisition.

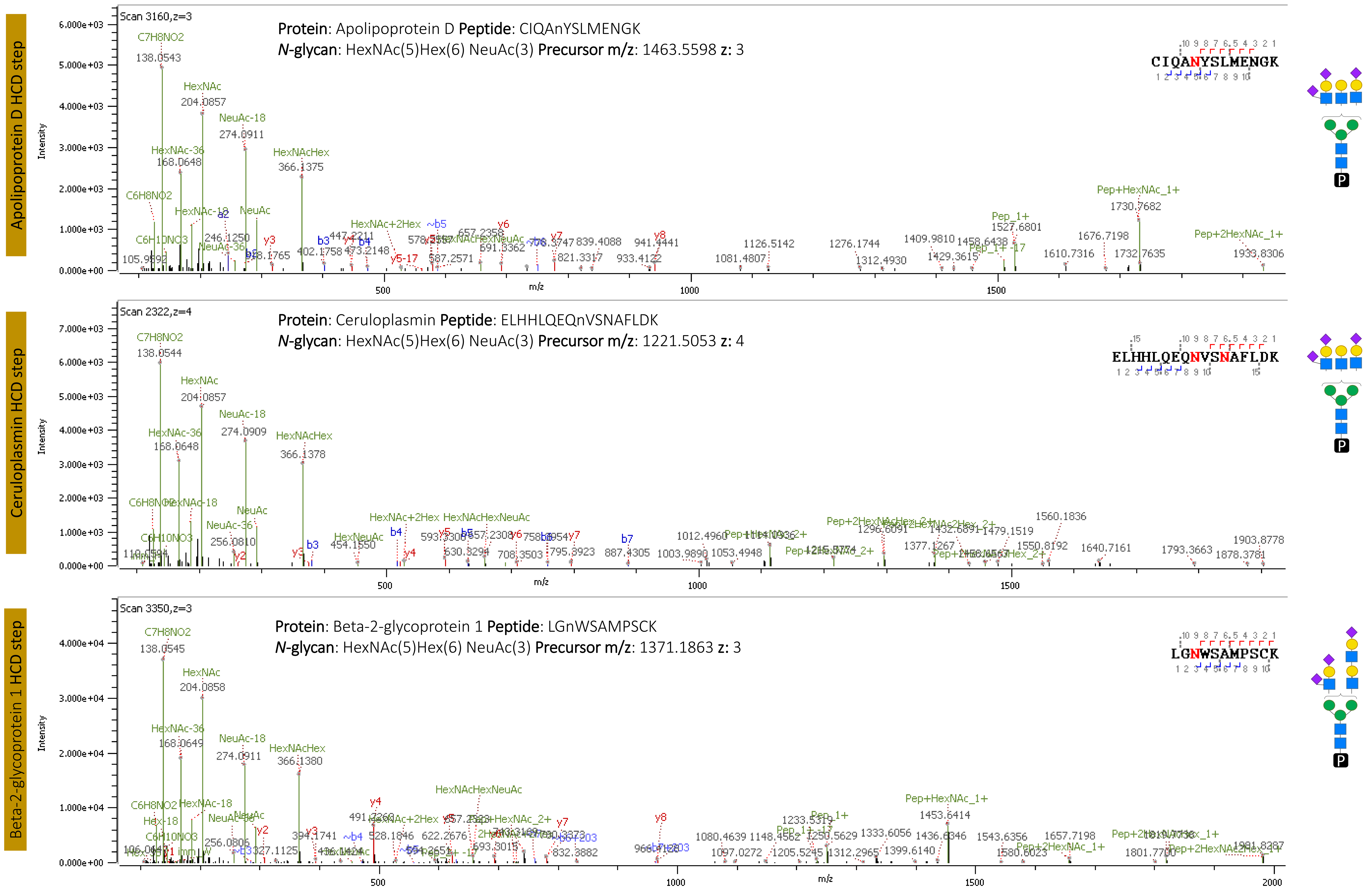


**Supplementary figure S21. Byonic software annotation of HCD.step fragment ion spectra from *N*-glycopeptides with disialo-antennary structure confirming HexNAc-NeuAc linkage.**
